# Probability-based methods for outlier detection in replicated high-throughput biological data

**DOI:** 10.1101/2020.08.07.240473

**Authors:** Matthew S. Smith, Karthik Devarajan

**Affiliations:** Program in Biophysics, University of California, San Francisco; Department of Biostatistics & Bioinformatics, Fox Chase Cancer Center, Temple University Health System, Philadelphia

**Keywords:** technical reproducibility, replicate, outlier, asymmetric Laplace, generalized gamma, coefficient of variation, high-throughput biology, anomaly detection

## Abstract

A problem that arises frequently in high-throughput biological studies is the assessment of technical reproducibility of data obtained under homogeneous experimental conditions. This is an important problem considering the significant growth in the number of high-throughput technologies that have become available to the researcher in the past decade. Although certain ad hoc and graphical methods for determining data quality have been in existence, these methods lack statistical rigor and are not broadly applicable across different technologies. There is an inherent need for the quantitative evaluation of the reproducibility of technical replicates from high-throughput compound and siRNA screening, next-generation sequencing and other modern “omics” studies. To this end, we have developed an approach that accounts for technical variability and potential asymmetry that arise naturally in the distribution of replicate data, and aids in the identification of outliers. Our methods for outlier detection rely on flexible statistical models, employ maximum likelihood methods for estimation and are broadly applicable to a variety of high-throughput biological studies. We discuss an adaptation of these methods when there are multiple replicates as well as current limitations to these techniques. We illustrate our methods using experimental data from high-throughput compound screening and protein expression studies as well as simulated data. Our methods are implemented in the R package replicateOutliers and are currently available at github.com/matthew-seth-smith/replicateOutliers.

## 1 Introduction

Advances in high-throughput technologies in the past two decades have given rise to large-scale biological data that is measured on a variety of scales. For example, high-throughput compound and siRNA screening assays are specifically designed to detect interactions with compounds by directly measuring inhibition of siRNA or kinase activity. Gene expression studies, on the other hand, enable the simultaneous measurement of the expression profiles of tens of thousands of genes and proteins, often from a relatively small number of biological samples. Traditionally this has involved the use of microarray technology to measure mRNA expression, and more recently, the use of SNP arrays to measure allele-specific expression and DNA copy number variation, methylation arrays to quantify DNA methylation and next-generation sequencing technologies such as RNA-Seq, ChIP-Seq etc. for the measurement of digital gene expression.

These studies have resulted in massive amounts of data requiring analysis and interpretation while offering tremendous potential for growth in our understanding of the pathophysiology of many diseases. An important problem that often arises in these studies is the assessment of technical reproducibility of replicate data obtained under homogeneous experimental conditions. Methods for determining the quality of data obtained from microarrays have been in existence for many years; however these methods are not readily applicable to data obtained from other technologies. Moreover, these methods tend to rely primarily on visualization and do not employ rigorous statistical methods. In this paper, we demonstrate that replicate data from high-throughput biological studies exhibit asymmetry in their distribution due to the data generating mechanism and provide an interpretation for it. We then describe a model-based approach that accounts for inherent noise in replicate data and aids in the identification of outlying observations. This data driven, probability-based, approach borrows strength from the large volume of data available in these studies and can be used for assessing technical reproducibility independent of the technology used to generate the data. We illustrate our methods using case studies from high-throughput compound screening and protein expression.

Previous methods of outlier detection (also called anomaly detection in machine learning) tend to focus on supervised or unsupervised learning (Chandola et al., 2009). We have data to which we would like to fit a regression model or decision tree, and we wish to remove the points that lead to incorrect models. We can plot the residuals from the model or use more precise methods like Cook’s distance (Cook, 1977). Once we find the anomalies, we can remove them from the training data and re-fit the model. These methods work in contexts where we have single measurements of our covariates, and we want to see which ones are outliers when fitting a model. In our new context, we will be working with replicated measurements of a large number of variables and looking for observations that are significantly different from each other. The proposed methods are useful for testing whether a new instrument, high-throughput biological assay, or compound screening experiment is reproducible.

Let *X*_1_ and *X*_2_ denote duplicate, non-negative measurements from independent experiments obtained for each of *p* variables in a high-throughput study. For example, the *p* variables could represent genes, probes or sequence tags in gene or protein expression studies (such as microarrays, RNA-Seq or mass spectrometry) or kinase-inhibitor pairs in a compound screening study; and *X*_1_ and *X*_2_ represent measurements of gene expression, protein expression or compound activity. In these studies, data is measured naturally on the non-negative scale and can be used directly for our purpose without the need for transformations. Two quantities are of interest in assessing the reproducibility of technical replicates – the difference, Δ = *X*_1_ − *X*_2_, and variability in duplicate measurements. In order to account for the potential signal-dependence of variability in assay measurements, the coefficient of variation (*Z*) is used to estimate variation rather than standard deviation (or variance). *Z* is the standard deviation of duplicate measurements normalized to their mean, and in this case, reduces to

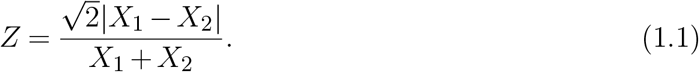

A scatter plot of *Z* versus Δ would provide a visual display of variability in duplicates as a function of their difference. This approach is somewhat similar in principle to an MA plot commonly used to assess data quality in gene expression microarrays where *M* and *A* denote, respectively, the difference and mean of replicates on the logarithmic scale. The MA plot itself owes its origins to the difference versus mean plot, proposed by Altman and Bland (1983), to visually assess the mean-variance relationship in replicate data. However important differences exist between this and our proposed approach, as will be shown in subsequent sections. In order to harness the distributional properties of Δ and *Z*, the original scale of measurements is used in all computations. The use of *Z* is a robust alternative to estimate the variability of replicates that automatically incorporates its mean dependence, and plotting it against Δ provides an approach for identifying outlying observations while accounting for inherent noise in assay measurements.

It is easy to show that for *X*_1_, *X*_2_ ∈ ℝ^+^, we have Δ ∈ ℝ and 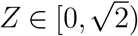. By using both Δ and *Z*, we have measures of absolute and relative differences between measurements. The method presented by Anastassiadis et al. (2011) provides a framework for duplicated data *X*_1_ and *X*_2_. For measurements of inhibition for different pairs of kinases and commercially-available inhibitors presented in the aforementioned paper, the transformation from (*X*_1_, *X*_2_)-data to (Δ, *Z*)-data produces a plot resembling a “volcano plot” commonly seen in graphs of fold-change (also known as biological significance) against statistical significance in genomic studies (see Supplemental Information (SI), Supplementary Figures 1 & 2).

While we do not know the true data generating models for *X*_1_ and *X*_2_, we can fit different probability density functions (PDF) with the proper support to the data points. If we are dealing with non-negative data, we may start by assuming that the data are generated by independent exponential models, where *X*_*i*_ ∼ *Exp*(*λ*_*i*_), *λ*_*i*_ > 0 has the PDF

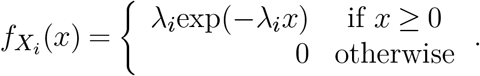

Exponential models arise naturally in modeling biological data. If an underlying hidden variable takes values greater than 1, then we may model it as a Pareto distribution, and its logarithm (which we can measure) will have an exponential distribution (Rice, 2007). If we wish to measure the difference between these log-transformed variables, we should expect our data to have an asymmetric Laplace distribution (see SI, §1, for more details). Purdom and Holmes (2005) used this reasoning to demonstrate that two-color microarray data, expressed as the log-ratio of red and green channel intensities, exhibits the asymmetric Laplace distribution, meaning we can theoretically model the log-intensities of the red and green channels as exponential distributions. Other applications of Pareto distributions and their generalizations for high-throughput biological data include modeling mRNA expression in Serial Analysis of Gene Expression (SAGE) studies and modeling the distribution of PM (perfect match) expression intensities in oligonucleotide Affymetrix microarray data (Kuznetsov, 2001; Wu et al. 2003). After further adjustments, Robinson and Smyth (2007) showed that data obtained in SAGE studies is similar in structure to digital gene expression data obtained from next-generation sequencing technologies such as RNA-Seq and ChIP-Seq. Marioni et al. (2008) compared the technical reproducibility of RNA-seq with that of gene expression arrays. Assessing reproducibility also has many applications in compound and siRNA screening studies as evidenced in the works of Anastassiadis et al. (2011, 2013) and Duong-Ly et al. (2016).

We first describe the compound screening data and methods presented in Anastassiadis et al. (2011, 2013) and expand it to more generalizable probability-based methods. These methods work explicitly for measurements done in duplicate, but when there are more than two replicates, we can consider every pair-wise comparison. The most conservative comparison of the pairwise method would then consider a set of replicates to be an outlier if any of the pairs would count as an outlier (we could alternatively make a different arbitrary choice). We then illustrate the utility of our new methods using this data as well as protein expression data from another study, both generated at Fox Chase Cancer Center (FCCC).

## 2 Fitting statistical models to Δ and *Z*

Using data from duplicated experiments measuring the effect of 178 commercially available kinase inhibitors against a panel of 300 recombinant protein kinases, Anastassiadis et al. (2011) fit the difference Δ in kinase activities to a symmetric Laplace distribution. Later, Anastassiadis et al. (2013) extended this approach to the asymmetric Laplace distribution using 181 kinase inhibitors against a panel of 292 kinases that significantly overlapped with the previous set. Formally, for parameters *σ* > 0, *κ* > 0, and *θ* ∈ ℝ, if 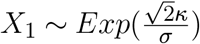 and 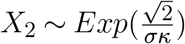 are independent exponential random variables, then Δ = *θ* + *X*_1_ − *X*_2_ has an asymmetric Laplace distribution (Kotz et al., 2001; Johnson and Kotz, 1970^*b*^). We denote this as 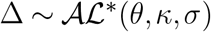, with CDF

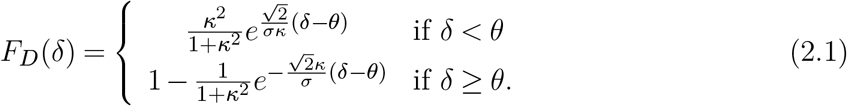

The star notation indicates this choice of parametrization. The expected value of Δ is *E*[Δ] = *μ* + *θ*, where 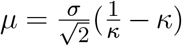, and its variance is *V ar*(Δ) = *σ*^2^ + *μ*^2^. It is important to note that the variance depends on the mean unless *μ* = 0 (or, equivalently, *κ* = 1) which results in the symmetric Laplace distribution (Kotz et al., 2001). An interesting property of the Laplace family is that the tails are heavier relative to the Gaussian family which makes it particularly useful for modeling data with a higher probability of extreme values. Supplementary Figure 3 illustrates the shapes of members of this family based on different choices of parameters.

Anastassiadis et al. (2011, 2013) used the `vglm` function in the `VGAM` package in `R` to compute maximum likelihood estimates (MLE) 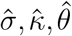 (and hence 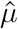) by fitting a symmetric or asymmetric Laplace distribution to Δ using the data (Yee, 2018; R Core Team, 2018). In addition, we could also consider 95% confidence intervals for log *κ* and *θ*, setting either to the null value of 0 if the confidence interval contains 0. Using the MLEs, we consider all of the data pairs for which Δ is within *k* = 1 or *k* = 2 standard deviations away from 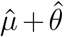 (the mean of the distribution, not the mode) to not be outliers. We remove these non-outliers (as well as the values with the top *t* = 1% and bottom *b* = 1% of *Z*-values) and then fit either a log-normal or Weibull distribution to the remaining points’ *Z* values, with the `R` package `extremevalues` (van der Loo, 2010^*a*^,^*b*^). This package finds MLEs for the model parameters. We determine the value above which we would not expect more than *N*_*u*_ = 1 observations (and below which we would not expect more than *N*_*l*_ = 1 observations) for *Z* given our remaining pairs of data points. Any *Z*-values above this level (including the top *t* and bottom *b*, if necessary) are considered outliers in our data.

The above approach does not assume an underlying data generating model and benefits from readily-available `R` code. The log-normal and Weibull distributions give us flexibility for fitting non-negative distributions to *Z*, and Kundu and Manglick (2004) created a guide for which one to use for different data sets. This method, however, does not reflect the support of the random variable 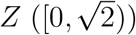 given positive random variables *X*_1_*, X*_2_ and, thus, motivates the development of probability-based methods that depend on the joint distribution of (Δ, *Z*) or marginal distributions of Δ and *Z*.

## 3 Methods based on the exponential model

As alluded to earlier, we may not know that the data is exponentially distributed, only that it is non-negative. We would make this assumption if we have reason to believe that it might be correct or simply for the computational ease of calculating the joint and marginal PDFs of Δ and *Z* (Rice, 2007; Kotz et al., 2001; Purdom & Holmes, 2005; Kuznetsov, 2001; Wu et al., 2003).

The method in Anastassiadis et al. (2013) assumed Δ had an asymmetric Laplace distribution, meaning *X*_1_ and *X*_2_ could be modeled as independent exponential distributions. We did not use this information about the distributions of *X*_1_ and *X*_2_ for finding the distribution of *Z*, meaning we also lose the dependence between Δ and *Z*. Our new approach, then, would be to start with the assumption that *X*_1_ ∼ *Exp*(*λ*_1_) and *X*_2_ ∼ *Exp*(*λ*_2_) are independent exponential random variables with *λ*_1_ *>* 0 and *λ*_2_ > 0. For simplicity, define 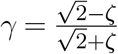, and hence 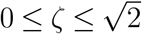 implies 0 ≤ *γ* ≤ 1. If *λ*_1_ ≠ *λ*_2_, which we will call the asymmetric case, we have the following joint cumulative density function for (Δ, *Z*):

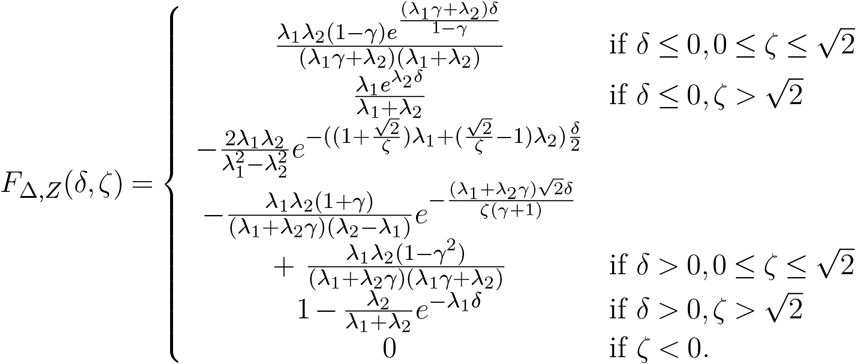

The joint CDF (Δ, *Z*) for the symmetric case, i.e, when *λ*_1_ = *λ*_2_ = *λ*, is shown in SI, §2.1.2. The forms of the symmetric and asymmetric CDFs only differ in the region where *δ* > 0 and 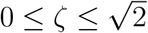. We can find the marginal distributions of Δ and *Z* from these CDFs, using *F*_Δ_(*δ*) = lim_*ζ*→∞_ *F*_Δ,*Z*_ (*δ, ζ*) and *F*_*Z*_ (*ζ*) = lim_*δ*→∞_ *F*_Δ,*Z*_ (*δ, ζ*). For the asymmetric case,

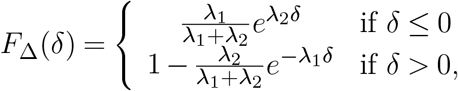

which means 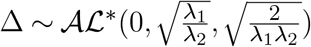 marginally. For the symmetric case,

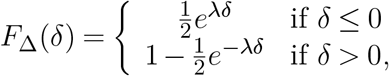

which means 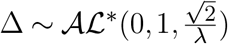, marginally, which we can also write as 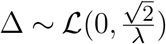 for a symmetric Laplace distribution (Kotz et al., 2001).

The marginal distribution for *Z* is not familiar. For the asymmetric case, *Z* has the CDF

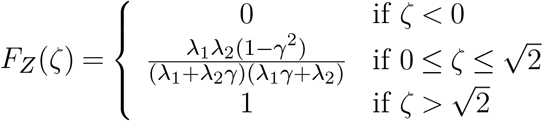

and the PDF

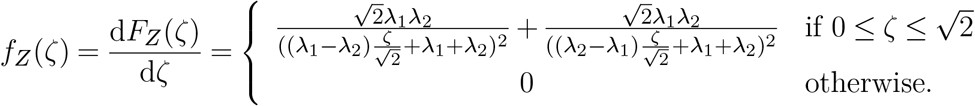

For the symmetric case, *Z* has the CDF

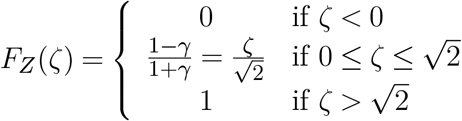

and the PDF

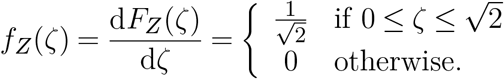

The forms of the marginal CDFs and PDFs agree between the two cases because the only differences between their joint CDFs above disappear as either *ζ* → ∞ or *δ* → ∞. We have also shown that when *X*_1_, *X*_2_ ∼ *Exp*(*λ*) are *i.i.d.*, 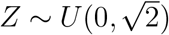. The derivations of these marginal PDFs and CDFs are provided in SI, §2.1.1.

Now that we have these CDFs, we can use them for outlier detection. After first deciding to fit exponential distributions to *X*_1_ and *X*_2_, we use the `vglm` function in the `R` package `VGAM` to fit an asymmetric Laplace distribution to the difference Δ. We could alternatively fit an exponential distribution to each data vector independently and obtain the MLEs 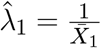 and 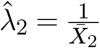 (Rice, 2007). We take the former approach instead so that we can get a 95%-confidence interval for log *κ*, deciding to use the symmetric case if this interval includes 0 (with 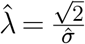) and the asymmetric case if it does not include 0 (with 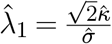 and 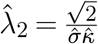). We can still test the hypothesis *θ* = 0 for the asymmetric Laplace distribution, even though our derivation of the distributions for Δ and *Z* above did not assume this. If we determine that 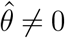, we can use the same probability functions from before for 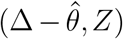 instead of (Δ, *Z*).

Once we have either 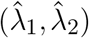 or 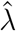, we decide whether we will detect outliers using the joint or marginal CDFs above. If we have reason to believe there should be a heavily-populated band around Δ = 0 on the (Δ, *Z*)-plot, for example due to consistent measurement error, then we will use the marginal method. We consider all of the points with values for Δ within *k* = 1 or *k* = 2 standard deviations (of Δ) away from 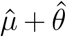 to not be outliers. We next calculate

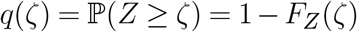

for the remaining points, without re-fitting them to another distribution. Using a pre-specified cutoff-value 0 < *q*^∗^ < 1 (like 0.05), we decide that all the points not in the central band with *q*(*ζ*) ≤ *q*^∗^ are outliers. We assign the probability *q* = 1 to points in the central band (for Δ) so that all points have a *q*-value, and all the points in the middle will have *q*(*δ, ζ*) ≥ *q*^∗^ for 0 ≤ *q*^∗^ ≤ 1.

If we decide to use the joint CDFs, for example when there is not a well-defined band around Δ = 0, then for each point, we calculate

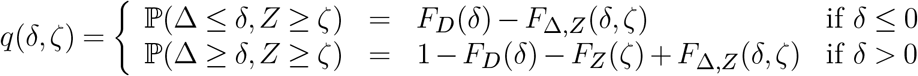

because *F*_*D*_(*δ*) = ℙ(Δ ≤ *δ*), *F*_*Z*_ (*ζ*) = ℙ(*Z* ≤ *ζ*), and *F*_Δ,*Z*_ (*δ, ζ*) = ℙ(Δ ≤ *δ, Z* ≤ *ζ*) by definition. Using the same value for *q*^∗^, we decide that all points with *q*(*δ, ζ*) ≤ *q*^∗^ are outliers. In practice, we will really calculate these probabilities for 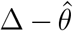, setting 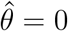 if the 95% confidence interval for *θ* includes 0.

The primary advantage of these methods over the previous one, besides the more accurate distribution for *Z*, is the interpretability. Instead of defining a set of parameters (*k, t, b, N*_*u*_, *N*_*l*_) and deciding whether to use a Weibull or log-normal distribution for *Z*, we get an estimated probability for every point after only deciding whether to use the joint or marginal method, the confidence level for *θ*, the confidence level for log *κ*, and the choice of *k* (in the marginal method). We can compare which points are more likely to be outliers than others based on the value of *q*(*δ, ζ*) or *q*(*ζ*) assigned to every pair of data points (*x*_1_, *x*_2_). The quantities *q*(*δ, ζ*) and *q*(*ζ*) quantify the reproducibility of a given pair of replicates and can be interpreted as their reliability. We can also adapt this method more easily when there are more than two replicates: we check every pair-wise comparison for outliers using this method and we divide the cutoff value *q*^∗^ by the number of comparisons (similar to a Bonferroni correction in multiple hypothesis testing) so that we do not over-estimate how many outliers there are (Rice, 2007).

## 4 Methods based on the generalized gamma model

While we were able to find closed-form CDFs for (Δ, *Z*) jointly and for Δ and *Z* marginally when we assumed *X*_1_ and *X*_2_ had independent exponential models, these cover only a fairly limited class of probability models. Exponential models only have one parameter *λ* > 0 and this parameter only determines the scale of the PDF, not the shape.

We might want to use a richer class of probability models with more parameters (which we would estimate using maximum likelihood methods). Both gamma and Weibull models generalize the exponential model, but we would then need to decide which one to use. Instead, we use the generalized gamma (GG) model (Jackson, 2016), of which the gamma, Weibull, and exponential models are all special cases. There are two different parameterizations for the generalized gamma PDF. For our purposes we will use the parametrization used by the `R` package `flexsurv`, given by Johnson and Kotz (1970)^*a*^. The PDF of *X*_*i*_ ∼ *GG*(*α*_*i*_, *β*_*i*_, *c*_*i*_) is

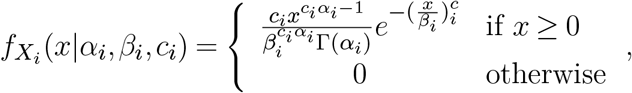

where *α*_*i*_ > 0, *β*_*i*_ > 0, and *c*_*i*_ > 0, *i* = 1, 2. The GG model has three different parameters we can tune (as opposed to the usual one or two for most probability models), so many familiar models with support on ℝ_+_ ⋃{0} are special cases. These include the gamma, exponential, chi-square, Weibull, half-normal, chi, Rayleigh and Maxwell-Boltzmann, thus making it an extremely flexible family that allows modeling of a variety of shapes. These models have been described in Rice (2007), Johnson and Kotz (1970)^*a*^, German (2010) and Stacy (1962). Specific details on the connection between these models and the GG family are provided in SI, §2.3. We cannot directly find a closed-form solution for the joint CDF or PDF for (Δ, *Z*) like we did under the exponential assumption. Instead, we will use a variable transform and change of variables from the joint PDF of (*X*_1_, *X*_2_), assuming *X*_1_ ∼ *GG*(*α*_1_, *β*_1_, *c*_1_) and *X*_2_ ∼ *GG*(*α*_2_, *β*_2_, *c*_2_) are independent. We will find MLEs for each of these six parameters and will not test the hypotheses that any of them are equal. The derivation of the joint density of (Δ, *Z*) is detailed in SI, §2.3.1, and this density is given by

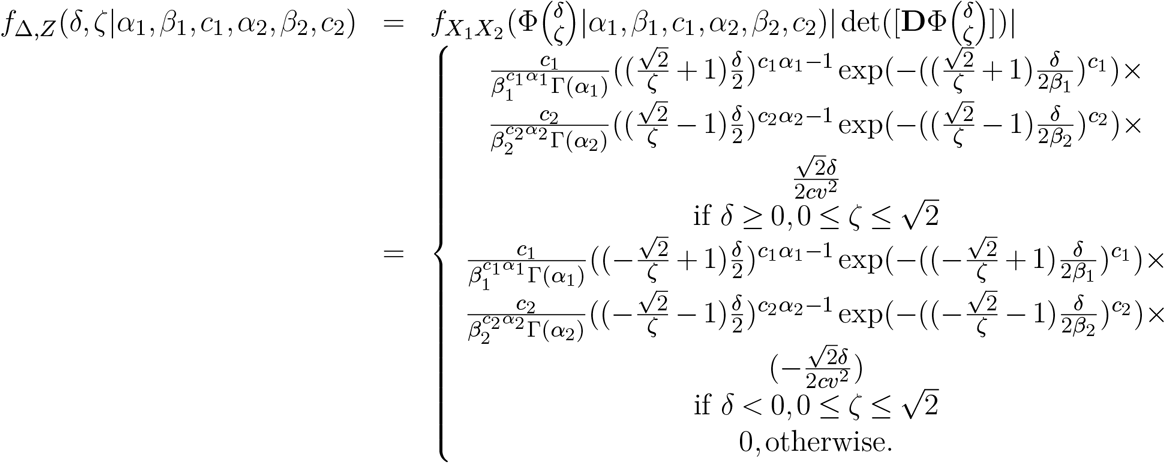

We do not have a closed-form solution for the joint CDF of (Δ, *Z*), but we can use numerical integration to find the probability that a point is not an outlier, such as the `adaptIntegrate` function in the `R` package `cubature` (Narasimhan and Johnson, 2018). We can use the `flexsurvreg` function in the `R` package `flexsurv` to get MLEs for 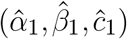 and 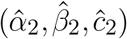 (Jackson, 2016). In some cases, it would be helpful to provide `flexsurvreg` with initial values for the parameters. For example, if we suspect the data follows a distribution similar to a Weibull, then we start with 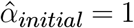. We then define

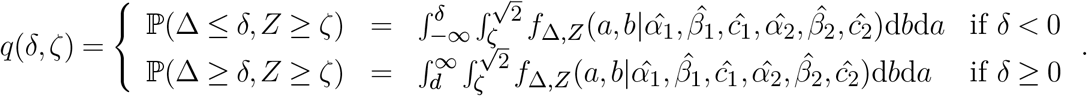

and the outlier determination parameter 0 < *q*^∗^ < 1. We decide that if (*x*_1_, *x*_2_) has *q*(*δ, ζ*) ≤ *q*^∗^, then (*x*_1_, *x*_2_) is an outlier.

We cannot directly adapt the marginal method to the GG assumption, since we would need the marginal mean and standard deviation for Δ. Our work-around will be to fit GG models to *X*_1_ and *X*_2_ to get MLEs for 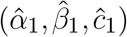 and 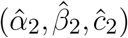, and also to fit an asymmetric Laplace model to Δ = *X*_1_ − *X*_2_ like we did before. We use 95%-confidence intervals to determine if the parameters *κ* or *θ* should be 1 or 0, respectively. We then filter out the points with Δ within *k* = 1 or *k* = 2 standard deviations from 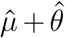 as non-outliers (assigning them *q*(*δ, ζ*) = 1). Even though the difference of two independent GG random variables is only an asymmetric Laplace distribution in the special case of exponentially distributed data, we make this simplification because there is not a closed-form solution in the more general cases. However, empirical evidence suggests that the asymmetric Laplace model closely approximates the distribution of Δ in this case.

For the remaining points, we use the estimates for 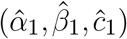 and 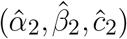 calculated from all of the points to find the marginal probabilities. Using the expression for 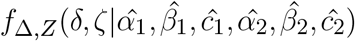 above, the marginal PDF for *Z* is

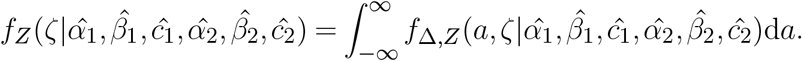

Like before, we can accomplish this integration numerically with the `adaptIntegrate` function in the `R` package `cubature` (Narasimhan and Johnson, 2018). Then the probability that we observe a value *ζ* for *Z* is

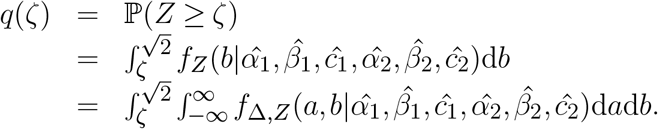

We define an outlier determination parameter 0 < *q*^∗^ < 1 and decide that if (*x*_1_, *x*_2_) was not removed as part of the middle band on the (Δ, *Z*) plot and has *q*(*ζ*) ≤ *q*^∗^, then (*x*_1_, *x*_2_) is an outlier. We can adapt these pairwise methods to the case of more than two replicates by using adjusted values for *q*^∗^ as outlined before. Methods based on the Weibull and gamma models, two important special cases of GG, are outlined in SI, §2.2.

## 5 Application to simulated data

We will now simulate data that looks like the “sideways volcano plot” and use the joint and marginal methods to find the outliers. We will create these plots by simulating exponentially distributed data for *X*_1_ and *X*_2_ to populate points in the middle band of these plots. For the “wings,” we will generate data from gamma, Weibull, or GG models for *X*_1_ and *X*_2_ with different parameters tuned to make the overall (Δ, *Z*)-plot look like the plots from real data (Supplementary Figures 4, 5 & 6, respectively). In each case, there will be a total of 104 data points.

To find outliers among the simulated data, we use the original asymmetric Laplace-Weibull method, the joint and marginal exponential methods, the joint and marginal GG model-based methods. The parameters for these methods will be the defaults: *k* = 1, *t* = *b* = 0.01, *N*_*u*_ = *N*_*l*_ = 1, and 95%-confidence intervals for *θ* and *κ*. The asymmetric Laplace-Weibull method divides the points by putting about 78.1% in the middle band (Status = 0), 21.1% in the lower “wings” (Status = 1), and 0.7% in the outlier regions (Status = 2). The joint exponential method identifies about 0.4% of pairs of observations as outliers at the *q*^∗^ = 10^−4^ cutoff level. For GG model-based methods, we use the package `parallel` that is part of the `R` base package (R Core Team, 2018). We plot the calculated probabilities and outlier statuses in Supplementary Figures 7, 8 & 9.

The joint methods seem to achieve the desired results for all three types of simulated data. For GG data, some of the points with small Δ but *Z* close to 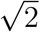 may have probabilities smaller than what we calculate for points with larger Δ values, thus indicating that these are outliers (Supplementary Figure 9). Since most of the middle band still contains smaller *Z* values, we can ignore this problem for few points with large *Z* values. These only occur with small Δ values when both *X*_1_ and *X*_2_ are close to 0. These methods can also consider points that are just slightly off of the middle band as non-outliers. Overall, the performance of the joint methods is similar across the three data generating mechanisms.

The marginal methods work the best for GG data, where the middle band is the thinnest. We need much larger values of the cut-off, *q*^∗^, to find outliers than we did when using the joint methods; this is acceptable because the newly proposed methods allow us to use different cut-offs for each data set. Unlike when we used the joint method, if a point has a value for Δ just slightly off of the middle band and a large *Z* value, then it can have a small *q*, particularly for data generated from the gamma (Supplementary Figure 7) or Weibull model (Supplementary Figure 8). If we have reason to believe or expect that that there will be points in this region, then the joint method - exponential or GG - would be the appropriate methods to use, depending on how much computational power we can afford to use.

## 6 Application to high-throughput biological data

Next we will use the methods developed in this paper to find outliers in real biological data. We will utilize the compound screening data from Anastassiadis et al. (2011, 2013) described in 1 and also data from a proteomics study in which triplicate measurements of protein expression of kinases were obtained from leukemia cell lines. These data sets were generated in two different laboratories at FCCC. Where we encounter missing replicates in the triplicate proteomics data set, we have a choice of computing the minimum *q* value only for the pairs present or computing *q*_min_ only for sets with all three replicates (a complete case analysis). For simplicity, we will do the latter.

### 6.1 Compound screening

We can see from the plot of the kinase inhibitor data in Supplementary Figure 1 that we do not have the clearly-defined sections needed for optimal use of the asymmetric Laplace-Weibull and marginal methods. The joint method will probably work the best. The maximum likelihood estimation for the GG model in the `R` function `flexsurvreg` assumes the data is non-zero; hence we need to replace all zeros in this data set with a small value (0.1) (Jackson, 2016). We create the (Δ, *Z*)-plot for this data colored by the outlier status and calculated probabilities in Figure 1. To better compare the different outputs, we use different outlier cutoff values *q*^∗^ (Figures 2, 3, 4 & 5).

**Figure 1:**
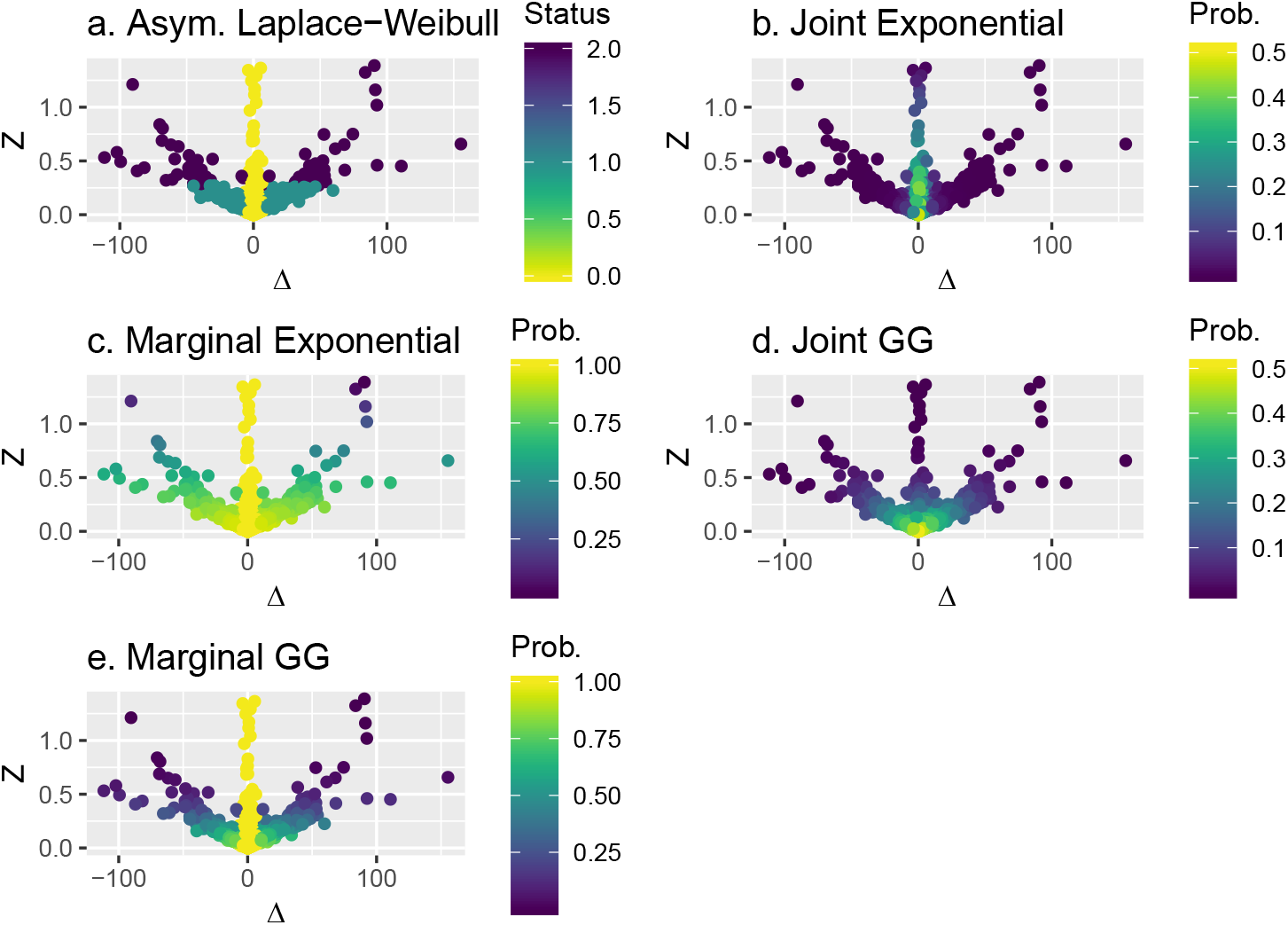
(Δ, *Z*)-plot for the kinase-inhibitor data labeled by the method used to identify outliers and colored by the outlier status and calculated probabilities. We color the plots in the same way we did in Supplementary Figure 7 and use the same R packages to make them (Wickham, 2016; Auguie, 2017; Garnier, 2018). To avoid overplotting, we only plot 20% (chosen randomly) of the points in each graph, using the function ‘sample frac’ function in the `R` package `dplyr` (Wickham et al., 2018).

**Figure 2:**
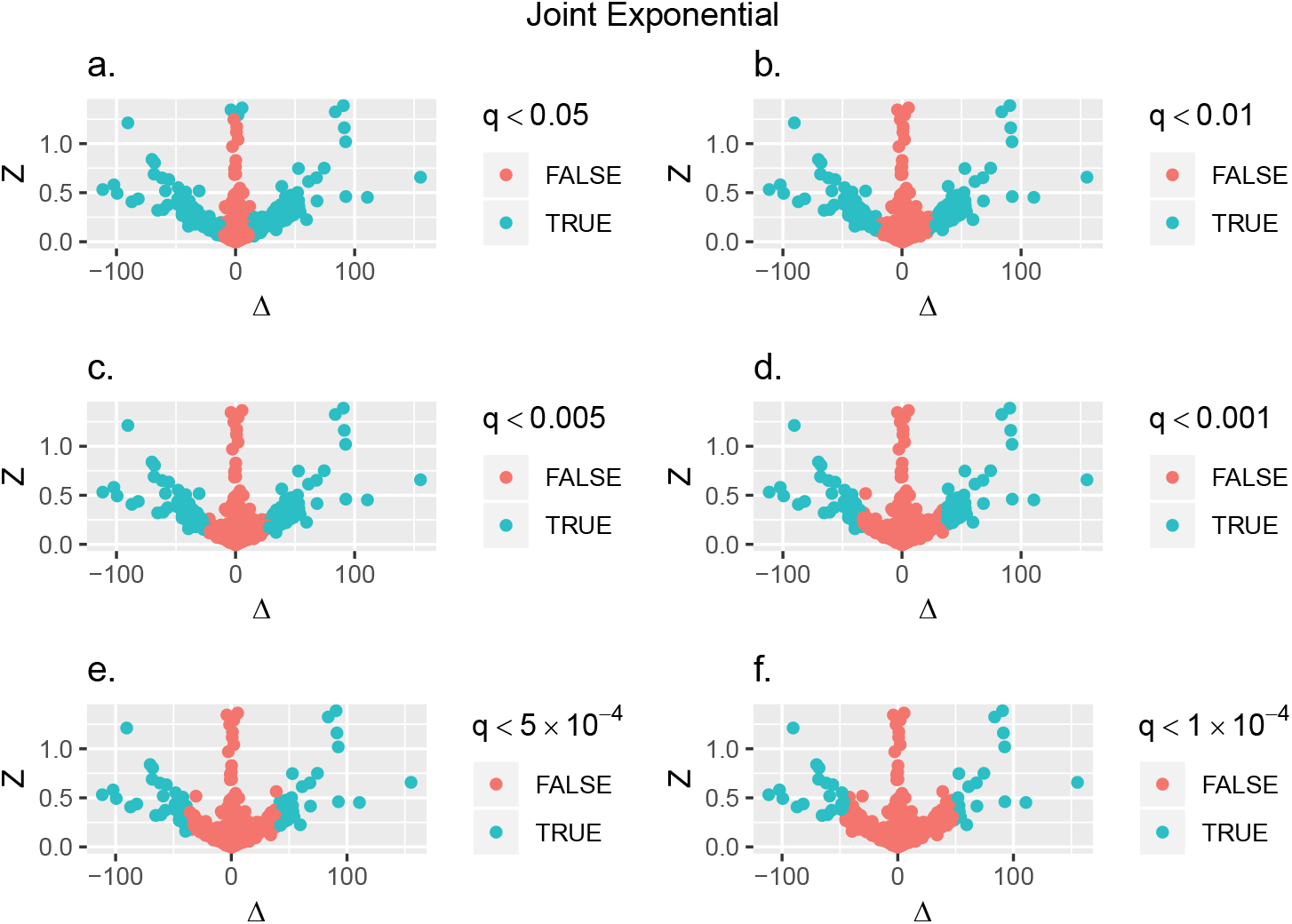
The kinase-inhibitor data with probabilities calculated using the joint exponential method, and with colors determined by various cutoff values *q*^∗^. We use the `R` packages `ggplot2` and `gridExtra` to make them (Wickham, 2016; Auguie, 2017).

**Figure 3:**
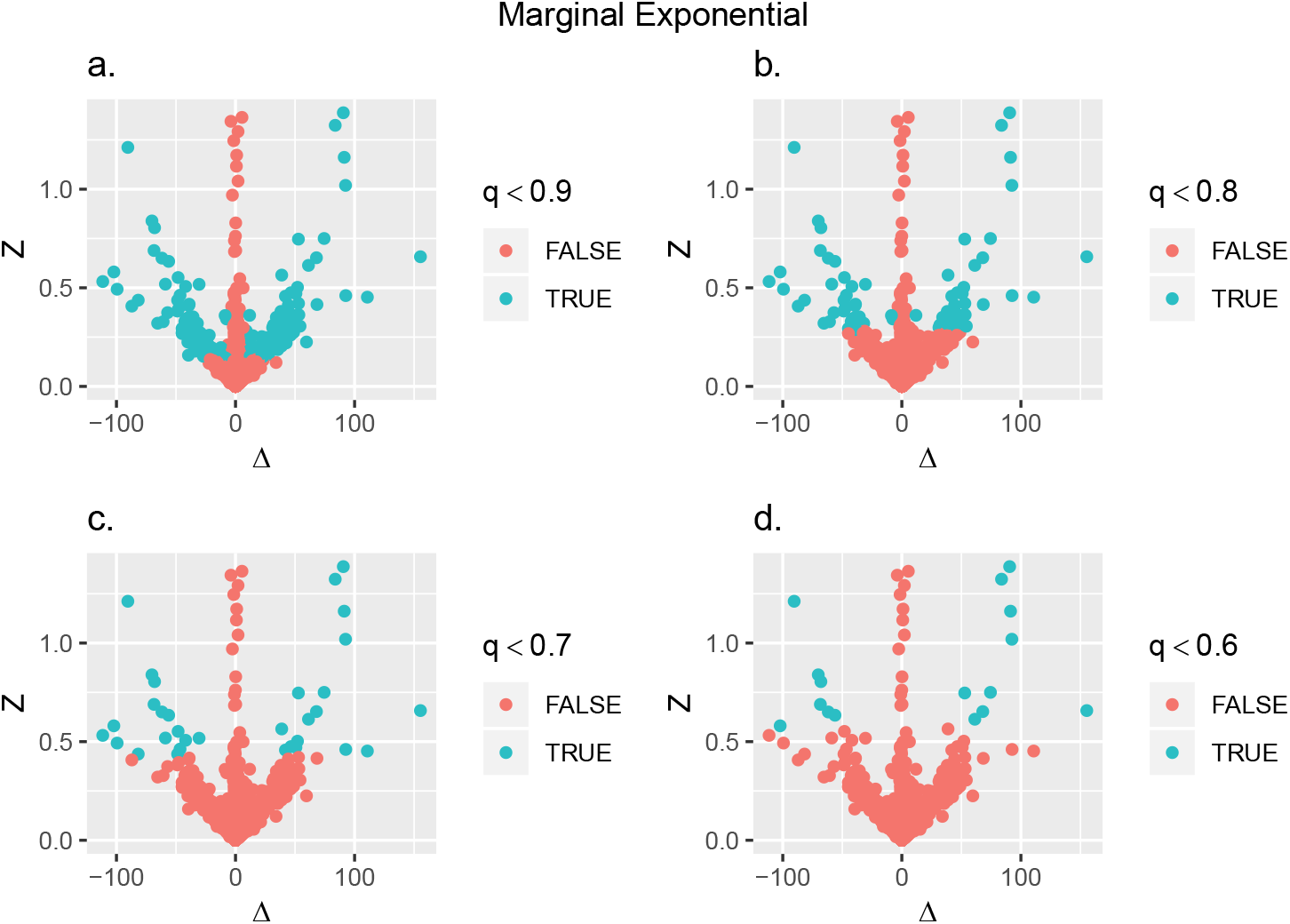
The kinase-inhibitor data with probabilities calculated using the marginal exponential method, and with colors determined by various cutoff values *q*^∗^. We use the `R` packages `ggplot2` and `gridExtra` to make them (Wickham, 2016; Auguie, 2017). Both the joint and marginal exponential methods prioritize keeping data in the middle band.

**Figure 4:**
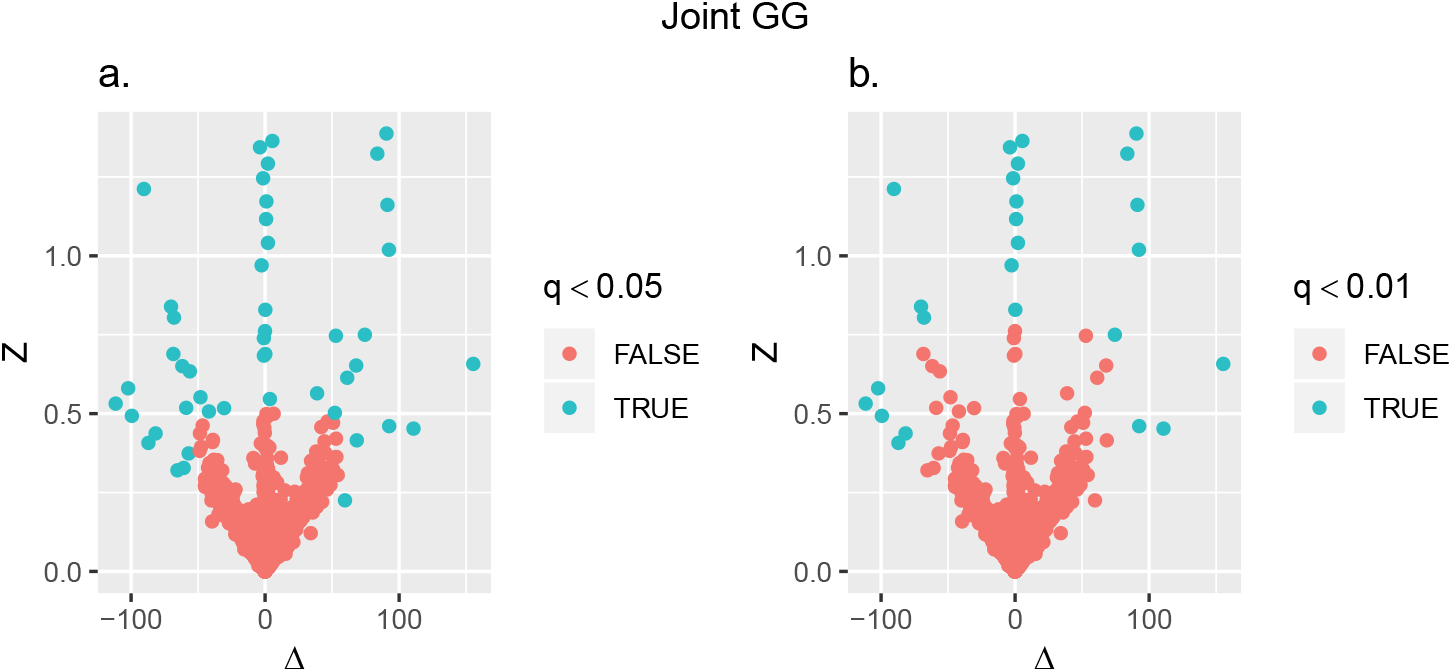
The kinase-inhibitor data with probabilities calculated using the joint GG method, and with colors determined by various cutoff values *q*^∗^. We use the `R` packages `ggplot2` and `gridExtra` to make them (Wickham, 2016; Auguie, 2017). Unlike both exponential methods, the joint GG method emphasizes *Z* more than Δ in finding outliers. At a *q*-cutoff that leaves around the same proportion of points as outliers (between 0.1% and 0.5% of the points), the joint GG method considers points with modest *Z* but small Δ to be outliers.

**Figure 5:**
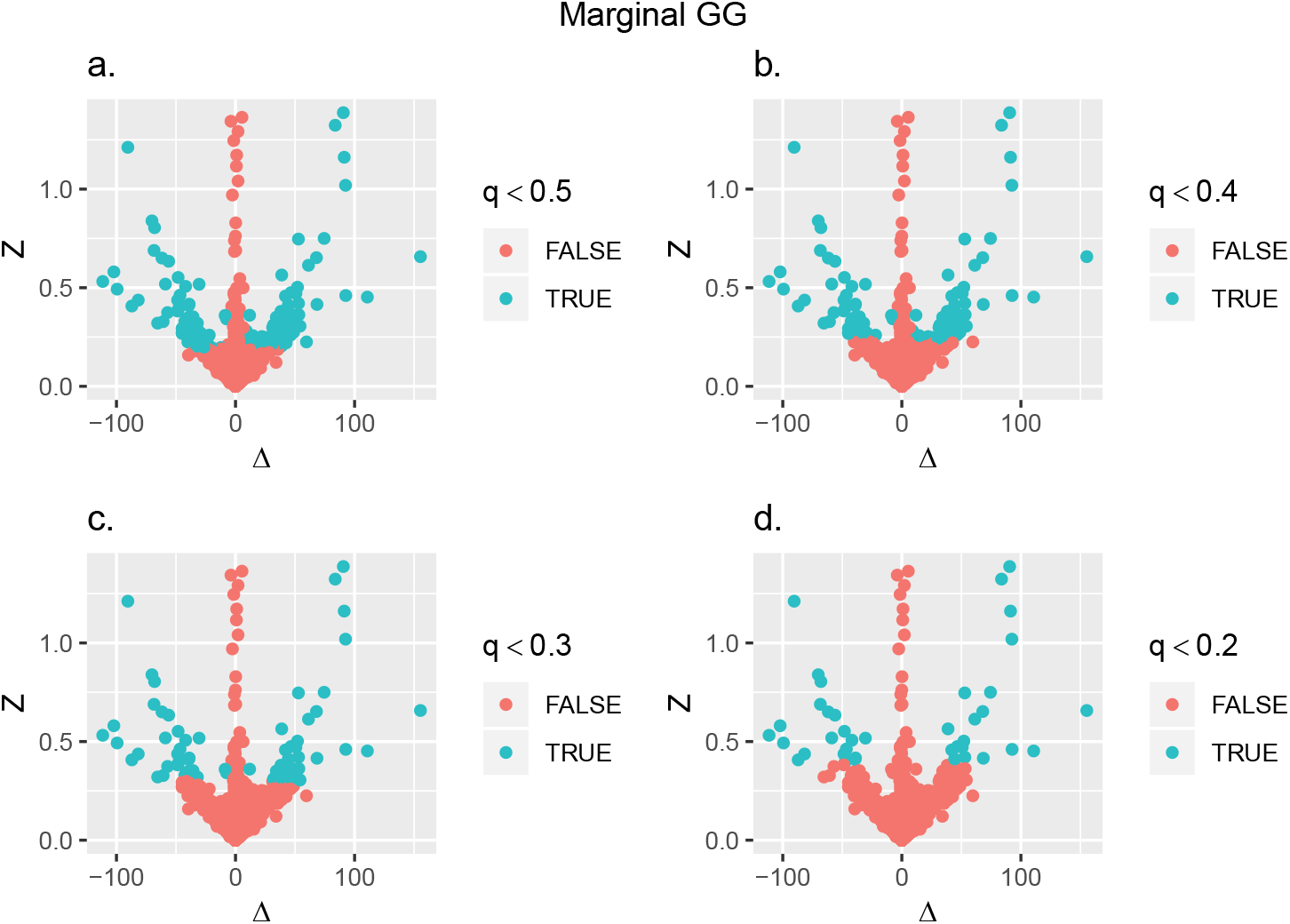
The kinase-inhibitor data with probabilities calculated using the marginal GG method, and with colors determined by various cutoff values *q*^∗^. We use the `R` packages `ggplot2` and `gridExtra` to make them (Wickham, 2016; Auguie, 2017). In both marginal methods, we prioritize keeping points with small Δ, regardless of the *Z* value, so these plots look more similar to the marginal exponential plots than the joint GG ones.

The joint exponential method identifies about 0.4% of the kinase-inhibitor pairs as outliers at the *q*^∗^ = 10^−4^ cutoff level. Except for the few points clustered around 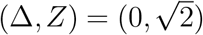 (which occur when both *X*_1_ and *X*_2_ are close to 0), this method identifies as outliers all of the points with large Δ (Figure 2). This method requires a much smaller *q*^∗^ value to find roughly the same numbers of outliers as the other three methods. The marginal exponential method identifies about 0.6% of the points as outliers at the *q*^∗^ = 0.8 cutoff level and about 0.2% at the *q*^∗^ = 0.7 cutoff level (Figure 3). The joint GG model-based method identifies about 0.4% of the points as outliers when *q*^∗^ = 0.05 and about 0.2% as outliers when *q*^∗^ = 0.01 (Figure 4). At *q*^∗^ = 0.3, the marginal GG model-based method considers about 0.5% of the points to be outliers, and at *q*^∗^ = 0.2, it considers about 0.2% of the points to be outliers (Figure 5). The different methods seem to agree on how many outliers there are, but not on exactly which points are the outliers.

Since so many points have Δ ≈ 0 in the volcano plot, both marginal methods consider all of the points away from this line to have significant Δ values. The marginal methods then only use the *Z* of these remaining points to calculate *q* and find outliers. The joint exponential method uses both Δ and *Z*, and considering how many points are around Δ = 0, the joint method considers points with larger Δ values to be outliers first. Unlike other methods, the joint GG model-based method seems to consider many of the points in the middle band to be outliers, i.e., points with high *Z* and Δ ≈ 0, instead keeping points with small *Z* and larger Δ values as non-outliers. The joint exponential and and marginal GG methods help isolate the middle band and some of the “wings” from the remainder of the points with smaller cut-offs *q*^∗^ and, thus, offer a flexible and more practical approach for identifying outliers.

### 6.2 Protein expression

Next, we consider proteomics data consisting of protein expression of kinases obtained using Multiplexed Inhibitor Bead (MIB) assays from leukemia cell lines. The measurements represent kinase SILAC (stable isotope labeling by amino acids in cell culture, Ong et al., 2002) ratios relative to a super-SILAC reference sample. Lysates from cancer cells were mixed in a 1 : 1 ratio with a super SILAC reference sample and protein kinases isolated using MIB assays and subjected to liquid chromatography with tandem mass spectrometry (LC-MS/MS). Quantitation of kinase levels was performed using MaxQuant Software (www.maxquant.org) as a ratio of kinase to super-SILAC reference sample (Kurimchak et al., 2019).

These experiments were done in three sets of triplicates for each of 234 kinases and resulted in a total of 579 with complete data. Thus, it provides us with the opportunity to evaluate our methods when there are more than two replicates. Since the original asymmetric Laplace-Weibull method does not produce a *q*-value, we can only test the joint and marginal methods (for both the exponential and GG assumptions). For the remaining four methods, we calculate the probabilities *q* for each pair-wise comparison of replicates (*X*_1_ vs. *X*_2_, *X*_2_ vs. *X*_3_, and *X*_3_ vs. *X*_1_) and determine the minimum *q*-value for each triplicate for each method. For both marginal methods, we assign *q* = 1 in the middle band. We plot the empirical distributions of the minimum *q*-values in Supplementary Figure 10.

The empirical plots in Supplementary Figure 10 suggest that we use cutoffs of 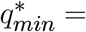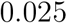 for the joint exponential method, 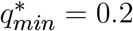 for the marginal exponential method, 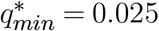 for the joint GG method, and 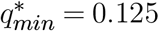 for the marginal GG method. Out of 579 total data points (each in triplicate), these cutoffs identify 70, 3, 4, and 3 outliers, respectively. We should probably use a smaller cutoff value for the joint exponential method, and after decreasing 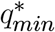 to 10^−4^ we finally get 8 remaining outliers. Since the joint exponential method produces so many more *q*_*min*_ values clustered around 0, we probably should rely on one of the other three methods for finding the outliers in this data set.

## 7 Summary and Discussion

We presented five different methods for finding outliers in high-throughput biological data and highlighted their utility in novel applications involving experimental data from cancer studies. Ideally, one would use exponential model-based methods when there are enough data points where computational intensity becomes an issue but when one still wants more accuracy than the original asymmetric Laplace-Weibull method. One should also check the empirical distribution of (Δ, *Z*) in order to choose a joint probability or marginal probability model-based method. For exponential and GG model-based methods, the estimated probabilities allow the user to set their own cutoff for finding outliers, allowing the analysis to be as conservative as needed. These four methods also allow different schemes for comparing the probabilities calculated for each pair of replicates when there are more than 2 replicates. Our analysis assigned the minimum *q*-value to each trio of measurements, but other approaches (like using the maximum, mean, or median) are possible. There are also more schemes for dealing with missing data than the complete case analysis used for the triplicate proteomics data.

The methods developed in this paper are implemented in the `R` package `replicateOutliers` which is available at Github (www.github.com/matthew-seth-smith/replicateOutliers). While the asymmetric Laplace-Weibull method and exponential methods run quickly and reliably, the GG model-based methods’ reliance on two-dimensional numerical integration makes the run time of their implementation longer. We needed to run numerical integration in parallel on FCCC’s High-Performance Computing (HPC) cluster, while the other three methods worked quickly on a laptop. A total of 10 nodes with 8 cores each, and two 2.4 GHz Intel E5-2620 v3 CPUs per core, with 128Gb of memory per node were utilized for this purpose on the HPC cluster. GG model-based methods also require the data to be non-zero, meaning any zeros in a real data set should be set to some small positive value. Exactly which small positive value (or values) we choose is of course arbitrary and depends on the scale of the positive data. One approach is to use randomly generated data from *U* (0, *ϵ*) for some small *ϵ* > 0 to replace the zeros. If possible, the user should still try to use GG model-based methods because of their flexible structure; and fitting more parameters to the data is likely to make outlier identification more accurate. This is made possible by the large volume of data available in high-throughput studies. Overall, GG model-based methods offer modeling flexibility while exponential model-based methods offer a faster approach based on a closed form solution to evaluating the reproducibility of replicates. We were able to see increased accuracy in the triplicate proteomics data analysis, where marginal exponential and both GG model-based methods roughly agreed on the number of outliers, while the joint exponential method did not.

The cut-offs *q*^∗^ play an important role in identifying outliers and its choice should be data driven and utilize any prior knowledge on the data generating mechanism. It may not be possible to use the same *q*^∗^-value for every method because of variations in their scales. This becomes apparent from comparing the joint and marginal methods. For each method, there might be a way to set standards for *q*^∗^; but it would really depend on the specific data set at hand. For example, it would depend on the number of replicate pairs; the numbers of points in the central band, “wings,” and “upper probable-outlier regions”; and how close the wings and upper regions are. A practical approach would be to try different *q*^∗^-values in order to identify the ones that give an acceptable outlier frequency, and then finding those outliers. If a joint method gives a high outlier frequency even with a large *q*^∗^ cutoff, then it suggests that the data are not of good quality, i.e., not reproducible. Overall, our results indicate that the underlying model assumptions are much less important when applying these methods to high-throughput biological data. Each proposed method provides unique insight and offers a way to interpret the data; in addition, methods based on other special cases of interest such as the Weibull and gamma are detailed in SI, §2.2.

Although we developed these methods with high-throughput biological data in mind, we can easily apply them to other biomedical data, public health screening, and testing the performance of mechanical and electronic devices. The approaches used in developing these methods can also lead to creating schemes for data following a known distribution, including both bounded intervals and all of ℝ.

## Acknowledgements

The work of MSS was supported by a Summer Undergraduate Research Fellowship at FCCC while he was pursuing his dual degrees in mathematics and biostatistics at Yale University as part of its five year BS/MPH program. The work of KD was supported in part by NIH grant P30 CA 006927. The authors are grateful to Dr. Jeffrey Peterson and Dr. James Duncan of the Cancer Biology Program at FCCC for providing their data sets to be used in the evaluation of the proposed methods.

