## Supplemental Information for "Probability-based methods for outlier detection in replicated high-throughput biological data"

#### 1 Interpretation of the Laplace model

Purdom and Holmes (2005) provide some insight into the potential interpretation of Laplace models for microarray gene expression data. In this section, we extend their ideas and argue that the Laplace model provides a good fit to replicate data obtained from a variety of high-throughput studies such as compound and siRNA screening, next-generation RNA sequencing and SNP arrays. The random variable $X$ in equation (2.1) (in the main paper) can be expressed as the log-ratio of two independent random variables, $Y_1$ and $Y_2$, with Type I Pareto ($PI$) distributions. Using the PDF of a random variable $Y$ with Type I Pareto distribution, given by

$$f_Y(y) = \frac{\alpha\beta^\alpha}{y^{\alpha+1}}, \quad y > \beta, \alpha > 0, \beta > 0, \tag{1.1}$$

we obtain

$$X = \mu + \frac{\sigma}{\sqrt{2}} \log\left(\frac{Y_1}{Y_2}\right) \tag{1.2}$$

where $Y_1 \sim PI(\alpha = \kappa, \beta = 1)$ and $Y_2 \sim PI(\alpha = 1/\kappa, \beta = 1)$ and $\kappa$ is as in equation (2.1) (of the paper). Purdom & Holmes (2005) employ this reasoning and demonstrate that two-color microarray data, expressed as the log-ratio of the red and green channel intensities, exhibits the $AL$ distribution. It is evident from equation (1.2) above that $\kappa \neq 1$ only if $Y_1$ and $Y_2$ are $PI$ with different parameters which implies that skewness in the data, quantified by $\kappa$, arises from a difference in channel distributions. For replicate data from high-throughput studies, we demonstrate that skewness in the data arises from a difference in the distribution of duplicates. Other applications of Pareto distributions for high-throughput data are also found in the literature. For example, Kuznetsov (2001) proposes the use of Type II Pareto distributions, a variant of $PI$ with a location parameter, for mRNA expression data in Serial Analysis of Gene Expression (SAGE) studies. It is important to note that data obtained in SAGE studies is similar in structure to digital gene expression (DGE) data obtained from next-generation sequencing technologies such as RNA-Seq, ChIP-Seq etc. (Robinson and Smyth, 2008). Furthermore, Wu et al. (2003) showed that the distribution of PM expression intensities in oligonucleotide Affymetrix microarray data resembled the $PI$ distribution. Thus there is some evidence from prior studies to support the use of $AL$ in DGE data and expression data from SNP arrays.

If $Y \sim PI(\alpha = 1, \beta = 1)$, it is well-known that $W = \log Y$ has a standard exponential distribution with density $f_W(w) = e^{-w}, w > 0$. Using this result and equation (1.2) above, the random variable $X$ can be expressed as a linear combination of two independent and identically distributed (*i.i.d.*) standard exponential random variables $W_1$ and $W_2$,

$$X = \mu + \frac{\sigma}{\sqrt{2}} \left( \frac{1}{\kappa} W_1 - \kappa W_2 \right). \tag{1.3}$$

#### 1.1 The generalized asymmetric Laplace model

A random variable $Z$ is said to have the generalized asymmetric Laplace ($GAL$) distribution if its characteristic function is given by

$$\psi(t) = \frac{e^{i\mu t}}{\left(1 + \frac{1}{2}\sigma^2 t^2 - i\nu t\right)^{\tau}}, -\infty < t < \infty \tag{1.4}$$

where $\mu, \nu \in \Re$, $\sigma, \tau \geq 0$ and $\nu = \frac{\sigma}{\sqrt{2}} \left( \frac{1}{\kappa} - \kappa \right)$. The $GAL$ distribution is also known as the variance-gamma distribution or the Bessel function distribution due to the involvement of the Bessel function of the third kind in its PDF (Kotz et al., 2001). This PDF has a complex mathematical form and hence we have defined it using its characteristic function here. The $GAL$ distribution places the $AL$ family within a larger class of distributions that are useful for modeling replicate data in high-throughput studies. When $\tau = 1, \sigma > 0$ and $\nu \neq 0$ we obtain the $AL$ distribution and if $\tau = 0, \sigma > 0$ and $\nu = 0$ we get the $SL$ distribution. The usefulness of the class of distributions represented by $Z$ can summarized in the expression,

$$Z = \mu + \frac{\sigma}{\sqrt{2}} \left( \frac{1}{\kappa} G_1 - \kappa G_2 \right), \tag{1.5}$$

where $G_1$ and $G_2$ are *i.i.d.* gamma random variables with density $g(x) = \frac{1}{\Gamma(\tau)} x^{\tau - 1} e^{-x}$, $x > 0, \tau > 0$. This includes the representations given in equations (1.3) and (1.4) above as special cases when $\tau = 1$.

### 2 Derivation of joint and marginal models

In this section, we show the algebra and calculus in deriving $F_{\Delta,Z}$ for exponential data in both the symmetric and asymmetric cases. Using this function, we can find the marginal CDF and PDF for $Z$ when $X_1$ and $X_2$ are independent and exponentially distributed. Next, we derive the marginal distribution for $Z$ when $X_1$ and $X_2$ are independent and have Weibull distributions with the same shape parameter $c$. This new distribution will be a generalization of the one from the exponential case and will similarly have a closed-form solution. In addition, we consider the case of the gamma model. Lastly, we provide specific details for the case of the generalized gamma model.

#### 2.1 Exponential model

##### 2.1.1 Asymmetric case

We begin by defining two independent random variables $X_1 \sim Exp(\lambda_1)$ and $X_2 \sim Exp(\lambda_2)$. If $X \sim Exp(\lambda)$, its PDF is

$$f_X(x) = \begin{cases} \lambda e^{-\lambda x} & \text{if } x \geq 0 \\ 0 & \text{otherwise} \end{cases}.$$

Then if $X_1$ and $X_2$ are independent, their joint PDF is just

$$f_{X_1 X_2}(x_1, x_2) = f_{X_1}(x_1) f_{X_2}(x_2) = \begin{cases} \lambda_1 \lambda_2 e^{-\lambda_1 x_1} e^{-\lambda_2 x_2} & \text{if } x_1 \geq 0, x_2 \geq 0 \\ 0 & \text{otherwise} \end{cases}.$$

Now we define the two random variables of interest: $\Delta = X_1 - X_2$ and $Z = \frac{\sqrt{2}|X_1 - X_2|}{X_1 + X_2}$. Let $\delta$ be a value that $\Delta$ can take and $\zeta$ be a value that $Z$ can take. We wish to find the joint CDF $F_{\Delta,Z}(\delta,\zeta)$ but note that there is a high degree of dependence between $\Delta$ and $Z$. We cannot even find the PDF for $Z$ easily as the ratio of two independent variables, since the numerator and denominator there are clearly not independent. Lastly, since the function (from $\mathbb{R}^2$ to $\mathbb{R}^2$) from $(x_1, x_2)$ to $(\delta, \zeta)$ does not have an invertible derivative for every point in $\mathbb{R}^2$, we cannot do a change of variables and use $f_{X_1,X_2}(x_1, x_2)$ to find the joint PDF $f_{\Delta,Z}(\delta,\zeta)$. Hence, we need to find the CDF directly.

The CDF is defined as the joint probability that $\Delta$ and $Z$ take values less than or equal to $\delta$ and $\zeta$, respectively. Then

$$\begin{aligned} F_{\Delta,Z}(\delta,\zeta) &= \mathbb{P}(\Delta \leq \delta, Z \leq \zeta) \\ &= \mathbb{P}(X_1 - X_2 \leq \delta, \tfrac{\sqrt{2}|X_1 - X_2|}{X_1 + X_2} \leq \zeta) \\ &= \mathbb{P}(X_1 - X_2 \leq \delta, |X_1 - X_2| \leq \tfrac{Z}{\sqrt{2}}(X_1 + X_2)). \end{aligned}$$

Since $X_1$ and $X_2$ are both exoponentially distributed, they are both non-negative. So $X_1 + X_2$ is also non-negative and we do not have to change the $\leq$

(yet). To deal with the absolute value, we must split up this region into two smaller, disjoint ones:

$$\begin{aligned}
F_{\Delta,Z}(\delta,\zeta) &= \mathbb{P}(X_1-X_2\leq\delta, |X_1-X_2|\leq\tfrac{\zeta}{\sqrt{2}}(X_1+X_2), X_1-X_2\geq 0)\\
&\quad+\mathbb{P}(X_1-X_2\leq\delta, |X_1-X_2|\leq\tfrac{\zeta}{\sqrt{2}}(X_1+X_2), X_1-X_2< 0)\\
&= \mathbb{P}(X_1-X_2\leq\delta, X_1-X_2\leq\tfrac{\zeta}{\sqrt{2}}(X_1+X_2), X_1-X_2\geq 0)\\
&\quad+\mathbb{P}(X_1-X_2\leq\delta, -(X_1-X_2)\leq\tfrac{\zeta}{\sqrt{2}}(X_1+X_2), X_1-X_2< 0)\\
&= \mathbb{P}(X_1-X_2\leq\delta, X_1-X_2\leq\tfrac{\zeta}{\sqrt{2}}(X_1+X_2), X_1-X_2\geq 0)\\
&\quad+\mathbb{P}(X_1-X_2\leq\delta, X_1-X_2\geq-\tfrac{\zeta}{\sqrt{2}}(X_1+X_2), X_1-X_2< 0)\\
&= \mathbb{P}(0\leq X_1-X_2\leq\delta, X_1-X_2\leq\tfrac{\zeta}{\sqrt{2}}(X_1+X_2))\\
&\quad+\mathbb{P}(X_1-X_2<\min(\delta,0), X_1-X_2\geq-\tfrac{c}{\sqrt{2}}(X_1+X_2))\\
&= \mathbb{P}(0\leq X_1-X_2\leq\min(\delta,\tfrac{\zeta}{\sqrt{2}}(X_1+X_2)))\\
&\quad+\mathbb{P}(-\tfrac{\zeta}{\sqrt{2}}(X_1+X_2)\leq X_1-X_2\leq\min(\delta,0)).
\end{aligned}$$

We now break the first of the two regions into two smaller ones, depending on if $\frac{\zeta}{\sqrt{2}}(X_1+X_2)$ or $\delta$ is smaller. Then

$$\begin{aligned}
F_{\Delta,Z}(\delta,\zeta) &= \mathbb{P}(0\leq X_1-X_2\leq\delta, \delta<\tfrac{\zeta}{\sqrt{2}}(X_1+X_2)))\\
&\quad+\mathbb{P}(0\leq X_1-X_2\leq\tfrac{\zeta}{\sqrt{2}}(X_1+X_2), \tfrac{\zeta}{\sqrt{2}}(X_1+X_2)\leq\delta)\\
&\quad+\mathbb{P}(-\tfrac{\zeta}{\sqrt{2}}(X_1+X_2)\leq X_1-X_2\leq\min(\delta,0)).
\end{aligned}$$

We call these three regions $A$, $B$, and $C$, respectively. We can now simplify the expressions for each of these regions. The final forms will depend on $\delta$ and $\zeta$. For $A$, the equations simplify to

$$A=\{(x_1,x_2)\in\mathbb{R}^2 | x_2\leq x_1, x_2\geq x_1-\delta, x_2>-x_1+\frac{\sqrt{2}}{\zeta}\delta\}.$$

For $B$, we have

$$B=\{(x_1,x_2)\in\mathbb{R}^2 | x_2\leq x_1, x_2\geq\frac{\sqrt{2}-\zeta}{\sqrt{2}+\zeta}x_1, x_2\leq -x_1+\frac{\sqrt{2}}{\zeta}\delta\}.$$

We did not need to change the direction of the second inequality for $B$ because $\zeta>1$. For $C$, we need to know if $\zeta\geq\sqrt{2}$, because if it is, a $\sqrt{2}-\zeta$ term that will be divided will be negative. So if $0 \le \zeta < \sqrt{2}$ (the CDF is only non-zero for $\zeta$ non-negative), the region is

$$C = \{(x_1,x_2) \in \mathbb{R}^2 | x_2 \le \frac{\sqrt{2}+\zeta}{\sqrt{2}-\zeta} x_1, x_2 \ge x_1 - \min(\delta,0)\}.$$

If $\zeta \ge \sqrt{2}$, then

$$C = \{(x_1,x_2) \in \mathbb{R}^2 | x_2 \ge \frac{\sqrt{2}+\zeta}{\sqrt{2}-\zeta} x_1, x_2 \ge x_1 - \min(\delta,0)\}.$$

In any case, $F_{\Delta,Z}(\delta,\zeta) = \mathbb{P}(A) + \mathbb{P}(B) + \mathbb{P}(C)$.

###### 2.1.1.1 Region $A$

We first make a simplification to the expression for $A$. Let $\beta = \frac{\sqrt{2}}{\zeta}\delta$. Since $F_{\Delta,Z}(\delta,\zeta)$ is only non-zero for $\zeta > 0$, the sign of $\beta$ is the sign of $\delta$. We also know that $f_{X_1,X_2}(x_1,x_2) = 0$ for $x_1 < 0$ or $x_2 < 0$. Thus we need to determine the intersection between $A$ and Quadrant I of $\mathbb{R}^2$. This depends on the slopes and intercepts of the lines in $\mathbb{R}^2$ that determinee $A$, which in turn depend on the values $\delta$ and $\zeta$ take. Graphs for the different cases are included. Again, $F_{\Delta,Z}(\delta,\zeta) = 0$ for $\zeta < 0$.

First, we look at $\delta > 0$ and $0 \le \zeta \le \sqrt{2}$. This means the intercept between $x_2 = x_1 - \delta$ and $x_2 = -x_1 + \beta$ is above the $x$-axis. So

$$\begin{aligned}
\mathbb{P}(A) &= \iint_A f_{X_1,X_2}(x_1,x_2)\mathrm{d}^2\vec{x} \\
&= \int_{\frac{\beta-\delta}{2}}^{\frac{\beta}{2}} \int_{\beta-x_2}^{\delta+x_2} \lambda_1\lambda_2 e^{-\lambda_1 x_1} e^{-\lambda_2 x_2}\mathrm{d}x_1\mathrm{d}x_2 \\
&\quad + \int_{\frac{\beta}{2}}^{\infty} \int_{x_2}^{x_2+\delta} \lambda_1\lambda_2 e^{-\lambda_1 x_1} e^{-\lambda_2 x_x}\mathrm{d}x_1\mathrm{d}x_2 \\
&= \int_{\frac{\beta-\delta}{2}}^{\frac{\beta}{2}} \lambda_2 e^{-\lambda_2 x_2}(-e^{-\lambda_1 x_1})\Big]_{\beta-x_2}^{\delta+x_2}\mathrm{d}x_2 + \int_{\frac{\beta}{2}}^{\infty} \lambda_2 e^{-\lambda_2 x_2}(-e^{-\lambda_1 x_1})\Big]_{x_2}^{x_2+\delta}\mathrm{d}x_2 \\
&= \int_{\frac{\beta-\delta}{2}}^{\frac{\beta}{2}} \lambda_2 e^{-\lambda_2 x_2}(e^{\lambda_1 x_2 - \lambda_1\beta} - e^{-\lambda_1 x_2 - \lambda_1 d})\mathrm{d}x_2 \\
&\quad + \int_{\frac{\beta}{2}}^{\infty} \lambda_2 e^{-\lambda_2 x_2}(e^{-\lambda_1 x_2} - e^{-\lambda_2 x_2 - \lambda_1 d})\mathrm{d}x_2 \\
&= \int_{\frac{\beta-\delta}{2}}^{\frac{\beta}{2}} \lambda_2(e^{-\lambda_1\beta}e^{(\lambda_1-\lambda_2)x_2} - e^{-\lambda_1 d}e^{-(\lambda_1+\lambda_2)x_2})\mathrm{d}x_2 \\
&\quad + \int_{\frac{\beta}{2}}^{\infty} \lambda_2 e^{-\lambda_2 x_2}e^{-\lambda_1 x_2}(1 - e^{-\lambda_1 d})\mathrm{d}x_2 \\
&= \int_{\frac{\beta-\delta}{2}}^{\frac{\beta}{2}} \lambda_2 e^{-\lambda_1\beta}e^{(\lambda_1-\lambda_2)x_2}\mathrm{d}x_2 - \int_{\frac{\beta-\delta}{2}}^{\frac{\beta}{2}} \lambda_2 e^{-\lambda_1 d}e^{-(\lambda_1+\lambda_2)x_2}\mathrm{d}x_2 \\
&\quad + \int_{\frac{\beta}{2}}^{\infty} \lambda_2(1 - e^{-\lambda_1 d})e^{-(\lambda_1+\lambda_2)x_2}\mathrm{d}x_2.
\end{aligned}$$

For the first integral, we assume that $\lambda_1 \neq \lambda_2$; if they are equal, then this is an integral of a constant function from $\frac{\beta-\delta}{2}$ to $\frac{\beta}{2}$, the result of which would just be $\frac{\lambda_2 e^{-\lambda_1 \beta}\delta}{2}$. Assuming they are not equal (that $X_1$ and $X_2$ are independent but not *i.i.d.*),

$$\begin{aligned}
\mathbb{P}(A) &= \frac{\lambda_2 e^{-\lambda_1 \beta}}{\lambda_1-\lambda_2} e^{(\lambda_1-\lambda_2)x_2}\Big]_{\frac{\beta-\delta}{2}}^{\frac{\beta}{2}} + \frac{\lambda_2 e^{-\lambda_1 d}}{\lambda_1+\lambda_2} e^{-(\lambda_1+\lambda_2)x_2}\Big]_{\frac{\beta-\delta}{2}}^{\frac{\beta}{2}} \\
&\quad + \frac{\lambda_2(1-e^{-\lambda_1 d})}{\lambda_1+\lambda_2}(-e^{-(\lambda_1+\lambda_2)x_2})\Big]_{\frac{\beta}{2}}^{\infty} \\
&= \frac{\lambda_2 e^{-\lambda_1 \beta}}{\lambda_1-\lambda_2}(e^{(\lambda_1-\lambda_2)\frac{\beta}{2}} - e^{(\lambda_1-\lambda_2)\frac{\beta-\delta}{2}}) \\
&\quad + \frac{\lambda_2 e^{-\lambda_1 d}}{\lambda_1+\lambda_2}(e^{-(\lambda_1+\lambda_2)\frac{\beta}{2}} - e^{-(\lambda_1+\lambda_2)\frac{\beta-\delta}{2}}) \\
&\quad + \frac{\lambda_2(1-e^{-\lambda_1 d})}{\lambda_1+\lambda_2}(e^{-(\lambda_1+\lambda_2)\frac{\beta}{2}} - 0) \\
&= \frac{\lambda_2}{\lambda_1-\lambda_2}(e^{-(\lambda_1+\lambda_2)\frac{\beta}{2}} - e^{-(\lambda_1+\lambda_2)\frac{\beta}{2}} e^{\frac{(\lambda_2-\lambda_1)\delta}{2}}) \\
&\quad + \frac{\lambda_2}{\lambda_1+\lambda_2}(e^{-\lambda_1 d} e^{-(\lambda_1+\lambda_2)\frac{\beta}{2}} - e^{-(\lambda_1+\lambda_2)\frac{\beta}{2}} e^{\frac{(\lambda_2-\lambda_1)\delta}{2}}) \\
&\quad + \frac{\lambda_2}{\lambda_1+\lambda_2}(e^{-(\lambda_1+\lambda_2)\frac{\beta}{2}} - e^{-\lambda_1 d} e^{-(\lambda_1+\lambda_2)\frac{\beta}{2}}) \\
&= \frac{\lambda_2}{\lambda_1-\lambda_2}(e^{-(\lambda_1+\lambda_2)\frac{\beta}{2}} - e^{-(\lambda_1+\lambda_2)\frac{\beta}{2}} e^{\frac{(\lambda_2-\lambda_1)\delta}{2}}) \\
&\quad + \frac{\lambda_2}{\lambda_1+\lambda_2}(-e^{-(\lambda_1+\lambda_2)\frac{\beta}{2}} e^{\frac{(\lambda_2-\lambda_1)\delta}{2}} + e^{-(\lambda_1+\lambda_2)\frac{\beta}{2}}) \\
&= \lambda_2 e^{-(\lambda_1+\lambda_2)\frac{\beta}{2}}(1 - e^{\frac{(\lambda_2-\lambda_1)\delta}{2}})(\frac{1}{\lambda_1-\lambda_2} + \frac{1}{\lambda_1+\lambda_2}) \\
&= \frac{2\lambda_1\lambda_2}{\lambda_1^2-\lambda_2^2} e^{-(\lambda_1+\lambda_2)\frac{\beta}{2}}(1 - e^{\frac{(\lambda_2-\lambda_1)\delta}{2}}) \\
&= \frac{2\lambda_1\lambda_2}{\lambda_1^2-\lambda_2^2} e^{-(\lambda_1+\lambda_2)\frac{\delta}{\sqrt{2}\zeta}}(1 - e^{\frac{(\lambda_2-\lambda_1)\delta}{2}}),
\end{aligned}$$

using $\beta = \frac{\sqrt{2}}{\zeta}\delta$.

Now we look at $\delta > 0$ and $\zeta > \sqrt{2}$. Since $\beta = \frac{\sqrt{2}}{\zeta}\delta$, we have $0 < \beta < \delta$. If this is the case, the intercept $(\frac{\beta+\delta}{2}, \frac{\beta-\delta}{2})$ between the lines $x_2 = x_1 - \delta$ and $x_2 = -x_1 + \beta$ is below $x_2 = 0$. But since $f_{X_1,X_2}(x_1, x_2)$ is zero here, we must change our bounds of integration to include $x_2 = 0$. So,

$$
\begin{aligned}
\mathbb{P}(A) &= \iint\limits_A f_{X_1,X_2}(x_1,x_2)\mathrm{d}^2\vec{x} \\
&= \int_{\frac{\beta}{2}}^{\beta}\int_{-x_1+\beta}^{x_1}\lambda_1\lambda_2 e^{-\lambda_1 x_1}e^{-\lambda_2 x_2}\mathrm{d}x_2\mathrm{d}x_1 + \int_{\beta}^{\delta}\int_0^{x_1}\lambda_1\lambda_2 e^{-\lambda_1 x_1}e^{-\lambda_2 x_2}\mathrm{d}x_2\mathrm{d}x_1 \\
&+ \int_d^{\infty}\int_{x_1-\delta}^{x_1}\lambda_1\lambda_2 e^{-\lambda_1 x_1}e^{-\lambda_2 x_2}\mathrm{d}x_2\mathrm{d}x_1 \\
&= \int_{\frac{\beta}{2}}^{\beta}\lambda_1 e^{-\lambda_1 x_1}(-e^{-\lambda_2 x_2})\Big]_{-x_1+\beta}^{x_1}\mathrm{d}x_1 + \int_{\beta}^{\delta}\lambda_1 e^{-\lambda_1 x_1}(-e^{-\lambda_2 x_2})\Big]_0^{x_1}\mathrm{d}x_1 \\
&+ \int_d^{\infty}\lambda_1 e^{-\lambda_1 x_1}(-e^{-\lambda_2 x_2})\Big]_{x_1-\delta}^{x_1}\mathrm{d}x_1 \\
&= \int_{\frac{\beta}{2}}^{\beta}\lambda_1 e^{-\lambda_1 x_1}(e^{\lambda_2 x_1-\lambda_2\beta} - e^{-\lambda_2 x_1})\mathrm{d}x_1 + \int_{\beta}^{\delta}\lambda_1 e^{-\lambda_1 x_1}(1 - e^{-\lambda_2 x_1})\mathrm{d}x_1 \\
&+ \int_d^{\infty}\lambda_1 e^{-\lambda_1 x_1}(e^{-\lambda_2 x_1+\lambda_2 d} - e^{-\lambda_2 x_1})\mathrm{d}x_1 \\
&= \int_{\frac{\beta}{2}}^{\beta}\lambda_1(e^{(\lambda_2-\lambda_1)x_1-\lambda_2\beta} - e^{-(\lambda_1+\lambda_2)x_1})\mathrm{d}x_1 + \int_{\beta}^{\delta}\lambda_1(e^{-\lambda_1 x_1} - e^{-(\lambda_1+\lambda_2)x_1})\mathrm{d}x_1 \\
&+ \int_d^{\infty}\lambda_1(e^{-(\lambda_1+\lambda_2)x_1+\lambda_2 d} - e^{-(\lambda_1+\lambda_2)x_1})\mathrm{d}x_1.
\end{aligned}
$$

Again, we assume $\lambda_1 \neq \lambda_2$. If $\lambda_1 = \lambda_2$, then the integral of the first term above would just be $\frac{\lambda_1\beta e^{-\lambda_2\beta}}{2}$. But using $\lambda_1 \neq \lambda_2$,

$$
\begin{aligned}
\mathbb{P}(A) &= \frac{\lambda_1 e^{-\lambda_2\beta}}{\lambda_2-\lambda_1}e^{(\lambda_2-\lambda_1)x_1}\Big]_{\frac{\beta}{2}}^{\beta} + \frac{\lambda_1}{\lambda_1+\lambda_2}e^{-(\lambda_1+\lambda_2)x_1}\Big]_{\frac{\beta}{2}}^{\beta} - e^{-\lambda_1 x_1}\Big]_{\beta}^{\delta} + \frac{\lambda_1}{\lambda_1+\lambda_2}e^{-(\lambda_1+\lambda_2)x_1}\Big]_{\beta}^{\delta} \\
&- \frac{\lambda_1 e^{\lambda_2 d}}{\lambda_1+\lambda_2}e^{-(\lambda_1+\lambda_1)x_1}\Big]_d^{\infty} + \frac{\lambda_1}{\lambda_1+\lambda_2}e^{-(\lambda_1+\lambda_2)x_1}\Big]_d^{\infty} \\
&= \frac{\lambda_1 e^{-\lambda_2\beta}}{\lambda_2-\lambda_1}(e^{(\lambda_2-\lambda_1)\beta} - e^{(\lambda_2-\lambda_1)\frac{\beta}{2}}) + \frac{\lambda_1}{\lambda_1+\lambda_2}(e^{-(\lambda_1+\lambda_2)\beta} - e^{-(\lambda_1+\lambda_2)\frac{\beta}{2}}) \\
&+ \frac{\lambda_1}{\lambda_1+\lambda_2}(e^{-(\lambda_1+\lambda_2)\delta} - e^{-(\lambda_1+\lambda_2)\beta}) + \frac{\lambda_1 e^{\lambda_2 d}}{\lambda_1+\lambda_2}(e^{-(\lambda_1+\lambda_2)\delta} - 0) \\
&+ e^{-\lambda_1\beta} - e^{-\lambda_1 d} + \frac{\lambda_1}{\lambda_1+\lambda_2}(0 - e^{-(\lambda_1+\lambda_2)\delta}) \\
&= \lambda_1(\frac{e^{-\lambda_1\beta}}{\lambda_2-\lambda_1} - \frac{e^{-(\lambda_1+\lambda_2)\frac{\beta}{2}}}{\lambda_2-\lambda_1} - \frac{e^{-(\lambda_1+\lambda_2)\frac{\beta}{2}}}{\lambda_1+\lambda_2} + \frac{e^{-(\lambda_1+\lambda_2)\delta}}{\lambda_1+\lambda_2} + \frac{e^{-\lambda_1 d}}{\lambda_1+\lambda_2} - \frac{e^{-(\lambda_1+\lambda_2)\delta}}{\lambda_1+\lambda_2}) \\
&+ e^{-\lambda_1\beta} - e^{-\lambda_1 d} \\
&= \lambda_1(\frac{e^{-\lambda_1\beta}}{\lambda_2-\lambda_1} - \frac{e^{-(\lambda_1+\lambda_2)\frac{\beta}{2}}}{\lambda_2-\lambda_1} - \frac{e^{-(\lambda_1+\lambda_2)\frac{\beta}{2}}}{\lambda_1+\lambda_2} + \frac{e^{-\lambda_1 d}}{\lambda_1+\lambda_2}) + e^{-\lambda_1\beta} - e^{-\lambda_1 d} \\
&= \lambda_1(\frac{e^{-\lambda_1\beta}}{\lambda_2-\lambda_1} - \frac{2\lambda_2 e^{-(\lambda_1+\lambda_2)\frac{\beta}{2}}}{\lambda_2^2-\lambda_1^2} + \frac{e^{-\lambda_1 d}}{\lambda_1+\lambda_2}) + e^{-\lambda_1\beta} - e^{-\lambda_1 d} \\
&= \lambda_1(\frac{e^{-\lambda_1\frac{\sqrt{2}}{\zeta}\delta}}{\lambda_2-\lambda_1} - \frac{2\lambda_2 e^{-(\lambda_1+\lambda_2)\frac{\delta}{\sqrt{2}\zeta}}}{\lambda_2^2-\lambda_1^2} + \frac{e^{-\lambda_1 d}}{\lambda_1+\lambda_2}) + e^{-\lambda_1\beta} - e^{-\lambda_1 d}.
\end{aligned}
$$

If we look at $\delta < 0$, then we are integrating over the intersection of three regions that do not overlap. Thus $\mathbb{P}(A) = 0$.

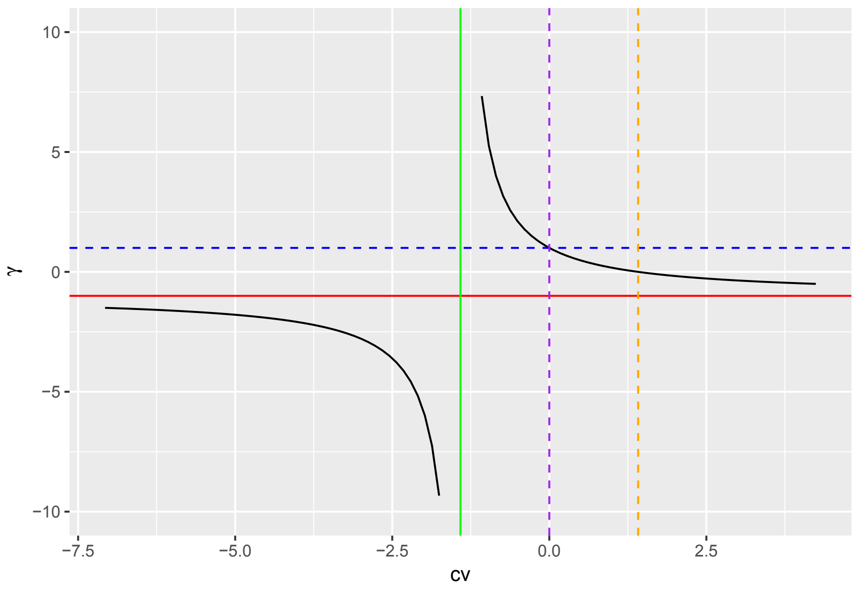


###### 2.1.1.2 Region $B$

We have $B = \{(x_1, x_2) \in \mathbb{R}^2 | x_2 \leq x_1, x_2 \geq \frac{\sqrt{2}-\zeta}{\sqrt{2}+\zeta} x_1, x_2 \leq -x_1 + \frac{\sqrt{2}}{\zeta}\delta\}$, but if we introduce the variable $\gamma = \frac{\sqrt{2}-\zeta}{\sqrt{2}+\zeta}$, we can simplify the region to $B = \{(x_1, x_2) \in \mathbb{R}^2 | x_2 \leq x_1, x_2 \geq \gamma x_1, x_2 \leq -x_1 + \beta\}$, again using $\beta = \frac{\sqrt{2}}{\zeta}\delta$. Note $\text{sgn}(\beta) = \text{sgn}(\delta)$. To understand the intersection of these three regions of $\mathbb{R}^2$, we need to understand the slope $\gamma$ as a function of $\zeta$. We only look at $\zeta \geq 0$ because $F_{\Delta,Z}(\delta, \zeta) = 0$ for $\zeta < 0$. Thus we plot $\gamma(\zeta) = \frac{\sqrt{2}-\zeta}{\sqrt{2}+\zeta}$ (shown above for region $B$).

There is a vertical asymptote at $\zeta = -\sqrt{2}$ and a horizontal asymptote at $\gamma = -1$. Since we are only interested in $\zeta \geq 0$, we see that $-1 < \gamma \leq 1$. For $0 \leq \zeta < \sqrt{2}$, we have $0 \leq \gamma \leq 1$, and for $\zeta > \sqrt{2}$, $-1 < \gamma < 0$.

Let us first look at the region where $0 \leq \zeta < \sqrt{2}$, meaning the slope of one of the lines that defines $B$ is $0 < \gamma \leq 1$, and also $\delta > 0$ (so $\beta > 0$). In this case, we have

$$\begin{aligned}
\mathbb{P}(B) &= \iint_B f_{X_1,X_2}(x_1,x_2)\mathrm{d}^2\vec{x} \\
&= \int_0^{\frac{\beta}{2}} \int_{\gamma x_1}^{x_1} \lambda_1\lambda_2 e^{-\lambda_1 x_1} e^{-\lambda_2 x_2}\mathrm{d}x_2\mathrm{d}x_1 + \int_{\frac{\beta}{2}}^{\frac{\beta}{\gamma+1}} \int_{\gamma x_1}^{-x_1+\beta} \lambda_1\lambda_2 e^{-\lambda_1 x_1} e^{-\lambda_2 x_2}\mathrm{d}x_2\mathrm{d}x_1 \\
&= \int_0^{\frac{\beta}{2}} \lambda_1 e^{-\lambda_1 x_1}(-e^{-\lambda_2 x_2})\Big]_{\gamma x_1}^{x_1}\mathrm{d}x_1 + \int_{\frac{\beta}{2}}^{\frac{\beta}{\gamma+1}} \lambda_1 e^{-\lambda_1 x_1}(-e^{-\lambda_2 x_2})\Big]_{\gamma x_1}^{-x_1+\beta}\mathrm{d}x_1 \\
&= \int_0^{\frac{\beta}{2}} \lambda_1 e^{-\lambda_1 x_1}(e^{-\lambda_2\gamma x_1} - e^{-\lambda_2 x_1})\mathrm{d}x_1 \\
&\quad + \int_{\frac{\beta}{2}}^{\frac{\beta}{\gamma+1}} \lambda_1 e^{-\lambda_1 x_1}(e^{-\lambda_2\gamma x_1} - e^{\lambda_2 x_1 - \lambda_2\beta})\mathrm{d}x_1 \\
&= \int_0^{\frac{\beta}{2}} \lambda_1(e^{-(\lambda_1+\lambda_2\gamma)x_1} - e^{-(\lambda_1+\lambda_2)x_1})\mathrm{d}x_1 \\
&\quad + \int_{\frac{\beta}{2}}^{\frac{\beta}{\gamma+1}} \lambda_1(e^{-(\lambda_1+\lambda_2\gamma)x_1} - e^{(\lambda_2-\lambda_1)x_1-\lambda_2\beta})\mathrm{d}x_1.
\end{aligned}$$

If $\lambda_1 = \lambda_2$, then the integral of the fourth term (which becomes constant) is just $\lambda_1\beta e^{-\lambda_2\beta}(\frac{1}{2} - \frac{1}{\gamma+1})$. Assuming $\lambda_1 \neq \lambda_2$,

$$\begin{aligned}
\mathbb{P}(B) &= \lambda_1(\tfrac{-1}{\lambda_1+\lambda_2\gamma}e^{-(\lambda_1+\lambda_2\gamma)x_1}\Big]_0^{\frac{\beta}{2}} + \tfrac{1}{\lambda_1+\lambda_2}e^{-(\lambda_1+\lambda_2)x_1}\Big]_0^{\frac{\beta}{2}} + \tfrac{-1}{\lambda_1+\lambda_2\gamma}e^{-(\lambda_1+\lambda_2\gamma)x_1}\Big]_{\frac{\beta}{2}}^{\frac{\beta}{\gamma+1}} \\
&\quad - \tfrac{e^{-\lambda_2\beta}}{\lambda_2-\lambda_1}e^{(\lambda_2-\lambda_1)x_1}\Big]_{\frac{\beta}{2}}^{\frac{\beta}{\gamma+1}}) \\
&= \lambda_1(\tfrac{1}{\lambda_1+\lambda_2\gamma} - \tfrac{1}{\lambda_1+\lambda_2\gamma}e^{-(\lambda_1+\lambda_2\gamma)\frac{\beta}{2}} + \tfrac{1}{\lambda_1+\lambda_2}e^{-(\lambda_1+\lambda_2)\frac{\beta}{2}} - \tfrac{1}{\lambda_1+\lambda_2} \\
&\quad - \tfrac{1}{\lambda_1+\lambda_2\gamma}e^{-(\lambda_1+\lambda_2\gamma)\frac{\beta}{\gamma+1}} + \tfrac{1}{\lambda_1+\lambda_2\gamma}e^{-(\lambda_1+\lambda_2\gamma)\frac{\beta}{2}} - \tfrac{e^{-\lambda_2\beta}}{\lambda_2-\lambda_1}e^{(\lambda_2-\lambda_1)\frac{\beta}{\gamma+1}} \\
&\quad + \tfrac{e^{-\lambda_2\beta}}{\lambda_2-\lambda_1}e^{(\lambda_2-\lambda_1)\frac{\beta}{2}}) \\
&= \lambda_1(\tfrac{\lambda_2(1-\gamma)}{(\lambda_1+\lambda_2\gamma)(\lambda_1+\lambda_2)} + \tfrac{1}{\lambda_1+\lambda_2}e^{-(\lambda_1+\lambda_2)\frac{\beta}{2}} - \tfrac{1}{\lambda_1+\lambda_2\gamma}e^{-(\lambda_1+\lambda_2\gamma)\frac{\beta}{\gamma+1}} \\
&\quad - \tfrac{1}{\lambda_2-\lambda_1}e^{-(\lambda_1+\lambda_2\gamma)\frac{\beta}{\gamma+1}} + \tfrac{1}{\lambda_2-\lambda_1}e^{-(\lambda_1+\lambda_2)\frac{\beta}{2}}) \\
&= \lambda_1\lambda_2(\tfrac{1-\gamma}{(\lambda_1+\lambda_2\gamma)(\lambda_1+\lambda_2)} + \tfrac{2}{\lambda_2^2-\lambda_1^2}e^{-(\lambda_1+\lambda_2)\frac{\beta}{2}} - \tfrac{1+\gamma}{(\lambda_1+\lambda_2\gamma)(\lambda_2-\lambda_1)}e^{-(\lambda_1+\lambda_2\gamma)\frac{\beta}{\gamma+1}}).
\end{aligned}$$

Now we look at $\zeta \geq \sqrt{2}$ and $\delta > 0$. Then $-1 < \gamma \leq 0$ and $\beta > 0$. Looking at the graph of this region, we see that we must integrate over the intersection between $B$ and Quadrant I, since $f_{X_1,X_2}(x_1,x_2) = 0$ if $x_2 < 0$. So we do not even need the line $x_2 = \gamma x_1$, and

$$\begin{aligned}
\mathbb{P}(B) &= \iint\limits_B f_{X_1,X_2}(x_1,x_2)\mathrm{d}^2\vec{x} \\
&= \int_0^{\frac{\beta}{2}} \int_{x_2}^{-x_2+\beta} \lambda_1\lambda_2 e^{-\lambda_1 x_1} e^{-\lambda_2 x_2} \mathrm{d}x_1 \mathrm{d}x_2 \\
&= \int_0^{\frac{\beta}{2}} \lambda_2 e^{-\lambda_2 x_2}(-e^{-\lambda_1 x_1})\Big]_{x_2}^{-x_2+\beta} \mathrm{d}x_2 \\
&= \int_0^{\frac{\beta}{2}} \lambda_2 e^{-\lambda_2 x_2}(e^{-\lambda_1 x_2} - e^{\lambda_1 x_2 - \lambda_1 \beta})\mathrm{d}x_2 \\
&= \int_0^{\frac{\beta}{2}} \lambda_2 (e^{-(\lambda_1+\lambda_2)x_2} - e^{(\lambda_1-\lambda_2)x_2 - \lambda_1\beta})\mathrm{d}x_2.
\end{aligned}$$

If $\lambda_1 = \lambda_2$, then the integral of the second term becomes $-\frac{\lambda_2 \beta e^{-\lambda_1 \beta}}{2}$. If $\lambda_1 \neq \lambda_2$, then

$$\begin{aligned}
\mathbb{P}(B) &= \lambda_2\Big(\frac{-1}{\lambda_1+\lambda_2} e^{-(\lambda_1+\lambda_2)x_2}\Big]_0^{\frac{\beta}{2}} - \frac{e^{-\lambda_1\beta}}{\lambda_1-\lambda_2} e^{(\lambda_1-\lambda_2)x_2}\Big]_0^{\frac{\beta}{2}}\Big) \\
&= \lambda_2\Big(\frac{1}{\lambda_1+\lambda_2} - \frac{1}{\lambda_1+\lambda_2} e^{-(\lambda_1+\lambda_2)\frac{\beta}{2}} - \frac{e^{-\lambda_1\beta}}{\lambda_1-\lambda_2} e^{(\lambda_1-\lambda_2)\frac{\beta}{2}} + \frac{e^{-\lambda_1\beta}}{\lambda_1-\lambda_2}\Big) \\
&= \lambda_2\Big(\frac{1}{\lambda_1+\lambda_2} - \frac{1}{\lambda_1+\lambda_2} e^{-(\lambda_1+\lambda_2)\frac{\beta}{2}} - \frac{1}{\lambda_1-\lambda_2} e^{-(\lambda_1+\lambda_2)\frac{\beta}{2}} + \frac{e^{-\lambda_1\beta}}{\lambda_1-\lambda_2}\Big) \\
&= \lambda_2\Big(\frac{1}{\lambda_1+\lambda_2} - \frac{2\lambda_1}{\lambda_1^2-\lambda_2^2} e^{-(\lambda_1+\lambda_2)\frac{\beta}{2}} + \frac{e^{-\lambda_2\beta}}{\lambda_1-\lambda_2}\Big) \\
&= \frac{\lambda_2}{\lambda_1^2-\lambda_2^2}(\lambda_1 - \lambda_2 - 2\lambda_1 e^{-(\lambda_1+\lambda_2)\frac{\beta}{2}} + (\lambda_1+\lambda_2)e^{-\lambda_1\beta}).
\end{aligned}$$

As is the case for region $A$, when $\delta < 0$ for region $B$, we are integrating over the intersection of three regions that do not overlap. So $\mathbb{P}(B) = 0$ when $\delta < 0$.

###### 2.1.1.3 Region $C$

Lastly, we have $C = \{(x_1,x_2) \in \mathbb{R}^2 | x_2 \leq \frac{\sqrt{2}+\zeta}{\sqrt{2}-\zeta} x_1, x_2 \geq x_1 - \min(\delta,0)\}$. Again if $\gamma = \frac{\sqrt{2}-\zeta}{\sqrt{2}+\zeta}$, then $C = \{(x_1,x_2) \in \mathbb{R}^2 | x_2 \leq \frac{1}{\gamma} x_1, x_2 \geq x_1 - \min(\delta,0)\}$. Like we did for region $B$, we need to understand $\frac{1}{\gamma}$ as a function of $\zeta$ to understand the slope of one of the lines that defines the region $C$.

We are only interested in $\zeta \geq 0$, since $F_{\Delta,Z} = 0$ when $\zeta < 0$. Looking at the figure shown on the next page (of the region $C$), we see that $1 \leq \frac{1}{\gamma} < \infty$ when $0 \leq \zeta \leq \sqrt{2}$ and $-\infty < \frac{1}{\gamma} < -1$ when $\zeta > \sqrt{2}$.

Let us first look at $0 \leq \zeta \leq \sqrt{2}$ and $\delta \leq 0$. Then $1 \leq \frac{1}{\gamma} < \infty$ and $-\min(\delta,0) = -\delta \geq 0$. We see here that the region between the two lines of

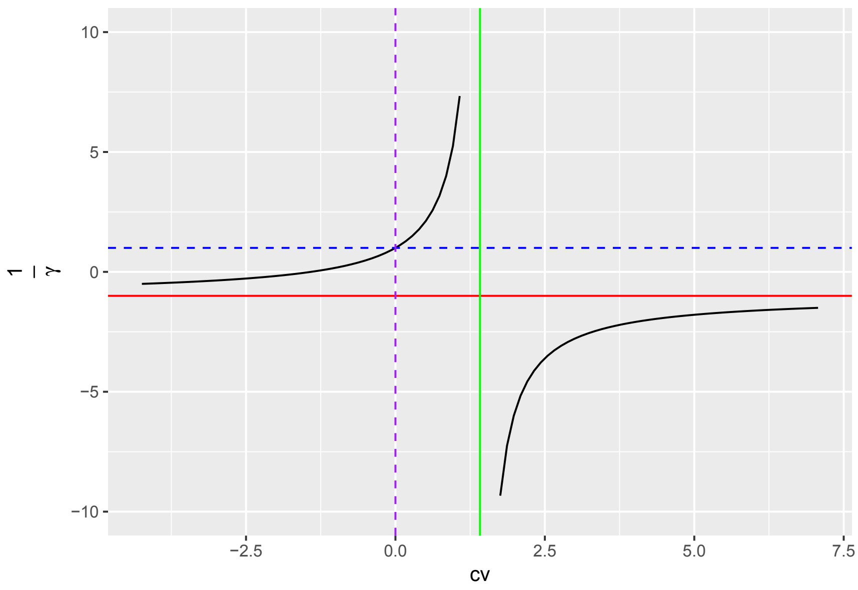


interest lies above the $x_1$-axis, where $f_{X_1,X_2}(x_1,x_2)$ is non-zero. The lines' intercept is at $(\frac{-\delta}{\frac{1}{\gamma}-1}, \frac{-\delta}{1-\gamma})$, which is in Quadrant I of $\mathbb{R}^2$. Then

$$\begin{aligned}
\mathbb{P}(C) &= \iint\limits_C f_{X_1,X_2}(x_1,x_2)\mathrm{d}^2\vec{x} \\
&= \int_{\frac{-\delta}{1-\gamma}}^{\infty} \int_{\gamma x_2}^{x_2+\delta} \lambda_1\lambda_2 e^{-\lambda_1 x_1} e^{-\lambda_2 x_2} \mathrm{d}x_1 \mathrm{d}x_2 \\
&= \int_{\frac{-\delta}{1-\gamma}}^{\infty} \lambda_2 e^{-\lambda_2 x_2} (-e^{-\lambda_1 x_1})\Big]_{\gamma x_2}^{x_2+\delta} \mathrm{d}x_2 \\
&= \int_{\frac{-\delta}{1-\gamma}}^{\infty} \lambda_2 e^{-\lambda_2 x_2} (e^{-\lambda_1 \gamma x_2} - e^{-\lambda_1 x_2 - \lambda_1 d}) \mathrm{d}x_2 \\
&= \int_{\frac{-\delta}{1-\gamma}}^{\infty} \lambda_2 (e^{-(\lambda_1\gamma+\lambda_2)x_2} - e^{-\lambda_1 d} e^{-(\lambda_1+\lambda_2)x_2}) \mathrm{d}x_2 \\
&= \lambda_2 (\frac{-1}{\lambda_1\gamma+\lambda_2} e^{-(\lambda_1\gamma+\lambda_2)x_2}\Big]_{\frac{-\delta}{1-\gamma}}^{\infty} + \frac{e^{-\lambda_1 d}}{\lambda_1+\lambda_2} e^{-(\lambda_1+\lambda_2)x_2}\Big]_{\frac{-\delta}{1-\gamma}}^{\infty}) \\
&= \lambda_2 (-0 + \frac{1}{\lambda_1\gamma+\lambda_2} e^{\frac{(\lambda_1\gamma+\lambda_2)\delta}{1-\gamma}} + 0 - \frac{e^{-\lambda_1 d}}{\lambda_1+\lambda_2} e^{\frac{(\lambda_1+\lambda_2)\delta}{1-\gamma}}) \\
&= \lambda_2 (\frac{1}{\lambda_1\gamma+\lambda_2} e^{\frac{(\lambda_1\gamma+\lambda_2)\delta}{1-\gamma}} - \frac{1}{\lambda_1+\lambda_2} e^{\frac{(\lambda_1\gamma+\lambda_2)\delta}{1-\gamma}}) \\
&= \frac{\lambda_1\lambda_2(1-\gamma)e^{\frac{(\lambda_1\gamma+\lambda_2)\delta}{1-\gamma}}}{(\lambda_1\gamma+\lambda_2)(\lambda_1+\lambda_2)}.
\end{aligned}$$

If $\delta > 0$, then $\min(\delta, 0) = 0$, and the region $C$ is defined only by $x_2 \leq \frac{1}{\gamma}x_1$ and $x_2 \geq x_1$. The intersection of the lines that define $C$ is then just $(0,0)$, and,

$$\begin{aligned}
\mathbb{P}(C) &= \iint\limits_C f_{X_1,X_2}(x_1,x_2)\mathrm{d}^2\vec{x} \\
&= \int_0^\infty \int_{x_1}^{\frac{1}{\gamma}x_1} \lambda_1\lambda_2 e^{-\lambda_1 x_1} e^{-\lambda_2 x_2}\mathrm{d}x_2\mathrm{d}x_1 \\
&= \int_0^\infty \lambda_1 e^{-\lambda_1 x_1}(-e^{-\lambda_2 x_2})\Big]_{x_1}^{\frac{1}{\gamma}x_1}\mathrm{d}x_1 \\
&= \int_0^\infty \lambda_1 e^{-\lambda_1 x_1}(e^{-\lambda_2 x_1} - e^{-\lambda_2 \frac{1}{\gamma}x_1})\mathrm{d}x_1 \\
&= \int_0^\infty \lambda_1 (e^{-(\lambda_1+\lambda_2)x_1} - e^{-(\lambda_1+\lambda_2\frac{1}{\gamma})x_1})\mathrm{d}x_1 \\
&= \lambda_1\left(\frac{-1}{\lambda_1+\lambda_2}e^{-(\lambda_1+\lambda_2)x_1}\Big]_0^\infty + \frac{1}{\lambda_1+\lambda_2\frac{1}{\gamma}}e^{-(\lambda_1+\lambda_2\frac{1}{\gamma})x_1}\Big]_0^\infty\right) \\
&= \lambda_1\left(-0 + \frac{1}{\lambda_1+\lambda_2} + 0 - \frac{1}{\lambda_1+\lambda_2\frac{1}{\gamma}}\right) \\
&= \lambda_1\left(\frac{1}{\lambda_1+\lambda_2} - \frac{\gamma}{\lambda_1\gamma+\lambda_2}\right) \\
&= \frac{\lambda_1}{(\lambda_1\gamma+\lambda_2)(\lambda_1+\lambda_2)}(\lambda_1\gamma + \lambda_2 - \lambda_1\gamma - \lambda_2\gamma) \\
&= \frac{\lambda_1\lambda_2(1-\gamma)}{(\lambda_1\gamma+\lambda_2)(\lambda_1+\lambda_2)}.
\end{aligned}$$

This is the same result as we had previously if we just let $\delta \to 0$ since we are just shifting the lower line that defines $C$ down to $(0,0)$.

If $\delta \leq 0$ and $\zeta > \sqrt{2}$, then $-\infty < \frac{1}{\gamma} < -1$ and $-\min(\delta,0) = -\delta \geq 0$. Since $f_{X_1,X_2}(x_1,x_2)$ is non-zero only in Quadrant I, based on the graph of region $C$, we do not even need the $x_2 = \frac{1}{\gamma}x_1$ line in our integration. Then

$$\begin{aligned}
\mathbb{P}(C) &= \iint\limits_C f_{X_1,X_2}(x_1,x_2)\mathrm{d}^2\vec{x} \\
&= \int_0^\infty \int_{x_1-\delta}^\infty \lambda_1\lambda_2 e^{-\lambda_1 x_1} e^{-\lambda_2 x_2}\mathrm{d}x_2\mathrm{d}x_1 \\
&= \int_0^\infty \lambda_1 e^{-\lambda_1 x_1}(-e^{-\lambda_2 x_2})\Big]_{x_1-\delta}^\infty\mathrm{d}x_1 \\
&= \int_0^\infty \lambda_1 e^{-\lambda_1 x_1}(-0 + e^{-\lambda_2 x_1 + \lambda_2 d})\mathrm{d}x_1 \\
&= \int_0^\infty \lambda_1 e^{\lambda_2 d} e^{-(\lambda_1+\lambda_2)x_1}\mathrm{d}x_1 \\
&= \frac{-\lambda_1 e^{\lambda_2 d}}{\lambda_1+\lambda_2}e^{-(\lambda_1+\lambda_2)x_1}\Big]_0^\infty \\
&= -0 + \frac{\lambda_1 e^{\lambda_2 d}}{\lambda_1+\lambda_2} \\
&= \frac{\lambda_1 e^{\lambda_2 d}}{\lambda_1+\lambda_2}.
\end{aligned}$$

Finally, if $\delta > 0$, then we replace $\delta$ with $\min(\delta,0) = 0$ in the derivation above, which would be the same as taking $\delta \to 0$ in the result. So if $\delta > 0$ and $\zeta > \sqrt{2}$, then $\mathbb{P}(C) = \frac{\lambda_1}{\lambda_1+\lambda_2}$.

###### 2.1.1.4 Joint CDF

Now we add together the contributions from the different regions to get the joint CDF for $\Delta$ and $Z$, which is defined piecewise. Again, we let $\beta = \frac{\sqrt{2}}{\zeta}\delta$ and $\gamma = \frac{\sqrt{2}-\zeta}{\sqrt{2}+\zeta}$. We also assume $\lambda_1 \neq \lambda_2$. Since $F_{\Delta,Z}(\delta,\zeta) = \mathbb{P}(A) + \mathbb{P}(B) + \mathbb{P}(C)$, we have

$$F_{\Delta,Z}(\delta,\zeta) = \begin{cases} \frac{\lambda_1\lambda_2(1-\gamma)e^{\frac{(\lambda_1\gamma+\lambda_2)\delta}{1-\gamma}}}{(\lambda_1\gamma+\lambda_2)(\lambda_1+\lambda_2)} & \text{if } \delta \le 0, 0 \le \zeta \le \sqrt{2} \\ \frac{\lambda_1 e^{\lambda_2 d}}{\lambda_1+\lambda_2} & \text{if } \delta \le 0, \zeta > \sqrt{2} \\ \frac{2\lambda_1\lambda_2}{\lambda_1^2-\lambda_2^2} e^{-(\lambda_1+\lambda_2)\frac{\delta}{\sqrt{2}\zeta}}(1-e^{\frac{(\lambda_2-\lambda_1)\delta}{2}}) \\ \quad + \lambda_1\lambda_2(\frac{1-\gamma}{(\lambda_1+\lambda_2\gamma)(\lambda_1+\lambda_2)} + \frac{2}{\lambda_2^2-\lambda_1^2}e^{-(\lambda_1+\lambda_2)\frac{\delta}{\sqrt{2}\zeta}} \\ \quad - \frac{1+\gamma}{(\lambda_1+\lambda_2\gamma)(\lambda_2-\lambda_1)}e^{-(\lambda_1+\lambda_2\gamma)\frac{\beta}{\gamma+1}}) \\ \quad + \frac{\lambda_1\lambda_2(1-\gamma)}{(\lambda_1\gamma+\lambda_2)(\lambda_1+\lambda_2)} & \text{if } \delta > 0, 0 \le \zeta \le \sqrt{2} \\ \lambda_1(\frac{e^{-\lambda_1\frac{\sqrt{2}}{\zeta}\delta}}{\lambda_2-\lambda_1} - \frac{2\lambda_2 e^{-(\lambda_1+\lambda_2)\frac{\delta}{\sqrt{2}\zeta}}}{\lambda_2^2-\lambda_1^2} + \frac{e^{-\lambda_1 d}}{\lambda_1+\lambda_2}) \\ \quad + e^{-\lambda_1\frac{\sqrt{2}}{\zeta}\delta} - e^{-\lambda_1 d} + \frac{\lambda_1}{\lambda_1+\lambda_2} \\ \quad + \frac{\lambda_2}{\lambda_1^2-\lambda_2^2}(\lambda_1 - \lambda_2 - 2\lambda_1 e^{-(\lambda_1+\lambda_2)\frac{\delta}{\sqrt{2}\zeta}} + (\lambda_1+\lambda_2)e^{-\lambda_1\frac{\sqrt{2}}{\zeta}\delta}) & \text{if } \delta > 0, \zeta > \sqrt{2} \\ 0 & \text{if } \zeta < 0. \end{cases}$$

We can simplify this function for $\delta > 0$, which will make marginalizing the CDF for $\Delta$ and $Z$ easier. Beginning with $\delta > 0$ and $0 \leq \zeta \leq \sqrt{2}$, we compute

$$
\begin{aligned}
F_{\Delta,Z}(\delta,\zeta) &= \tfrac{2\lambda_1\lambda_2}{\lambda_1^2-\lambda_2^2} e^{-(\lambda_1+\lambda_2)\frac{\delta}{\sqrt{2}\zeta}}(1-e^{\frac{(\lambda_2-\lambda_1)\delta}{2}}) \\
&\quad + \lambda_1\lambda_2\big(\tfrac{1-\gamma}{(\lambda_1+\lambda_2\gamma)(\lambda_1+\lambda_2)} + \tfrac{2}{\lambda_2^2-\lambda_1^2} e^{-(\lambda_1+\lambda_2)\frac{\delta}{\sqrt{2}\zeta}} - \tfrac{1+\gamma}{(\lambda_1+\lambda_2\gamma)(\lambda_2-\lambda_1)} e^{-(\lambda_1+\lambda_2\gamma)\frac{\beta}{\gamma+1}}\big) \\
&\quad + \tfrac{\lambda_1\lambda_2(1-\gamma)}{(\lambda_1\gamma+\lambda_2)(\lambda_1+\lambda_2)} \\
&= \tfrac{2\lambda_1\lambda_2}{\lambda_1^2-\lambda_2^2} e^{-(\lambda_1+\lambda_2)\frac{\delta}{\sqrt{2}\zeta}} - \tfrac{2\lambda_1\lambda_2}{\lambda_1^2-\lambda_2^2} e^{-(\lambda_1+\lambda_2)\frac{\delta}{\sqrt{2}\zeta}+\frac{(\lambda_2-\lambda_1)\delta}{2}} \\
&\quad + \tfrac{\lambda_1\lambda_2(1-\gamma)}{(\lambda_1+\lambda_2\gamma)(\lambda_1+\lambda_2)} - \tfrac{2\lambda_1\lambda_2}{\lambda_1^2-\lambda_2^2} e^{-(\lambda_1+\lambda_2)\frac{\delta}{\sqrt{2}\zeta}} - \tfrac{\lambda_1\lambda_2(1+\gamma)}{(\lambda_1+\lambda_2\gamma)(\lambda_2-\lambda_1)} e^{-(\lambda_1+\lambda_2\gamma)\frac{\beta}{\gamma+1}} \\
&\quad + \tfrac{\lambda_1\lambda_2(1-\gamma)}{(\lambda_1\gamma+\lambda_2)(\lambda_1+\lambda_2)} \\
&= -\tfrac{2\lambda_1\lambda_2}{\lambda_1^2-\lambda_2^2} e^{(-\frac{\lambda_1}{\sqrt{2}\zeta}-\frac{\lambda_2}{\sqrt{2}\zeta}+\frac{\lambda_2}{2}-\frac{\lambda_1}{2})\delta} - \tfrac{\lambda_1\lambda_2(1+\gamma)}{(\lambda_1+\lambda_2\gamma)(\lambda_2-\lambda_1)} e^{-(\lambda_1+\lambda_2\gamma)\frac{\beta}{\gamma+1}} \\
&\quad + \tfrac{\lambda_1\lambda_2(1-\gamma)}{\lambda_1+\lambda_2}\big(\tfrac{1}{\lambda_1+\lambda_2\gamma} + \tfrac{1}{\lambda_1\gamma+\lambda_2}\big) \\
&= -\tfrac{2\lambda_1\lambda_2}{\lambda_1^2-\lambda_2^2} e^{(-(\frac{1}{2}+\frac{1}{\sqrt{2}\zeta})\lambda_1+(\frac{1}{2}-\frac{1}{\sqrt{2}\zeta})\lambda_2)\delta} - \tfrac{\lambda_1\lambda_2(1+\gamma)}{(\lambda_1+\lambda_2\gamma)(\lambda_2-\lambda_1)} e^{-(\lambda_1+\lambda_2\gamma)\frac{1}{\gamma+1}\frac{\sqrt{2}}{\zeta}\delta} \\
&\quad + \tfrac{\lambda_1\lambda_2(1-\gamma)}{\lambda_1+\lambda_2}\big(\tfrac{\lambda_1\gamma+\lambda_2+\lambda_1+\lambda_2\gamma}{(\lambda_1+\lambda_2\gamma)(\lambda_1\gamma+\lambda_2)}\big) \\
&= -\tfrac{2\lambda_1\lambda_2}{\lambda_1^2-\lambda_2^2} e^{(-(1+\frac{\sqrt{2}}{\zeta})\lambda_1+(1-\frac{\sqrt{2}}{\zeta})\lambda_2)\frac{\delta}{2}} - \tfrac{\lambda_1\lambda_2(1+\gamma)}{(\lambda_1+\lambda_2\gamma)(\lambda_2-\lambda_1)} e^{-\frac{(\lambda_1+\lambda_2\gamma)\sqrt{2}\delta}{\zeta(\gamma+1)}} \\
&\quad + \tfrac{\lambda_1\lambda_2(1-\gamma)}{\lambda_1+\lambda_2}\big(\tfrac{(1+\gamma)\lambda_1+(1+\gamma)\lambda_2}{(\lambda_1+\lambda_2\gamma)(\lambda_1\gamma+\lambda_2)}\big) \\
&= -\tfrac{2\lambda_1\lambda_2}{\lambda_1^2-\lambda_2^2} e^{-((1+\frac{\sqrt{2}}{\zeta})\lambda_1+(\frac{\sqrt{2}}{\zeta}-1)\lambda_2)\frac{\delta}{2}} - \tfrac{\lambda_1\lambda_2(1+\gamma)}{(\lambda_1+\lambda_2\gamma)(\lambda_2-\lambda_1)} e^{-\frac{(\lambda_1+\lambda_2\gamma)\sqrt{2}\delta}{\zeta(\gamma+1)}} \\
&\quad + \tfrac{\lambda_1\lambda_2(1-\gamma)}{\lambda_1+\lambda_2}\big(\tfrac{(1+\gamma)(\lambda_1+\lambda_2)}{(\lambda_1+\lambda_2\gamma)(\lambda_1\gamma+\lambda_2)}\big) \\
&= -\tfrac{2\lambda_1\lambda_2}{\lambda_1^2-\lambda_2^2} e^{-((1+\frac{\sqrt{2}}{\zeta})\lambda_1+(\frac{\sqrt{2}}{\zeta}-1)\lambda_2)\frac{\delta}{2}} - \tfrac{\lambda_1\lambda_2(1+\gamma)}{(\lambda_1+\lambda_2\gamma)(\lambda_2-\lambda_1)} e^{-\frac{(\lambda_1+\lambda_2\gamma)\sqrt{2}\delta}{\zeta(\gamma+1)}} \\
&\quad + \tfrac{\lambda_1\lambda_2(1-\gamma^2)}{(\lambda_1+\lambda_2\gamma)(\lambda_1\gamma+\lambda_2)}.
\end{aligned}
$$

For $\delta > 0$ and $\zeta > \sqrt{2}$, we get

$$
\begin{aligned}
F_{\Delta,Z}(\delta,\zeta) &= \lambda_1\left(\frac{e^{-\lambda_1\frac{\sqrt{2}}{\zeta}\delta}}{\lambda_2-\lambda_1} - \frac{2\lambda_2 e^{-(\lambda_1+\lambda_2)\frac{\delta}{\sqrt{2}\zeta}}}{\lambda_2^2-\lambda_1^2} + \frac{e^{-\lambda_1 d}}{\lambda_1+\lambda_2}\right) + e^{-\lambda_1\frac{\sqrt{2}}{\zeta}\delta} - e^{-\lambda_1 d} \\
&\quad + \frac{\lambda_2}{\lambda_1^2-\lambda_2^2}\left(\lambda_1-\lambda_2-2\lambda_1 e^{-(\lambda_1+\lambda_2)\frac{\delta}{\sqrt{2}\zeta}} + (\lambda_1+\lambda_2)e^{-\lambda_1\frac{\sqrt{2}}{\zeta}\delta}\right) + \frac{\lambda_1}{\lambda_1+\lambda_2} \\
&= \frac{\lambda_1 e^{-\lambda_1\frac{\sqrt{2}}{\zeta}\delta}}{\lambda_2-\lambda_1} - \frac{2\lambda_1\lambda_2 e^{-(\lambda_1+\lambda_2)\frac{\delta}{\sqrt{2}\zeta}}}{\lambda_2^2-\lambda_1^2} + \frac{\lambda_1 e^{-\lambda_1 d}}{\lambda_1+\lambda_2} + e^{-\lambda_1\frac{\sqrt{2}}{\zeta}\delta} - e^{-\lambda_1 d} \\
&\quad + \frac{\lambda_2(\lambda_1-\lambda_2)}{\lambda_1^2-\lambda_2^2} - \frac{2\lambda_1\lambda_2 e^{-(\lambda_1+\lambda_2)\frac{\delta}{\sqrt{2}\zeta}}}{\lambda_1^2-\lambda_2^2} + \frac{\lambda_2(\lambda_1+\lambda_2)e^{-\lambda_1\frac{\sqrt{2}}{\zeta}\delta}}{\lambda_1^2-\lambda_2^2} + \frac{\lambda_1}{\lambda_1+\lambda_2} \\
&= \frac{\lambda_1 e^{-\lambda_1\frac{\sqrt{2}}{\zeta}\delta}}{\lambda_2-\lambda_1} + \frac{2\lambda_1\lambda_2 e^{-(\lambda_1+\lambda_2)\frac{\delta}{\sqrt{2}\zeta}}}{\lambda_1^2-\lambda_2^2} + \left(\frac{\lambda_1}{\lambda_1+\lambda_2} - 1\right)e^{-\lambda_1 d} + e^{-\lambda_1\frac{\sqrt{2}}{\zeta}\delta} \\
&\quad + \frac{\lambda_2}{\lambda_1+\lambda_2} - \frac{2\lambda_1\lambda_2 e^{-(\lambda_1+\lambda_2)\frac{\delta}{\sqrt{2}\zeta}}}{\lambda_1^2-\lambda_2^2} + \frac{\lambda_2 e^{-\lambda_1\frac{\sqrt{2}}{\zeta}\delta}}{\lambda_1-\lambda_2} + \frac{\lambda_1}{\lambda_1+\lambda_2} \\
&= -\frac{\lambda_1 e^{-\lambda_1\frac{\sqrt{2}}{\zeta}\delta}}{\lambda_1-\lambda_2} + \frac{\lambda_1-\lambda_1-\lambda_2}{\lambda_1+\lambda_2}e^{-\lambda_1 d} + e^{-\lambda_1\frac{\sqrt{2}}{\zeta}\delta} + \frac{\lambda_1+\lambda_2}{\lambda_1+\lambda_2} + \frac{\lambda_2 e^{-\lambda_1\frac{\sqrt{2}}{\zeta}\delta}}{\lambda_1-\lambda_2} \\
&= \frac{(\lambda_2-\lambda_1)e^{-\lambda_1\frac{\sqrt{2}}{\zeta}\delta}}{\lambda_1-\lambda_2} - \frac{\lambda_2}{\lambda_1+\lambda_2}e^{-\lambda_1 d} + e^{-\lambda_1\frac{\sqrt{2}}{\zeta}\delta} + 1 \\
&= -e^{-\lambda_1\frac{\sqrt{2}}{\zeta}\delta} - \frac{\lambda_2}{\lambda_1+\lambda_2}e^{-\lambda_1 d} + e^{-\lambda_1\frac{\sqrt{2}}{\zeta}\delta} + 1 \\
&= 1 - \frac{\lambda_2}{\lambda_1+\lambda_2}e^{-\lambda_1 d}.
\end{aligned}
$$

So, overall,

$$
F_{\Delta,Z}(\delta,\zeta) = \begin{cases}
\frac{\lambda_1\lambda_2(1-\gamma)e^{\frac{(\lambda_1\gamma+\lambda_2)\delta}{1-\gamma}}}{(\lambda_1\gamma+\lambda_2)(\lambda_1+\lambda_2)} & \text{if } \delta \le 0, 0 \le \zeta \le \sqrt{2} \\
\frac{\lambda_1 e^{\lambda_2 d}}{\lambda_1+\lambda_2} & \text{if } \delta \le 0, \zeta > \sqrt{2} \\
-\frac{2\lambda_1\lambda_2}{\lambda_1^2-\lambda_2^2}e^{-((1+\frac{\sqrt{2}}{\zeta})\lambda_1+(\frac{\sqrt{2}}{\zeta}-1)\lambda_2)\frac{\delta}{2}} \\
\quad -\frac{\lambda_1\lambda_2(1+\gamma)}{(\lambda_1+\lambda_2\gamma)(\lambda_2-\lambda_1)}e^{-\frac{(\lambda_1+\lambda_2\gamma)\sqrt{2}\delta}{\zeta(\gamma+1)}} \\
\quad + \frac{\lambda_1\lambda_2(1-\gamma^2)}{(\lambda_1+\lambda_2\gamma)(\lambda_1\gamma+\lambda_2)} & \text{if } \delta > 0, 0 \le \zeta \le \sqrt{2} \\
1 - \frac{\lambda_2}{\lambda_1+\lambda_2}e^{-\lambda_1 d} & \text{if } \delta > 0, \zeta > \sqrt{2} \\
0 & \text{if } \zeta < 0.
\end{cases}
$$

##### 2.1.2 Symmetric case

Thus far, we assumed that $\lambda_1 \neq \lambda_2$, meaning $X_1$ and $X_2$ are independent but not identically distributed. If instead $\lambda_1 = \lambda_2$, we need to change some of the integrals we computed.

###### 2.1.2.1 Region $A$ for $\delta > 0,\ 0 \le \zeta \le \sqrt{2}$

We can start with the same derivation as above, but we need to stop and change directions when we get to the expression with $e^{(\lambda_1-\lambda_2)x_2}$:

$$\begin{aligned}
\mathbb{P}(A) &= \textstyle\int_{\frac{\beta-\delta}{2}}^{\frac{\beta}{2}} \lambda_2 e^{-\lambda_1\beta} e^{(\lambda_1-\lambda_2)x_2} \mathrm{d}x_2 - \int_{\frac{\beta-\delta}{2}}^{\frac{\beta}{2}} \lambda_2 e^{-\lambda_1 d} e^{-(\lambda_1+\lambda_2)x_2} \mathrm{d}x_2 \\
&\quad + \textstyle\int_{\frac{\beta}{2}}^{\infty} \lambda_2 (1-e^{-\lambda_1 d}) e^{-(\lambda_1+\lambda_2)x_2} \mathrm{d}x_2 \\
&= \textstyle\int_{\frac{\beta-\delta}{2}}^{\frac{\beta}{2}} \lambda e^{-\lambda\beta} e^{(\lambda-\lambda)x_2} \mathrm{d}x_2 - \int_{\frac{\beta-\delta}{2}}^{\frac{\beta}{2}} \lambda e^{-\lambda\delta} e^{-(\lambda+\lambda)x_2} \mathrm{d}x_2 \\
&\quad + \textstyle\int_{\frac{\beta}{2}}^{\infty} \lambda (1-e^{-\lambda\delta}) e^{-(\lambda+\lambda)x_2} \mathrm{d}x_2 \\
&= \textstyle\int_{\frac{\beta-\delta}{2}}^{\frac{\beta}{2}} \lambda e^{-\lambda\beta} \mathrm{d}x_2 - \int_{\frac{\beta-\delta}{2}}^{\frac{\beta}{2}} \lambda e^{-\lambda\delta} e^{-2\lambda x_2} \mathrm{d}x_2 + \int_{\frac{\beta}{2}}^{\infty} \lambda (1-e^{-\lambda\delta}) e^{-2\lambda x_2} \mathrm{d}x_2 \\
&= \lambda e^{-\lambda\beta} x_2 \Big]_{\frac{\beta-\delta}{2}}^{\frac{\beta}{2}} + \tfrac{1}{2} e^{-\lambda\delta} e^{-2\lambda x_2} \Big]_{\frac{\beta-\delta}{2}}^{\frac{\beta}{2}} - \tfrac{1}{2}(1-e^{-\lambda\delta}) e^{-2\lambda x_2} \Big]_{\frac{\beta}{2}}^{\infty} \\
&= \lambda e^{-\lambda\beta}(\tfrac{\beta}{2} - \tfrac{\beta-\delta}{2}) + \tfrac{1}{2} e^{-\lambda\delta}(e^{-\lambda\beta} - e^{-\lambda\beta+\lambda\delta}) - \tfrac{1}{2}(1-e^{-\lambda\delta})(0-e^{-\lambda\beta}) \\
&= \tfrac{\lambda e^{-\lambda\beta}\delta}{2} + \tfrac{1}{2} e^{-\lambda\beta-\lambda\delta} - \tfrac{1}{2} e^{-\lambda\beta} + \tfrac{1}{2} e^{-\lambda\beta} - \tfrac{1}{2} e^{-\lambda\beta-\lambda\delta} \\
&= \tfrac{\lambda e^{-\lambda\beta}\delta}{2}.
\end{aligned}$$

###### 2.1.2.2 Region $B$ for $\delta > 0,\ 0 \leq \zeta \leq \sqrt{2}$

Again, we change directions in our derivation when we get to an expression with $e^{(\lambda_1-\lambda_2)x_1}$:

$$
\begin{aligned}
\mathbb{P}(B) &= \textstyle\int_0^{\frac{\beta}{2}} \lambda_1(e^{-(\lambda_1+\lambda_2\gamma)x_1} - e^{-(\lambda_1+\lambda_2)x_1})\mathrm{d}x_1 \\
&\quad + \textstyle\int_{\frac{\beta}{2}}^{\frac{\beta}{\gamma+1}} \lambda_1(e^{-(\lambda_1+\lambda_2\gamma)x_1} - e^{(\lambda_2-\lambda_1)x_1-\lambda_2\beta})\mathrm{d}x_1 \\
&= \textstyle\int_0^{\frac{\beta}{2}} \lambda(e^{-(\lambda+\lambda\gamma)x_1} - e^{-(\lambda+\lambda)x_1})\mathrm{d}x_1 + \int_{\frac{\beta}{2}}^{\frac{\beta}{\gamma+1}} \lambda(e^{-(\lambda+\lambda\gamma)x_1} - e^{(\lambda-\lambda)x_1-\lambda\beta})\mathrm{d}x_1 \\
&= \textstyle\int_0^{\frac{\beta}{2}} \lambda(e^{-\lambda(\gamma+1)x_1} - e^{-2\lambda x_1})\mathrm{d}x_1 + \int_{\frac{\beta}{2}}^{\frac{\beta}{\gamma+1}} \lambda(e^{-\lambda(\gamma+1)x_1} - e^{-\lambda\beta})\mathrm{d}x_1 \\
&= -\tfrac{1}{\gamma+1}e^{-\lambda(\gamma+1)x_1}\Big]_0^{\frac{\beta}{2}} + \tfrac{1}{2}e^{-2\lambda x_1}\Big]_0^{\frac{\beta}{2}} - \tfrac{1}{\gamma+1}e^{-\lambda(\gamma+1)x_1}\Big]_{\frac{\beta}{2}}^{\frac{\beta}{\gamma+1}} - \lambda e^{-\lambda\beta}x_1\Big]_{\frac{\beta}{2}}^{\frac{\beta}{\gamma+1}} \\
&= -\tfrac{1}{\gamma+1}(e^{-\lambda(\gamma+1)\frac{\beta}{2}} - 1) + \tfrac{1}{2}e^{-\lambda\beta} - \tfrac{1}{2} - \tfrac{1}{\gamma+1}(e^{-\lambda\beta} - e^{-\lambda(\gamma+1)\frac{\beta}{2}}) \\
&\quad -\lambda e^{-\lambda\beta}(\tfrac{\beta}{\gamma+1} - \tfrac{\beta}{2}) \\
&= -\tfrac{1}{\gamma+1}e^{-\lambda(\gamma+1)\frac{\beta}{2}} + \tfrac{1}{\gamma+1} + \tfrac{1}{2}e^{-\lambda\beta} - \tfrac{1}{2} - \tfrac{1}{\gamma+1}e^{-\lambda\beta} + \tfrac{1}{\gamma+1}e^{-\lambda(\gamma+1)\frac{\beta}{2}} \\
&\quad +\lambda\beta e^{-\lambda\beta}(\tfrac{1}{2} - \tfrac{1}{\gamma+1}) \\
&= -(\tfrac{1}{2} - \tfrac{1}{\gamma+1}) + e^{-\lambda\beta}(\tfrac{1}{2} - \tfrac{1}{\gamma+1}) + \lambda\beta e^{-\lambda\beta}(\tfrac{1}{2} - \tfrac{1}{\gamma+1}) \\
&= (\tfrac{1}{2} - \tfrac{1}{\gamma+1})(-1 + e^{-\lambda\beta} + \lambda\beta e^{-\lambda\beta}).
\end{aligned}
$$

###### 2.1.2.3 $F_{\Delta,Z}(\delta,\zeta)$ for $\delta > 0,\ 0 \leq \zeta \leq \sqrt{2}$

Notice that we do not have to make any adjustments to region $C$, where $\mathbb{P}(C) = \frac{\lambda_1\lambda_2(1-\gamma)}{(\lambda_1\gamma+\lambda_2)(\lambda_1+\lambda_2)} = \frac{\lambda^2(1-\gamma)}{(\lambda\gamma+\lambda)(\lambda+\lambda)} = \frac{\lambda^2(1-\gamma)}{\lambda(\gamma+1)2\lambda} = \frac{1-\gamma}{2(1+\gamma)}$. So, in total,

$$
\begin{aligned}
F_{\Delta,Z}(\delta,\zeta) &= \mathbb{P}(A) + \mathbb{P}(B) + \mathbb{P}(C) \\
&= \tfrac{\lambda e^{-\lambda\beta}\delta}{2} + (\tfrac{1}{2} - \tfrac{1}{\gamma+1})(-1 + e^{-\lambda\beta} + \lambda\beta e^{-\lambda\beta}) + \tfrac{1-\gamma}{2(1+\gamma)}.
\end{aligned}
$$

###### 2.1.2.4 Region $A$ for $\delta > 0$, $\zeta > \sqrt{2}$

Now we move on to $\delta > 0$ and $\zeta > \sqrt{2}$, and start with Region $A$. Continuing like before from the first expression to include $e^{-(\lambda_1+\lambda_2)x_1}$:

$$
\begin{aligned}
\mathbb{P}(A) &= \textstyle\int_{\frac{\beta}{2}}^{\beta} \lambda_1(e^{(\lambda_2-\lambda_1)x_1-\lambda_2\beta} - e^{-(\lambda_1+\lambda_2)x_1})\mathrm{d}x_1 \\
&\quad + \textstyle\int_{\beta}^{\delta} \lambda_1(e^{-\lambda_1 x_1} - e^{-(\lambda_1+\lambda_2)x_1})\mathrm{d}x_1 \\
&\quad + \textstyle\int_{d}^{\infty} \lambda_1(e^{-(\lambda_1+\lambda_2)x_1+\lambda_2 d} - e^{-(\lambda_1+\lambda_2)x_1})\mathrm{d}x_1 \\
&= \textstyle\int_{\frac{\beta}{2}}^{\beta} \lambda(e^{(\lambda-\lambda)x_1-\lambda\beta} - e^{-(\lambda+\lambda)x_1})\mathrm{d}x_1 + \int_{\beta}^{\delta} \lambda(e^{-\lambda x_1} - e^{-(\lambda+\lambda)x_1})\mathrm{d}x_1 \\
&\quad + \textstyle\int_{d}^{\infty} \lambda(e^{-(\lambda+\lambda)x_1+\lambda\delta} - e^{-(\lambda+\lambda)x_1})\mathrm{d}x_1 \\
&= \textstyle\int_{\frac{\beta}{2}}^{\beta} \lambda(e^{-\lambda\beta} - e^{-2\lambda x_1})\mathrm{d}x_1 + \int_{\beta}^{\delta} \lambda(e^{-\lambda x_1} - e^{-2\lambda x_1})\mathrm{d}x_1 \\
&\quad + \textstyle\int_{d}^{\infty} \lambda(e^{-2\lambda x_1+\lambda\delta} - e^{-2\lambda x_1})\mathrm{d}x_1 \\
&= \lambda e^{-\lambda\beta} x_1\Big]_{\frac{\beta}{2}}^{\beta} + \tfrac{1}{2}e^{-2\lambda x_1}\Big]_{\frac{\beta}{2}}^{\beta} - e^{-\lambda x_1}\Big]_{\beta}^{\delta} + \tfrac{1}{2}e^{-2\lambda x_1}\Big]_{\beta}^{\delta} \\
&\quad -\tfrac{1}{2}e^{\lambda\delta}e^{-2\lambda x_1}\Big]_{d}^{\infty} + \tfrac{1}{2}e^{-2\lambda x_1}\Big]_{d}^{\infty} \\
&= \lambda e^{-\lambda\beta}(\beta - \tfrac{\beta}{2}) + \tfrac{1}{2}(e^{-2\lambda\beta} - e^{-\lambda\beta}) - (e^{-\lambda\delta} - e^{-\lambda\beta}) \\
&\quad + \tfrac{1}{2}(e^{-2\lambda\delta} - e^{-2\lambda\beta}) - \tfrac{1}{2}e^{\lambda\delta}(0 - e^{-2\lambda\delta}) + \tfrac{1}{2}(0 - e^{-2\lambda\delta}) \\
&= \tfrac{\lambda e^{-\lambda\beta}\beta}{2} + \tfrac{1}{2}e^{-2\lambda\beta} - \tfrac{1}{2}e^{-\lambda\beta} - e^{-\lambda\delta} + e^{-\lambda\beta} \\
&\quad + \tfrac{1}{2}e^{-2\lambda\delta} - \tfrac{1}{2}e^{-2\lambda\beta} + \tfrac{1}{2}e^{-\lambda\delta} - \tfrac{1}{2}e^{-2\lambda\delta} \\
&= \tfrac{\lambda e^{-\lambda\beta}\beta}{2} - \tfrac{1}{2}e^{-\lambda\delta} + \tfrac{1}{2}e^{-\lambda\beta}.
\end{aligned}
$$

###### 2.1.2.5 Region $B$ for $\delta > 0$, $\zeta > \sqrt{2}$

Lastly, we have Region $B$ for $\delta > 0$ and $\zeta > \sqrt{2}$. We find the first expression with $e^{(\lambda_1-\lambda_2)x_2}$ and continue from there:

$$
\begin{aligned}
\mathbb{P}(B) &= \textstyle\int_{0}^{\frac{\beta}{2}} \lambda_2(e^{-(\lambda_1+\lambda_2)x_2} - e^{(\lambda_1-\lambda_2)x_2-\lambda_1\beta})\mathrm{d}x_2 \\
&= \textstyle\int_{0}^{\frac{\beta}{2}} \lambda(e^{-(\lambda+\lambda)x_2} - e^{(\lambda-\lambda)x_2-\lambda\beta})\mathrm{d}x_2 \\
&= \textstyle\int_{0}^{\frac{\beta}{2}} \lambda(e^{-2\lambda x_2} - e^{-\lambda\beta})\mathrm{d}x_2 \\
&= -\tfrac{1}{2}e^{-2\lambda x_2}\Big]_{0}^{\frac{\beta}{2}} - \lambda e^{-\lambda\beta}x_2\Big]_{0}^{\frac{\beta}{2}} \\
&= -\tfrac{1}{2}(e^{-\lambda\beta} - 1) - \lambda e^{-\lambda\beta}(\tfrac{\beta}{2} - 0) \\
&= -\tfrac{1}{2}e^{-\lambda\beta} + \tfrac{1}{2} - \tfrac{\lambda e^{-\lambda\beta}\beta}{2}.
\end{aligned}
$$

###### 2.1.2.6 $F_{\Delta,Z}(\delta,\zeta)$ for $\delta > 0$, $\zeta > \sqrt{2}$

Again, we do not have to make any adjustments to region $C$, where $\mathbb{P}(C) = \frac{\lambda_1}{\lambda_1+\lambda_2} = \frac{\lambda}{\lambda+\lambda} = \frac{1}{2}$. So, we have in total for $\delta > 0$ and $\zeta > \sqrt{2}$,

$$\begin{aligned} F_{\Delta,Z}(\delta,\zeta) &= \mathbb{P}(A)+\mathbb{P}(B)+\mathbb{P}(C) \\ &= \frac{\lambda e^{-\lambda\beta}\beta}{2} - \frac{1}{2}e^{-\lambda\delta} + \frac{1}{2}e^{-\lambda\beta} - \frac{1}{2}e^{-\lambda\beta} + \frac{1}{2} - \frac{\lambda e^{-\lambda\beta}\beta}{2} + \frac{1}{2} \\ &= 1 - \frac{1}{2}e^{-\lambda\delta}. \end{aligned}$$

###### 2.1.2.7 Overall joint CDF

None of the calculations remaining in the original derivation change if $\delta < 0$, so we can just simplify them with $\lambda_1 = \lambda_2 = \lambda$. For $0 \le \zeta \le \sqrt{2}$,

$$\begin{aligned} F_{\Delta,Z}(\delta,\zeta) &= \frac{\lambda_1\lambda_2(1-\gamma)e^{\frac{(\lambda_1\gamma+\lambda_2)\delta}{1-\gamma}}}{(\lambda_1\gamma+\lambda_2)(\lambda_1+\lambda_2)} \\ &= \frac{\lambda\lambda(1-\gamma)e^{\frac{(\lambda\gamma+\lambda)\delta}{1-\gamma}}}{(\lambda\gamma+\lambda)(\lambda+\lambda)} \\ &= \frac{\lambda^2(1-\gamma)e^{\frac{\lambda(1+\gamma)\delta}{1-\gamma}}}{\lambda(1+\gamma)2\lambda} \\ &= \frac{1-\gamma}{2(1+\gamma)}e^{\frac{\lambda(1+\gamma)\delta}{1-\gamma}}, \end{aligned}$$

and for $\zeta > \sqrt{2}$,

$$\begin{aligned} F_{\Delta,Z}(\delta,\zeta) &= \frac{\lambda_1 e^{\lambda_2 d}}{\lambda_1+\lambda_2} \\ &= \frac{\lambda e^{\lambda\delta}}{\lambda+\lambda} \\ &= \frac{1}{2}e^{\lambda\delta}. \end{aligned}$$

As always, $F_{\Delta,Z}(\delta,\zeta) = 0$ for $\zeta < 0$. Then we finally get

$$F_{\Delta,Z}(\delta,\zeta) = \begin{cases} \frac{1-\gamma}{2(1+\gamma)}e^{\frac{\lambda(1+\gamma)\delta}{1-\gamma}} & \text{if } \delta \le 0, 0 \le \zeta \le \sqrt{2} \\ \frac{1}{2}e^{\lambda\delta} & \text{if } \delta \le 0, \zeta > \sqrt{2} \\ (\frac{1}{2} - \frac{1}{\gamma+1})(-1 + e^{-\lambda\beta} + \lambda\beta e^{-\lambda\beta}) & \\ \quad + \frac{\lambda e^{-\lambda\beta}\delta}{2} + \frac{1-\gamma}{2(1+\gamma)} & \text{if } \delta > 0, 0 \le \zeta \le \sqrt{2} \\ 1 - \frac{1}{2}e^{-\lambda\delta} & \text{if } \delta > 0, \zeta > \sqrt{2} \\ 0 & \text{if } \zeta < 0. \end{cases}$$

###### 2.1.2.8 Marginal CDF for $Z$

It may seem strange that for $\zeta > \sqrt{2}$, $F_{\Delta,Z}(\delta,\zeta)$ does not depend on $\zeta$, but this makes sense when we consider the marginal CDF for $Z$. If the CDF does not depend on $Z$ here, then there is no extra probability mass gained by increasing $Z$, so we would assume that the marginal CDF, $F_Z(\zeta)$, is 1 for any value of $\zeta$ greater than $\sqrt{2}$. We can check this when we have the complete marginal CDF.

We will start with $0 \le \zeta \le \sqrt{2}$, remembering that for $\zeta < 0$, $F_{\Delta,Z}(\delta,\zeta)$ is zero, so $F_Z(\zeta)$ would be too. We also begin with the asymmetric case with $\lambda_1 \ne \lambda_2$. Looking at $F_{\Delta,Z}(\delta,\zeta)$ for $0 \le \zeta \le \sqrt{2}$ and $\delta > 0$,

$$\begin{aligned} F_{\Delta,Z}(\delta,\zeta) &= -\tfrac{2\lambda_1\lambda_2}{\lambda_1^2-\lambda_2^2} e^{-((1+\frac{\sqrt{2}}{\zeta})\lambda_1+(\frac{\sqrt{2}}{\zeta}-1)\lambda_2)\frac{\delta}{2}} - \tfrac{\lambda_1\lambda_2(1+\gamma)}{(\lambda_1+\lambda_2\gamma)(\lambda_2-\lambda_1)} e^{-\frac{(\lambda_1+\lambda_2\gamma)\sqrt{2}\delta}{\zeta(\gamma+1)}} \\ &\quad + \tfrac{\lambda_1\lambda_2(1-\gamma^2)}{(\lambda_1+\lambda_2\gamma)(\lambda_1\gamma+\lambda_2)}. \end{aligned}$$

We know that $\lambda_1 > 0$, $\lambda_2 > 0$, and $0 \le \zeta \le \sqrt{2}$. From the figure shown earlier, for $\gamma = \frac{\sqrt{2}-\zeta}{\sqrt{2}+\zeta}$, $0 \le \gamma \le 1$ for $0 \le \zeta \le \sqrt{2}$. Then all of the coefficients $\alpha$ in the $e^{-\alpha\delta}$ expressions above are positive, and

$$\lim_{\delta\to\infty} e^{-\alpha\delta} = 0.$$

Using these facts and the continutiy of $F_{\Delta,Z}(\delta,\zeta)$, for $0 \le \zeta \le \sqrt{2}$, we have

$$\begin{aligned} F_Z(\zeta) &= F_{\Delta,Z}(\infty,\zeta) \\ &= \lim_{\delta\to\infty} F_{\Delta,Z}(\delta,\zeta) \\ &= \lim_{\delta\to\infty}(-\tfrac{2\lambda_1\lambda_2}{\lambda_1^2-\lambda_2^2} e^{-((1+\frac{\sqrt{2}}{\zeta})\lambda_1+(\frac{\sqrt{2}}{\zeta}-1)\lambda_2)\frac{\delta}{2}} \\ &\quad -\tfrac{\lambda_1\lambda_2(1+\gamma)}{(\lambda_1+\lambda_2\gamma)(\lambda_2-\lambda_1)} e^{-\frac{(\lambda_1+\lambda_2\gamma)\sqrt{2}\delta}{\zeta(\gamma+1)}} + \tfrac{\lambda_1\lambda_2(1-\gamma^2)}{(\lambda_1+\lambda_2\gamma)(\lambda_1\gamma+\lambda_2)}) \\ &= -\tfrac{2\lambda_1\lambda_2}{\lambda_1^2-\lambda_2^2}(0) - \tfrac{\lambda_1\lambda_2(1+\gamma)}{(\lambda_1+\lambda_2\gamma)(\lambda_2-\lambda_1)}(0) + \tfrac{\lambda_1\lambda_2(1-\gamma^2)}{(\lambda_1+\lambda_2\gamma)(\lambda_1\gamma+\lambda_2)} \\ &= \tfrac{\lambda_1\lambda_2(1-\gamma^2)}{(\lambda_1+\lambda_2\gamma)(\lambda_1\gamma+\lambda_2)} \end{aligned}$$

Similarly, for $\zeta > \sqrt{2}$, we have $F_{\Delta,Z}(\delta,\zeta) = 1 - \frac{\lambda_2}{\lambda_1+\lambda_2} e^{-\lambda_1 d}$, and

$$\begin{aligned} F_Z(\zeta) &= F_{\Delta,Z}(\infty,\zeta) \\ &= \lim_{\delta\to\infty} F_{\Delta,Z}(\delta,\zeta) \\ &= \lim_{\delta\to\infty}(1 - \tfrac{\lambda_2}{\lambda_1+\lambda_2} e^{-\lambda_1 d}) \\ &= 1. \end{aligned}$$

It makes sense that $F_Z(\zeta) = \mathbb{P}(Z \le \zeta) = 1$ for $\zeta > \sqrt{2}$, since $1 - \mathbb{P}(Z \le \sqrt{2}) = \mathbb{P}(Z > \sqrt{2}) = \mathbb{P}(\frac{\sqrt{2}|X_1-X_2|}{X_1+X_2} > \sqrt{2}) = \mathbb{P}(|X_1-X_2| > X_1+X_2) = 0$ for two positive random variables $X_1 \sim Exp(\lambda_1)$ and $X_2 \sim Exp(\lambda_2)$. We can see that $F_Z(\zeta)$ is continuous for all of $\mathbb{R}$. It clearly is at $\zeta < 0$, $0 < \zeta < \sqrt{2}$, and $\sqrt{2} < \zeta$. We just need to check the boundaries. Since $\gamma = \frac{\sqrt{2}-\zeta}{\sqrt{2}+\zeta}$, $\gamma \to 1^-$ as $\zeta \to 0^+$ and $\gamma \to 0^+$ as $\zeta \to \sqrt{2}^{(-)}$. Then

$$\begin{aligned}
\lim_{\zeta \to 0^+} F_Z(\zeta) &= \lim_{\zeta \to 0^+} \tfrac{\lambda_1\lambda_2(1-\gamma^2)}{(\lambda_1+\lambda_2\gamma)(\lambda_1\gamma+\lambda_2)} \\
&= \lim_{\gamma \to 1^-} \tfrac{\lambda_1\lambda_2(1-\gamma^2)}{(\lambda_1+\lambda_2\gamma)(\lambda_1\gamma+\lambda_2)} \\
&= \tfrac{\lambda_1\lambda_2(1-1^2)}{(\lambda_1+\lambda_2*1)(\lambda_1*1+\lambda_2)} \\
&= 0
\end{aligned}$$

and

$$\begin{aligned}
\lim_{\zeta \to \sqrt{2}^{(-)}} F_Z(\zeta) &= \lim_{\zeta \to \sqrt{2}^{(-)}} \tfrac{\lambda_1\lambda_2(1-\gamma^2)}{(\lambda_1+\lambda_2\gamma)(\lambda_1\gamma+\lambda_2)} \\
&= \lim_{\gamma \to 0^+} \tfrac{\lambda_1\lambda_2(1-\gamma^2)}{(\lambda_1+\lambda_2\gamma)(\lambda_1\gamma+\lambda_2)} \\
&= \tfrac{\lambda_1\lambda_2(1-0^2)}{(\lambda_1+\lambda_2*0)(\lambda_1*0+\lambda_2)} \\
&= \tfrac{\lambda_1\lambda_2}{\lambda_1\lambda_2} \\
&= 1.
\end{aligned}$$

With parameters $\lambda_1 > 0$ and $\lambda_2 > 0$, using the simplification $\gamma = \frac{\sqrt{2}-\zeta}{\sqrt{2}+\zeta}$ and for independent $X_1 \sim Exp(\lambda_1)$ and $X_2 \sim Exp(\lambda_2)$, $Z = \frac{\sqrt{2}|X_1-X_2|}{X_1+X_2}$ has the CDF

$$F_Z(\zeta) = \begin{cases} 0 & \text{if } \zeta < 0 \\ \frac{\lambda_1\lambda_2(1-\gamma^2)}{(\lambda_1+\lambda_2\gamma)(\lambda_1\gamma+\lambda_2)} & \text{if } 0 \le \zeta \le \sqrt{2} \\ 1 & \text{if } \zeta > \sqrt{2}. \end{cases}$$

Even though we derived the joint CDF for $(\Delta, Z)$ by integrating different functions in the symmetric case (dropping terms of the form $e^{(\lambda_1-\lambda_2)x_2}$ or $e^{(\lambda_1-\lambda_2)x_1}$ or $e^{(\lambda_1-\lambda_2)x_2}$), the marginal CDF in the symmetric case agrees with just plugging $\lambda_1 = \lambda_2 = \lambda$ in the asymmetric marginal CDF. Since $\beta = \frac{\sqrt{2}}{\zeta}\delta$ and $\lim_{\delta\to\infty} de^{-\lambda\delta} = 0$ for $\lambda > 0$ by L'Hospital's Rule, we get $\lim_{\delta\to\infty} de^{-\lambda\beta} = 0$ and $\lim_{\delta\to\infty} \beta e^{-\lambda\beta} = 0$. Then, using the continuity of this joint CDF, the marginal

CDF for $Z$ when $0 \le \zeta \le \sqrt{2}$ is

$$\begin{aligned}
F_Z(\zeta) &= \lim_{\delta\to\infty} F_{\Delta,Z}(\delta,\zeta) \\
&= \lim_{\delta\to\infty}\left(\tfrac{\lambda e^{-\lambda\beta}\delta}{2} + (\tfrac{1}{2} - \tfrac{1}{\gamma+1})(-1 + e^{-\lambda\beta} + \lambda\beta e^{-\lambda\beta}) + \tfrac{1-\gamma}{2(1+\gamma)}\right) \\
&= 0 + (\tfrac{1}{2} - \tfrac{1}{\gamma+1})(-1+0+0) + \tfrac{1-\gamma}{2(1+\gamma)} \\
&= \tfrac{1}{\gamma+1} - \tfrac{1}{2} + \tfrac{1-\gamma}{2(1+\gamma)} \\
&= \tfrac{2-\gamma-1}{2(\gamma+1)} + \tfrac{1-\gamma}{2(1+\gamma)} \\
&= \tfrac{1-\gamma}{2(1+\gamma)} + \tfrac{1-\gamma}{2(1+\gamma)} \\
&= \tfrac{1-\gamma}{1+\gamma}.
\end{aligned}$$

This agrees with what we would have had if we just plugged $\lambda_1 = \lambda_2 = \lambda$ in the asymmetric case. We can simplify this further:

$$\begin{aligned}
F_Z(\zeta) &= \tfrac{1-\gamma}{1+\gamma} \\
&= \frac{1-\frac{\sqrt{2}-\zeta}{\sqrt{2}+\zeta}}{1+\frac{\sqrt{2}-\zeta}{\sqrt{2}+\zeta}} \\
&= \frac{\sqrt{2}+\zeta-\sqrt{2}+\zeta}{\sqrt{2}+\zeta} \Bigg/ \frac{\sqrt{2}+\zeta+\sqrt{2}-\zeta}{\sqrt{2}+\zeta} \\
&= \tfrac{2cv}{2\sqrt{2}} \\
&= \tfrac{\zeta}{\sqrt{2}}.
\end{aligned}$$

This means that when $X_1, X_2 \sim Exp(\lambda)$ are *i.i.d.*, $Z \sim Unif(0,\sqrt{2})$.

When $\zeta > \sqrt{2}$,

$$\begin{aligned}
F_Z(\zeta) &= \lim_{\delta\to\infty} F_{\Delta,Z}(\delta,\zeta) \\
&= \lim_{\delta\to\infty}\left(1 - \tfrac{1}{2}e^{-\lambda\delta}\right) \\
&= 1,
\end{aligned}$$

which makes sense given that $Z$ cannot take values greater than $\sqrt{2}$.

###### 2.1.2.9 Marginal PDF for $Z$

Since the marginal form of $Z$ is neither log-normal nor Weibull, we should understand its PDF. This would give us a way to estimate our new random variable $Z$ as a log-normal or Weibull. This estimation would necessarily be imperfect, since log-normal and Weibull distributions have support on $(0,\infty)$ and $[0,\infty)$, respectively, while $Z$'s PDF is only supported on $[0,\sqrt{2}]$.

Rather than directly finding the PDF of $Z$, we will make a transformation to make the calculations easier. Let $Y = \frac{1}{\sqrt{2}}Z = \frac{|X_1-X_2|}{X_1+X_2}$, where

$X_1 \sim Exp(\lambda_1)$ and $X_2 \sim Exp(\lambda_2)$ are independent as before. Then $Z = \sqrt{2}Y$, meaning if $F_Z(\zeta)$ is the CDF for $Z$ and $f_Z(\zeta)$ is the PDF for $Z$, then $F_Z(\zeta) = F_Y(\frac{\zeta}{\sqrt{2}})$ and $f_Z(\zeta) = \frac{1}{\sqrt{2}} f_Y(\frac{\zeta}{\sqrt{2}})$ given the CDF $F_Y(y)$ and PDF $f_Y(y)$ for $Y$, respectively. Since $\gamma(\zeta) = \frac{\sqrt{2}-\zeta}{\sqrt{2}+\zeta} = \frac{1-\frac{\zeta}{\sqrt{2}}}{1+\frac{\zeta}{\sqrt{2}}}$, we should define $z(y) = \frac{1-y}{1+y}$, so $z(\frac{\zeta}{\sqrt{2}}) = \frac{1-\frac{\zeta}{\sqrt{2}}}{1+\frac{\zeta}{\sqrt{2}}} = \gamma(\zeta)$. Then the CDF of $Y$ is just

$$F_Y(y) = \begin{cases} 0 & \text{if } y < 0 \\ \frac{\lambda_1\lambda_2(1-z^2)}{(\lambda_1+\lambda_2 z)(\lambda_1 z+\lambda_2)} & \text{if } 0 \leq y \leq 1 \\ 1 & \text{if } y > 1, \end{cases}$$

which is much easier to differentiate. Notice we have $F_Y(\frac{\zeta}{\sqrt{2}}) = F_Z(\zeta)$, which verifies the calculation. Before we differentiate, we should simplify this for $0 \leq y \leq 1$, especially in the denominator:

$$\begin{aligned} F_Y(y) &= \frac{\lambda_1\lambda_2(1-z^2)}{(\lambda_1+\lambda_2 z)(\lambda_1 z+\lambda_2)} \\ &= \frac{\lambda_1\lambda_2(1-z^2)}{\lambda_1^2 z+\lambda_1\lambda_2+\lambda_1\lambda_2 z^2+\lambda_2^2 z} \\ &= \frac{\lambda_1\lambda_2(1-z^2)}{(\lambda_1^2+\lambda_2^2)z+(1+z^2)\lambda_1\lambda_2} \\ &= \frac{\lambda_1\lambda_2(1-z^2)}{(\frac{\lambda_1}{\lambda_2}+\frac{\lambda_2}{\lambda_1})\lambda_1\lambda_2 z+(1+z^2)\lambda_1\lambda_2} \\ &= \frac{1-z^2}{z^2+\lambda_0 z+1}, \end{aligned}$$

if we let $\lambda_0 = \frac{\lambda_1}{\lambda_2} + \frac{\lambda_2}{\lambda_2}$. We also take from this that $\lambda_1\lambda_2(z^2+\lambda_0 z+1) = (\lambda_1+\lambda_2 z)(\lambda_1 z+\lambda_2)$.

Now we differentiate $F_Y(y)$ using the chain rule. Obviously for $y < 0$ and $y > 1$, $f_Y(y) = 0$. For $0 < y < 1$,

$$\begin{aligned}
f_Y(y) &= \frac{\mathrm{d}F_Y(y)}{\mathrm{d}y} \\
&= \frac{\mathrm{d}F_Y(z)}{\mathrm{d}z}\frac{\mathrm{d}z}{\mathrm{d}y} \\
&= \frac{\mathrm{d}}{\mathrm{d}z}\Big(\frac{1-z^2}{z^2+\lambda_0 z+1}\Big)\frac{\mathrm{d}}{\mathrm{d}y}\Big(\frac{1-y}{1+y}\Big) \\
&= \frac{(z^2+\lambda_0 z+1)(-2z)-(1-z^2)(2z+\lambda_0)}{(z^2+\lambda_0 z+1)^2}\frac{(1+y)(-1)-(1-y)(1)}{(1+y)^2} \\
&= \frac{-2z^3-2\lambda_0 z^2-2z-2z-\lambda_0+2z^3+\lambda_0 z^2}{(\frac{(\lambda_1+\lambda_2 z)(\lambda_1 z+\lambda_2)}{\lambda_1\lambda_2})^2}\frac{-1-y-1+y}{(1+y)^2} \\
&= \frac{(-\lambda_0 z^2-4z-\lambda_0)(\lambda_1\lambda_2)^2}{(\lambda_1+\lambda_2 z)^2(\lambda_1 z+\lambda_2)^2}\frac{-2}{(1+y)^2} \\
&= \frac{\lambda_0\lambda_1\lambda_2 z^2+4\lambda_1\lambda_2 z+\lambda_0\lambda_1\lambda_2}{(\lambda_1+\lambda_2 z)^2(\lambda_1 z+\lambda_2)^2}\frac{2\lambda_1\lambda_2}{(1+y)^2} \\
&= \frac{(\lambda_1^2+\lambda_2^2)z^2+4\lambda_1\lambda_2 z+\lambda_1^2+\lambda_2^2}{(\lambda_1+\lambda_2 z)^2(\lambda_1 z+\lambda_2)^2}\frac{2\lambda_1\lambda_2}{(1+y)^2} \\
&= \frac{\lambda_1^2 z^2+2\lambda_1\lambda_2 z+\lambda_2^2+\lambda_2^2 z^2+2\lambda_1\lambda_2 z+\lambda_1^2}{(\lambda_1+\lambda_2 z)^2(\lambda_1 z+\lambda_2)^2}\frac{2\lambda_1\lambda_2}{(1+y)^2} \\
&= \frac{(\lambda_1 z+\lambda_2)^2+(\lambda_1+\lambda_2 z)^2}{(\lambda_1+\lambda_2 z)^2(\lambda_1 z+\lambda_2)^2}\frac{2\lambda_1\lambda_2}{(1+y)^2} \\
&= \Big(\frac{1}{(\lambda_1+\lambda_2 z)^2}+\frac{1}{(\lambda_1 z+\lambda_2)^2}\Big)\frac{2\lambda_1\lambda_2}{(1+y)^2}.
\end{aligned}$$

We can simplify by replacing $z$ with its value of $\frac{1-y}{1+y}$,

$$\begin{aligned}
f_Y(y) &= \Big(\frac{1}{(\lambda_1+\lambda_2\frac{1-y}{1+y})^2}+\frac{1}{(\lambda_1\frac{1-y}{1+y}+\lambda_2)^2}\Big)\frac{2\lambda_1\lambda_2}{(1+y)^2} \\
&= \Big(\frac{1}{(\frac{\lambda_1+\lambda_1 y+\lambda_2-\lambda_2 y}{1+y})^2}+\frac{1}{(\frac{\lambda_1-\lambda_1 y+\lambda_2+\lambda_2 y}{1+y})^2}\Big)\frac{2\lambda_1\lambda_2}{(1+y)^2} \\
&= \Big(\frac{(1+y)^2}{(\lambda_1+\lambda_1 y+\lambda_2-\lambda_2 y)^2}+\frac{(1+y)^2}{(\lambda_1-\lambda_1 y+\lambda_2+\lambda_2 y)^2}\Big)\frac{2\lambda_1\lambda_2}{(1+y)^2} \\
&= \frac{2\lambda_1\lambda_2}{((\lambda_1-\lambda_2)y+\lambda_1+\lambda_2)^2}+\frac{2\lambda_1\lambda_2}{((\lambda_2-\lambda_1)y+\lambda_1+\lambda_2)^2}.
\end{aligned}$$

We can set $f_Y(0) = \frac{2\lambda_1\lambda_2}{((\lambda_1-\lambda_2)0+\lambda_1+\lambda_2)^2}+\frac{2\lambda_1\lambda_2}{((\lambda_2-\lambda_1)0+\lambda_1+\lambda_2)^2} = \frac{4\lambda_1\lambda_2}{(\lambda_1+\lambda_2)^2}$ and $f_Y(1) = \frac{2\lambda_1\lambda_2}{((\lambda_1-\lambda_2)1+\lambda_1+\lambda_2)^2}+\frac{2\lambda_1\lambda_2}{((\lambda_2-\lambda_1)1+\lambda_1+\lambda_2)^2} = \frac{2\lambda_1\lambda_2}{(2\lambda_1)^2}+\frac{2\lambda_1\lambda_2}{(2\lambda_2)^2} = \frac{1}{2}(\frac{\lambda_1}{\lambda_2}+\frac{\lambda_2}{\lambda_1})$ to make $f_Y(y)$ be defined everywhere. Then

$$f_Y(y) = \begin{cases} \frac{2\lambda_1\lambda_2}{((\lambda_1-\lambda_2)y+\lambda_1+\lambda_2)^2}+\frac{2\lambda_1\lambda_2}{((\lambda_2-\lambda_1)y+\lambda_1+\lambda_2)^2} & \text{if } 0 \le y \le 1 \\ 0 & \text{otherwise.} \end{cases}$$

Using $f_Z(\zeta) = \frac{1}{\sqrt{2}}f_Y(\frac{\zeta}{\sqrt{2}})$, we finally have for the PDF of $Z$:

$$f_Z(\zeta) = \begin{cases} \frac{\sqrt{2}\lambda_1\lambda_2}{((\lambda_1-\lambda_2)\frac{\zeta}{\sqrt{2}}+\lambda_1+\lambda_2)^2}+\frac{\sqrt{2}\lambda_1\lambda_2}{((\lambda_2-\lambda_1)\frac{\zeta}{\sqrt{2}}+\lambda_1+\lambda_2)^2} & \text{if } 0 \le \zeta \le \sqrt{2} \\ 0 & \text{otherwise.} \end{cases}$$

The PDF in the symmetric case is just

$$f_Z(\zeta) = \frac{\mathrm{d}F_Z(\zeta)}{\mathrm{d}\zeta} = \begin{cases} \frac{1}{\sqrt{2}} & \text{if } 0 \le \zeta \le \sqrt{2} \\ 0 & \text{otherwise,} \end{cases}$$

which also agrees with plugging $\lambda_1 = \lambda_2 = \lambda$ into the asymmetric result.

#### 2.2 Gamma and Weibull models

If we have reason to believe that $X_1$ and $X_2$ follow distributions that are other special cases of the generalized gamma, like the gamma and Weibull distributions, we can use the same joint and marginal methods as we did for the generalized gamma distribution. We only use a different underlying joint PDF for $(X_1, X_2)$ before we make the same variable transform.

If we believe the underlying data has a gamma distribution ($c = 1$ in the parametrization of the generalized gamma distribution), then we use the `vglm` function in the `VGAM` package in `R` to find MLEs for the remaining parameters (Yee, 2018). If $X$ has a gamma distribution with shape parameter $\alpha > 0$ and scale parameter $\lambda > 0$, its PDF is

$$f_X(x) = \begin{cases} \frac{\lambda^\alpha}{\Gamma(\alpha)} x^{\alpha-1} e^{-\lambda x} & \text{if } x \ge 0 \\ 0 & \text{otherwise} \end{cases},$$

where $\Gamma(x) = \int_0^\infty u^{x-1} e^{-u} \mathrm{d}u$ is the gamma function (Rice, 2007; Johnson and Kotz, 1970[a]). If $\alpha = 1$, then $X \sim Exp(\lambda)$. If $X_1 \sim Gamma(\alpha, \lambda_1)$ and $X_2 \sim Gamma(\alpha, \lambda_2)$ are independent, then the joint PDF for $(X_1, X_2)$ is

$$f_{X_1 X_2}(x_1, x_2) = f_{X_1}(x_1) f_{X_2}(x_2) = \begin{cases} \frac{\lambda_1^\alpha}{\Gamma(\alpha)} x_1^{\alpha-1} e^{-\lambda_1 x_1} \frac{\lambda_2^\alpha}{\Gamma(\alpha)} x_2^{\alpha-1} e^{-\lambda_2 x_2} & \text{if } x_1 \ge 0, x_2 \ge 0 \\ 0 & \text{otherwise} \end{cases}.$$

Using the same variable transform as before, we have the following joint PDF for $(\Delta, Z)$,

$$\begin{aligned} f_{\Delta,Z}(\delta,\zeta) &= f_{X_1 X_2}(\Phi\tbinom{\delta}{\zeta}) |\det([\mathbf{D}\Phi\tbinom{\delta}{\zeta}])| \\ &= \begin{cases} \begin{array}{c} \frac{\lambda_1^\alpha}{\Gamma(\alpha)}((\frac{\sqrt{2}}{\zeta}+1)\frac{\delta}{2})^{\alpha-1} e^{-\lambda_1(\frac{\sqrt{2}}{\zeta}+1)\frac{\delta}{2}} \times \\ \frac{\lambda_2^\alpha}{\Gamma(\alpha)}((\frac{\sqrt{2}}{\zeta}-1)\frac{\delta}{2})^{\alpha-1} e^{-\lambda_2(\frac{\sqrt{2}}{\zeta}-1)\frac{\delta}{2}} \frac{\sqrt{2}\delta}{2cv^2} \end{array} & \text{if } \delta \ge 0 \\ \begin{array}{c} \frac{\lambda_1^\alpha}{\Gamma(\alpha)}(-(\frac{\sqrt{2}}{\zeta}-1)\frac{\delta}{2})^{\alpha-1} e^{\lambda_1(\frac{\sqrt{2}}{\zeta}-1)\frac{\delta}{2}} \times \\ \frac{\lambda_2^\alpha}{\Gamma(\alpha)}(-(\frac{\sqrt{2}}{\zeta}+1)\frac{\delta}{2})^{\alpha-1} e^{\lambda_2(\frac{\sqrt{2}}{\zeta}+1)\frac{\delta}{2}} (-\frac{\sqrt{2}\delta}{2cv^2}) \end{array} & \text{if } \delta < 0. \end{cases} \end{aligned}$$

Using MLEs for $(\alpha_1, \lambda_1)$ and $(\alpha_2, \lambda_2)$, we can use the same joint and marginal methods as before. We define

$$q(\delta,\zeta) = \begin{cases} \mathbb{P}(\Delta \leq \delta, Z \geq \zeta) &= \int_{-\infty}^{\delta} \int_{\zeta}^{\sqrt{2}} f_{\Delta,Z}(a,b|\hat{\alpha}_1,\hat{\lambda}_1,\hat{\alpha}_2,\hat{\lambda}_2)\mathrm{d}b\mathrm{d}a & \text{if } \delta < 0 \\ \mathbb{P}(\Delta \geq \delta, Z \geq \zeta) &= \int_{d}^{\infty} \int_{\zeta}^{\sqrt{2}} f_{\Delta,Z}(a,b|\hat{\alpha}_1,\hat{\lambda}_1,\hat{\alpha}_2,\hat{\lambda}_2)\mathrm{d}b\mathrm{d}a & \text{if } \delta \geq 0 \end{cases}$$

for all of the points in the joint method, with all of the points having $q(\delta,\zeta) \leq q^*$ being outliers. For the marginal method, we still fit $\Delta$ to an asymmetric Laplace distribution, remove all of the points having a value for $\delta$ that is within $k = 1$ or $k = 2$ standard deviations away from $\hat{\mu} + \hat{\theta}$, and then calculate

$$\begin{aligned} q(\zeta) &= \mathbb{P}(Z \geq \zeta) \\ &= \int_{\zeta}^{\sqrt{2}} f_Z(b|\hat{\alpha}_1,\hat{\lambda}_1,\hat{\alpha}_2,\hat{\lambda}_2)\mathrm{d}b \\ &= \int_{\zeta}^{\sqrt{2}} \int_{-\infty}^{\infty} f_{\Delta,Z}(a,b|\hat{\alpha}_1,\hat{\lambda}_1,\hat{\alpha}_2,\hat{\lambda}_2)\mathrm{d}a\mathrm{d}b \end{aligned}$$

for the remaining points. Any remaining point with $q(\zeta) \leq q^*$ is an outlier. In both the joint and marginal methods, we rely on numerical integration like the `adaptIntegrate` function in the `R` package `cubature` (Narasimhan and Johnson, 2018).

We could just as easily assume the data points follow independent Weibull distributions, with $\alpha = 1$ in the parametrization for the generalized gamma. If $X$ has a Weibull distribution with shape parameter $c > 0$ and scale parameter $\alpha > 0$, its PDF is

$$f_X(x) = \begin{cases} \frac{c}{\alpha}(\frac{x}{\alpha})^{c-1}e^{-(\frac{x}{\alpha})^c} & \text{if } x \geq 0 \\ 0 & \text{otherwise} \end{cases}.$$

Once again, we have the special case of the exponential distribution $X \sim Exp(\frac{1}{\alpha})$ when $c = 1$ (Johnson and Kotz, 1970[a]). We can find MLEs for these parameters for each data vector using the `vglm` function in the `VGAM` package in `R` (Yee, 2018).

If $X_1 \sim Weibull(c_1, \alpha_1)$ and $X_2 \sim Weibull(c_2, \alpha_2)$ are independent, then their joint PDF is

$$f_{X_1X_2}(x_1,x_2) = f_{X_1}(x_1)f_{X_2}(x_2) = \begin{cases} \frac{c}{\alpha_1}(\frac{x_1}{\alpha_1})^{c-1}e^{-(\frac{x_1}{\alpha_1})^c} \times & \\ \frac{c}{\alpha_2}(\frac{x_2}{\alpha_2})^{c-1}e^{-(\frac{x_2}{\alpha_2})^c} & \text{if } x_1 \geq 0, x_2 \geq 0 \\ 0 & \text{otherwise} \end{cases},$$

meaning the joint PDF of $(\Delta, Z)$ is

$$f_{\Delta,Z}(\delta,\zeta) = f_{X_1X_2}(\Phi\binom{\delta}{\zeta})|\det([\mathbf{D}\Phi\binom{\delta}{\zeta}])|$$
$$= \begin{cases} \frac{c}{\alpha_1}(\frac{(\frac{\sqrt{2}}{\zeta}+1)\delta}{2\alpha_1})^{c-1}e^{-(\frac{(\frac{\sqrt{2}}{\zeta}+1)\delta}{2\alpha_1})^c}\times \\ \frac{c}{\alpha_2}(\frac{(\frac{\sqrt{2}}{\zeta}-1)\delta}{2\alpha_2})^{c-1}e^{-(\frac{(\frac{\sqrt{2}}{\zeta}-1)\delta}{2\alpha_2})^c}\frac{\sqrt{2}\delta}{2cv^2} & \text{if } \delta \geq 0 \\ \frac{c}{\alpha_1}(-\frac{(\frac{\sqrt{2}}{\zeta}-1)\delta}{2\alpha_1})^{c-1}e^{-(-\frac{(\frac{\sqrt{2}}{\zeta}-1)\delta}{2\alpha_1})^c}\times \\ \frac{c}{\alpha_2}(-\frac{(\frac{\sqrt{2}}{\zeta}+1)\delta}{2\alpha_2})^{c-1}e^{-(-\frac{(\frac{\sqrt{2}}{\zeta}+1)\delta}{2\alpha_2})^c}(-\frac{\sqrt{2}\delta}{2cv^2}) & \text{if } \delta < 0, \end{cases}$$

and

$$q(\delta,\zeta) = \begin{cases} \mathbb{P}(\Delta \leq \delta, Z \geq \zeta) = \int_{-\infty}^{\delta}\int_{\zeta}^{\sqrt{2}} f_{\Delta,Z}(a,b|\hat{c}_1,\hat{\alpha}_1,\hat{c}_2,\hat{\alpha}_2)\mathrm{d}b\mathrm{d}a & \text{if } \delta < 0 \\ \mathbb{P}(\Delta \geq \delta, Z \geq \zeta) = \int_{d}^{\infty}\int_{\zeta}^{\sqrt{2}} f_{\Delta,Z}(a,b|\hat{c}_1,\hat{\alpha}_1,\hat{c}_2,\hat{\alpha}_2)\mathrm{d}b\mathrm{d}a & \text{if } \delta \geq 0 \end{cases}$$

for the joint method, calculated using numerical integration.

For the marginal method, we again fit Weibull distributions to $X_1$ and $X_2$ and an asymmetric Laplace to $\Delta = X_1 - X_2$, considering the points with $\delta$ within $k=1$ or $k=2$ standard deviations away from $\hat{\mu}+\hat{\theta}$ to not be outliers. For $Z$, we actually have a closed-form solution for its CDF when $c_1 = c_2 = c$. We can test this by looking at the 95%-confidence intervals for $\log c_1$ and $\log c_2$ and seeing if they overlap. If they do not, then we have

$$\begin{aligned} q(\zeta) &= \mathbb{P}(Z \geq \zeta) \\ &= \int_{\zeta}^{\sqrt{2}} f_Z(b|\hat{c}_1,\hat{\alpha}_1,\hat{c}_2,\hat{\alpha}_2)\mathrm{d}b \\ &= \int_{\zeta}^{\sqrt{2}}\int_{-\infty}^{\infty} f_{\Delta.Z}(\hat{c}_1,\hat{\alpha}_1,\hat{c}_2,\hat{\alpha}_2)\mathrm{d}a\mathrm{d}b. \end{aligned}$$

If we determine $c_1 = c_2 = c$, then using $\gamma = \frac{1-\frac{\zeta}{\sqrt{2}}}{1+\frac{\zeta}{\sqrt{2}}} = z(\frac{\zeta}{\sqrt{2}})$ again, we get for the CDF of $Z$,

$$F_Z(\zeta) = \begin{cases} 0 & \text{if } \zeta \leq 0 \\ \frac{\alpha_1^c\alpha_2^c(1-\gamma^{2c})}{(\alpha_1^c+\alpha_2^c\gamma^c)(\alpha_1^c\gamma^c+\alpha_2^c)} & \text{if } 0 < \zeta < \sqrt{2} \\ 1 & \text{if } \zeta \geq \sqrt{2} \end{cases}.$$

Its PDF is then

$$f_Z(\zeta) = \begin{cases} (\frac{1}{(\alpha_1^c+\alpha_2^c\gamma^c)^2} + \frac{1}{(\alpha_1^c\gamma^c+\alpha_2^c)^2})\frac{\sqrt{2}\alpha_1^c\alpha_2^c c\gamma^{c-1}}{(1+\frac{\zeta}{\sqrt{2}})^2} & \text{if } 0 \leq \zeta \leq \sqrt{2} \\ 0 & \text{otherwise} \end{cases}.$$

If $c = 1$, then we get back to the exponential case. For outlier determination,

$$q(\zeta) = \mathbb{P}(Z \geq \zeta) = 1 - F_Z(\zeta).$$

Whether we use the closed-form $q(\zeta)$ or not, we still determine a point to be an outlier if $q(\zeta) \leq q^*$. Like in the exponential and generalized gamma cases, we can use adjusted values for $q^*$ when we have more than two replicates.

Thus far, we assumed $X_1$ and $X_2$ were continuous random variables. If we had count data, then we cannot easily use discrete distributions for the data. For example, if we assume $X_1$ and $X_2$ follow independent Poisson distributions, then $\Delta = X_1 - X_2$ has a Skellig distribution (Alzaid and Omair, 2010). In this case, $Z = \frac{\sqrt{2}|X_1 - X_2|}{X_1 + X_2}$ does not have a known distribution. Since $X_1$ and $X_2$ take values on $\mathbb{N} \cup \{0\}$, the random variable $Y = \frac{|X_1 - X_2|}{X_1 + X_2}$ takes values on $\mathbb{Q} \cap [0,1]$ (so $Z$ takes values on $\sqrt{2}(\mathbb{Q} \cap [0,1])$), and a probability mass function for $Y$ would be defined for every rational number between 0 and 1. Defining such a probability mass function (PMF) is possible because the rational numbers are countable, although there are no well-known PMFs using enumerations of the rationals (Beals, 2004; Bass, 2013; Johnson et al., 2005). We also used integration for finding the CDFs under the assumptions of different continuous distributions, and we cannot do this when $X_1$, $X_2$, $Y$, and $Z$ take discrete values. Using a discrete PMF is not necessary, since the rational numbers are dense in the reals (Skellam,, 1946; Beals, 2004). Using the continuous distributions provide adequate approximations. Count data is typically encountered in next-generation sequencing studies such as RNA-seq and its many variants that measure digital gene expression.

##### 2.2.1 Generalization to independent Weibull models

Finally, assume $X_1 \sim Weibull(c, \alpha_1)$ and $X_2 \sim Weibull(c, \alpha_2)$ are independent. If $X \sim Weibull(c, \alpha)$ has a Weibull distribution with shape parameter $c > 0$ and scale parameter $\alpha > 0$, its PDF is

$$f_X(x) = \begin{cases} \frac{c}{\alpha}(\frac{x}{\alpha})^{c-1} e^{-(\frac{x}{\alpha})^c} & \text{if } x \geq 0 \\ 0 & \text{otherwise} \end{cases}$$

(Johnson and Kotz, 1970[a]). Notice that if $c = 1$, $X \sim Exp(\frac{1}{\alpha})$, so this is a generalization of our previous work. It will be helpful for later if we had the indefinite integral of this PDF:

$$\int f_X(x) \mathrm{d}x \;\; = \;\; \int \frac{c}{\alpha}(\frac{x}{\alpha})^{c-1} e^{-(\frac{x}{\alpha})^c} \mathrm{d}x.$$

We introduce the change of variable $u = \frac{x}{\alpha}$, so $\mathrm{d}u = \frac{\mathrm{d}u}{\alpha}$. Then

$$\int f_X(x)\mathrm{d}x \quad = \quad \int cu^{c-1}e^{-u^c}\mathrm{d}u.$$

We do one more change of variable $w = u^c$, so $\mathrm{d}w = cu^{c-1}\mathrm{d}u$. Then

$$\begin{aligned}\int f_X(x)\mathrm{d}x &= \int e^{-w}\mathrm{d}w \\ &= -e^{-w} + C \\ &= -e^{-u^c} + C \\ &= -e^{-(\frac{x}{\alpha})^c} + C,\end{aligned}$$

where $C$ (uppercase) is an arbitrary real constant. We can easily differentiate this to get the previous PDF for $X$.

Consider a linearly scaled random variable $Y = \frac{1}{\sqrt{2}}Z = \frac{|X_1 - X_2|}{X_1 + X_2}$ (so $Z = \sqrt{2}Y$). If $F_Z(\zeta)$ is the CDF for $Z$ and $f_Z(\zeta)$ is the PDF for $Z$, then $F_Z(\zeta) = F_Y(\frac{\zeta}{\sqrt{2}})$ and $f_Z(\zeta) = \frac{1}{\sqrt{2}}f_Y(\frac{\zeta}{\sqrt{2}})$ given the CDF $F_Y(y)$ and PDF $f_Y(y)$ for $Y$, respectively. We then look for the CDF and PDF of $Y$. The joint PDF for $X_1$ and $X_2$ is

$$f_{X_1X_2}(x_1,x_2) = f_{X_1}(x_1)f_{X_2}(x_2) = \begin{cases} \frac{c}{\alpha_1}(\frac{x_1}{\alpha_1})^{c-1}e^{-(\frac{x_1}{\alpha_1})^c} \times & \\ \frac{c}{\alpha_2}(\frac{x_2}{\alpha_2})^{c-1}e^{-(\frac{x_2}{\alpha_2})^c} & \text{if } x_1 \geq 0, x_2 \geq 0 \\ 0 & \text{otherwise} \end{cases}.$$

If we let $X_1$ and $X_2$ only take non-zero values, $Y$ cannot take on a negative value or a value greater than 1. Then $F_Y(y) = 0$ for $y \leq 0$ and $F_Y(y) = 1$ for $y \geq 1$. So we look at $0 < y < 1$. By the definition of a CDF:

$$\begin{aligned}F_Y(y) &= \mathbb{P}(Y \leq y) \\ &= \mathbb{P}(\tfrac{|X_1-X_2|}{X_1+X_2} \leq y) \\ &= \mathbb{P}(|X_1 - X_2| \leq yX_1 + yX_2) \\ &= \mathbb{P}(|X_1 - X_2| \leq yX_1 + yX_2, X_1 - X_2 \geq 0) \\ &\quad + \mathbb{P}(|X_1 - X_2| \leq yX_1 + yX_2, X_1 - X_2 < 0) \\ &= \mathbb{P}(X_1 - X_2 \leq yX_1 + yX_2, X_1 - X_2 \geq 0) \\ &\quad + \mathbb{P}(-(X_1 - X_2) \leq yX_1 + yX_2, X_1 - X_2 < 0) \\ &= \mathbb{P}(X_1 - yX_1 \leq X_2 + yX_2, X_1 \geq X_2) \\ &\quad + \mathbb{P}(X_2 - X_1 \leq yX_1 + yX_2, X_1 < X_2) \\ &= \mathbb{P}(\tfrac{1}{1+y}(X_1 - yX_1) \leq X_2, X_1 \geq X_2) \\ &\quad + \mathbb{P}(X_2 - yX_2 \leq X_1 + yX_1, X_1 < X_2) \\ &= \mathbb{P}(X_2 \geq \tfrac{1}{1+y}(X_1 - yX_1), X_2 \leq X_1) \\ &\quad + \mathbb{P}(X_2 \leq \tfrac{1}{1-y}(X_1 + yX_1), X_2 > X_1) \\ &= \mathbb{P}(X_2 \geq \tfrac{1-y}{1+y}X_1, X_2 \leq X_1) + \mathbb{P}(X_2 \leq \tfrac{1+y}{1-y}X_1, X_2 > X_1) \\ &= \mathbb{P}(\Delta) + \mathbb{P}(E),\end{aligned}$$

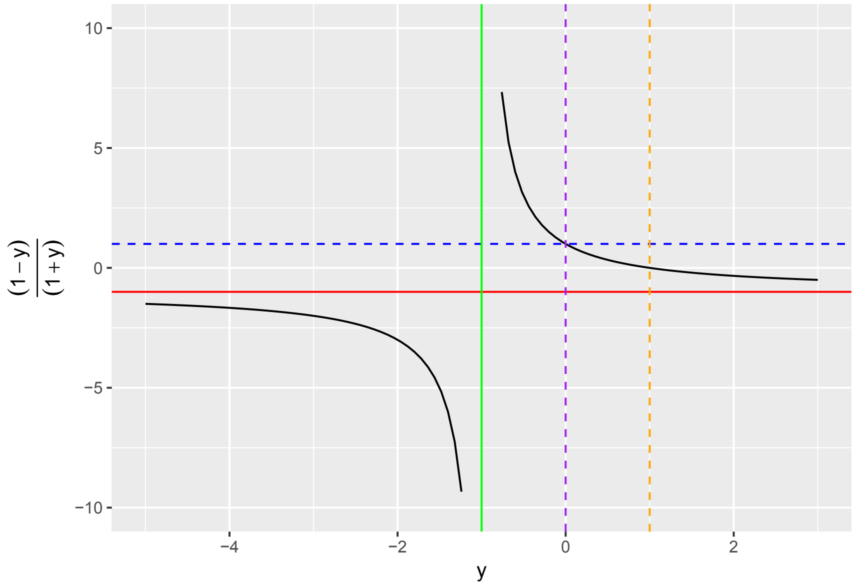


where $\Delta = \{X_2 \geq \frac{1-y}{1+y}X_1, X_2 \leq X_1\}$ and $E = \{X_2 \leq \frac{1+y}{1-y}X_1, X_2 > X_1\}$. Notice we can divide by $1-y$ without needing to change the direction of the inequality because $0 < y < 1$. Like we did in the exponential case, we need to understand the signs and magnitudes of $\frac{1-y}{1+y}$ and $\frac{1+y}{1-y}$ to know what regions we are integrating over. For $\frac{1-y}{1+y}$, we have the figure shown above (which we will use to determine $\Delta$): There is a vertical asymptote at $y = -1$ and a horizontal asymptote at $\frac{1-y}{1+y} = -1$. Since we are only interested in $y > 0$, we see that $-1 < \frac{1-y}{1+y} < 1$. For $0 < y < 1$, we have $0 < \frac{1-y}{1+y} < 1$.

For $\frac{1+y}{1-y}$, we have the figure shown on the next page (which we will use to determine $E$): Since we are only interested in $0 < y < 1$, we see that $1 < \frac{1+y}{1-y} < \infty$ in $E$.

Knowing that the slopes of the lines defining $\Delta$ and $E$ lie in these ranges, we can draw the proper graphs of these regions.

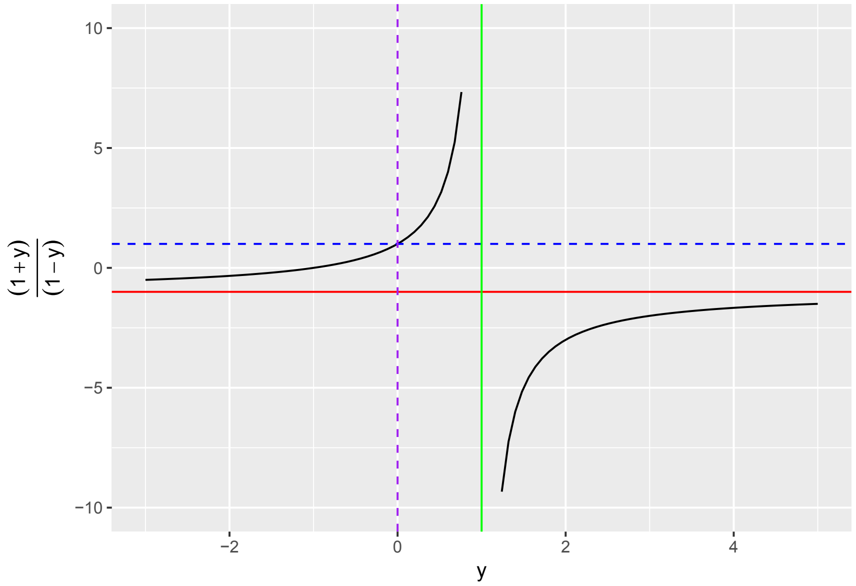


For $\Delta$, we have

$$
\begin{aligned}
\mathbb{P}(\Delta) &= \iint\limits_D f_{X_1X_2}(x_1,x_2)\mathrm{d}^2\vec{x}\\
&= \int_0^\infty \int_{\frac{1-y}{1+y}x_1}^{x_1} \frac{c}{\alpha_1}\big(\frac{x_1}{\alpha_1}\big)^{c-1}e^{-(\frac{x_1}{\alpha_1})^c}\frac{c}{\alpha_2}\big(\frac{x_2}{\alpha_2}\big)^{c-1}e^{-(\frac{x_2}{\alpha_2})^c}\mathrm{d}x_2\mathrm{d}x_1\\
&= \int_0^\infty \frac{c}{\alpha_1}\big(\frac{x_1}{\alpha_1}\big)^{c-1}e^{-(\frac{x_1}{\alpha_1})^c}\Big(\int_{\frac{1-y}{1+y}x_1}^{x_1}\frac{c}{\alpha_2}\big(\frac{x_2}{\alpha_2}\big)^{c-1}e^{-(\frac{x_2}{\alpha_2})^c}\mathrm{d}x_2\Big)\mathrm{d}x_1\\
&= \int_0^\infty \frac{c}{\alpha_1}\big(\frac{x_1}{\alpha_1}\big)^{c-1}e^{-(\frac{x_1}{\alpha_1})^c}\big(-e^{-(\frac{x_2}{\alpha_2})^c}\big)\Big]_{\frac{1-y}{1+y}x_1}^{x_1}\mathrm{d}x_1\\
&= \int_0^\infty \frac{c}{\alpha_1}\big(\frac{x_1}{\alpha_1}\big)^{c-1}e^{-(\frac{x_1}{\alpha_1})^c}\big(e^{-(\frac{(1-y)x_1}{(1+y)\alpha_2})^c}-e^{-(\frac{x_1}{\alpha_2})^c}\big)\mathrm{d}x_1\\
&= \int_0^\infty \frac{c}{\alpha_1}\big(\frac{x_1}{\alpha_1}\big)^{c-1}\big(e^{-(\frac{x_1}{\alpha_1})^c-(\frac{(1-y)x_1}{(1+y)\alpha_2})^c}-e^{-(\frac{x_1}{\alpha_1})^c-(\frac{x_1}{\alpha_2})^c}\big)\mathrm{d}x_1\\
&= \int_0^\infty \frac{c}{\alpha_1}\big(\frac{x_1}{\alpha_1}\big)^{c-1}\big(e^{-x_1^c(\frac{1}{\alpha_1^c}+\frac{(1-y)^c}{(1+y)^c\alpha_2^c})}-e^{-x_1^c(\frac{1}{\alpha_1^c}+\frac{1}{\alpha_2^c})}\big)\mathrm{d}x_1\\
&= \int_0^\infty \frac{c}{\alpha_1}\big(\frac{x_1}{\alpha_1}\big)^{c-1}e^{-x_1^c(\frac{1}{\alpha_1^c}+\frac{(1-y)^c}{(1+y)^c\alpha_2^c})}\mathrm{d}x_1-\int_0^\infty \frac{c}{\alpha_1}\big(\frac{x_1}{\alpha_1}\big)^{c-1}e^{-x_1^c(\frac{1}{\alpha_1^c}+\frac{1}{\alpha_2^c})}\mathrm{d}x_1.
\end{aligned}
$$

Surprisingly, integrals of this type are solvable. If we have positive numbers $a$, $b$, and $c$, then

$$\begin{aligned}
\int_0^\infty \tfrac{c}{a}(\tfrac{x}{a})^{c-1}e^{-(\frac{x}{b})^c}\mathrm{d}x &= \tfrac{1}{a^c}\int_0^\infty cx^{c-1}e^{-(\frac{x}{b})^c}\mathrm{d}x \\
&= \tfrac{b^c}{a^c}\int_0^\infty \tfrac{1}{b^c}cx^{c-1}e^{-(\frac{x}{b})^c}\mathrm{d}x \\
&= \tfrac{b^c}{a^c}\int_0^\infty \tfrac{c}{b}(\tfrac{x}{b})^{c-1}e^{-(\frac{x}{b})^c}\mathrm{d}x \\
&= \tfrac{b^c}{a^c}(-e^{-(\frac{x}{b})^c})\Big]_0^\infty \\
&= \tfrac{b^c}{a^c}(-0-(-e^0)) \\
&= \tfrac{b^c}{a^c}(0+1) \\
&= \tfrac{b^c}{a^c}.
\end{aligned}$$

We can show that $a$ and $b$ in the preceeding integrals are positive for the allowed values of $y$.

For $\int_0^\infty \frac{c}{\alpha_1}(\frac{x_1}{\alpha_1})^{c-1}e^{-x_1^c(\frac{1}{\alpha_1^c}+\frac{(1-y)^c}{(1+y)^c\alpha_2^c})}\mathrm{d}x_1$, $\frac{1}{b^c}=\frac{1}{\alpha_1^c}+\frac{(1-y)^c}{(1+y)^c\alpha_2^c}$ and $a=\alpha_1$, so

$$\frac{b^c}{a^c}=\frac{\frac{1}{\frac{1}{\alpha_1^c}+\frac{(1-y)^c}{(1+y)^c\alpha_2^c}}}{\alpha_1^c}=\frac{1}{1+\frac{(1-y)^c\alpha_1^c}{(1+y)^c\alpha_2^c}}=\frac{(1+y)^c\alpha_2^c}{(1-y)^c\alpha_1^c+(1+y)^c\alpha_2^c}.$$

For $\int_0^\infty \frac{c}{\alpha_1}(\frac{x_1}{\alpha_1})^{c-1}e^{-x_1^c(\frac{1}{\alpha_1^c}+\frac{1}{\alpha_2^c})}\mathrm{d}x_1$, $\frac{1}{b^c}=\frac{1}{\alpha_1^c}+\frac{1}{\alpha_2^c}$ and $a=\alpha_1$, so

$$\frac{b^c}{a^c}=\frac{\frac{1}{\frac{1}{\alpha_1^c}+\frac{1}{\alpha_2^c}}}{\alpha_1^c}=\frac{1}{1+\frac{\alpha_1^c}{\alpha_2^c}}=\frac{\alpha_2^c}{\alpha_1^c+\alpha_2^c}.$$

Therefore,

$$\begin{aligned}
\mathbb{P}(\Delta) &= \int_0^\infty \tfrac{c}{\alpha_1}(\tfrac{x_1}{\alpha_1})^{c-1}e^{-x_1^c(\frac{1}{\alpha_1^c}+\frac{(1-y)^c}{(1+y)^c\alpha_2^c})}\mathrm{d}x_1-\int_0^\infty \tfrac{c}{\alpha_1}(\tfrac{x_1}{\alpha_1})^{c-1}e^{-x_1^c(\frac{1}{\alpha_1^c}+\frac{1}{\alpha_2^c})}\mathrm{d}x_1 \\
&= \tfrac{(1+y)^c\alpha_2^c}{(1-y)^c\alpha_1^c+(1+y)^c\alpha_2^c}-\tfrac{\alpha_2^c}{\alpha_1^c+\alpha_2^c} \\
&= \alpha_2^c(\tfrac{(1+y)^c}{(1-y)^c\alpha_1^c+(1+y)^c\alpha_2^c}-\tfrac{1}{\alpha_1^c+\alpha_2^c}) \\
&= \alpha_2^c\tfrac{(1+y)^c\alpha_1^c+(1+y)^c\alpha_2^c-(1-y)^c\alpha_1^c-(1+y)^c\alpha_2^c}{((1-y)^c\alpha_1^c+(1+y)^c\alpha_2^c)(\alpha_1^c+\alpha_2^c)} \\
&= \alpha_1^c\alpha_2^c\tfrac{(1+y)^c-(1-y)^c}{((1-y)^c\alpha_1^c+(1+y)^c\alpha_2^c)(\alpha_1^c+\alpha_2^c)}.
\end{aligned}$$

Now we look at $E$:

$$\begin{aligned}
\mathbb{P}(E) &= \iint\limits_E f_{X_1X_2}(x_1,x_2)\mathrm{d}^2\vec{x} \\
&= \int_0^\infty \int_{x_1}^{\frac{1+y}{1-y}x_1} \tfrac{c}{\alpha_1}(\tfrac{x_1}{\alpha_1})^{c-1}e^{-(\frac{x_1}{\alpha_1})^c} \tfrac{c}{\alpha_2}(\tfrac{x_2}{\alpha_2})^{c-1}e^{-(\frac{x_2}{\alpha_2})^c}\mathrm{d}x_2\mathrm{d}x_1 \\
&= \int_0^\infty \tfrac{c}{\alpha_1}(\tfrac{x_1}{\alpha_1})^{c-1}e^{-(\frac{x_1}{\alpha_1})^c}\Big(\int_{x_1}^{\frac{1+y}{1-y}x_1} \tfrac{c}{\alpha_2}(\tfrac{x_2}{\alpha_2})^{c-1}e^{-(\frac{x_2}{\alpha_2})^c}\mathrm{d}x_2\Big)\mathrm{d}x_1 \\
&= \int_0^\infty \tfrac{c}{\alpha_1}(\tfrac{x_1}{\alpha_1})^{c-1}e^{-(\frac{x_1}{\alpha_1})^c}(-e^{-(\frac{x_2}{\alpha_2})^c})\Big]_{x_1}^{\frac{1+y}{1-y}x_1}\mathrm{d}x_1 \\
&= \int_0^\infty \tfrac{c}{\alpha_1}(\tfrac{x_1}{\alpha_1})^{c-1}e^{-(\frac{x_1}{\alpha_1})^c}(e^{-(\frac{x_1}{\alpha_2})^c} - e^{-(\frac{(1+y)x_1}{(1-y)\alpha_2})^c})\mathrm{d}x_1 \\
&= \int_0^\infty \tfrac{c}{\alpha_1}(\tfrac{x_1}{\alpha_1})^{c-1}(e^{-(\frac{x_1}{\alpha_1})^c-(\frac{x_1}{\alpha_2})^c} - e^{-(\frac{x_1}{\alpha_1})^c-(\frac{(1+y)x_1}{(1-y)\alpha_2})^c})\mathrm{d}x_1 \\
&= \int_0^\infty \tfrac{c}{\alpha_1}(\tfrac{x_1}{\alpha_1})^{c-1}(e^{-x_1^c(\frac{1}{\alpha_1^c}+\frac{1}{\alpha_2^c})} - e^{-x_1^c(\frac{1}{\alpha_1^c}+\frac{(1+y)^c}{(1-y)^c\alpha_2^c})})\mathrm{d}x_1 \\
&= \int_0^\infty \tfrac{c}{\alpha_1}(\tfrac{x_1}{\alpha_1})^{c-1}e^{-x_1^c(\frac{1}{\alpha_1^c}+\frac{1}{\alpha_2^c})}\mathrm{d}x_1 - \int_0^\infty \tfrac{c}{\alpha_1}(\tfrac{x_1}{\alpha_1})^{c-1}e^{-x_1^c(\frac{1}{\alpha_1^c}+\frac{(1+y)^c}{(1-y)^c\alpha_2^c})}\mathrm{d}x_1.
\end{aligned}$$

These are the same types of integrals we computed previously, and we can show that the values for $a$, $b$, and $c$ are all positive. For $\int_0^\infty \frac{c}{\alpha_1}(\frac{x_1}{\alpha_1})^{c-1}e^{-x_1^c(\frac{1}{\alpha_1^c}+\frac{1}{\alpha_2^c})}\mathrm{d}x_1$, $\frac{1}{b^c} = \frac{1}{\alpha_1^c}+\frac{1}{\alpha_2^c}$ and $a=\alpha_1$, so

$$\frac{b^c}{a^c} = \frac{\frac{1}{\frac{1}{\alpha_1^c}+\frac{1}{\alpha_2^c}}}{\alpha_1^c} = \frac{1}{1+\frac{\alpha_1^c}{\alpha_2^c}} = \frac{\alpha_2^c}{\alpha_1^c+\alpha_2^c},$$

like before in calculating $\Delta$.

For $\int_0^\infty \frac{c}{\alpha_1}(\frac{x_1}{\alpha_1})^{c-1}e^{-x_1^c(\frac{1}{\alpha_1^c}+\frac{(1+y)^c}{(1-y)^c\alpha_2^c})}\mathrm{d}x_1$, $\frac{1}{b^c} = \frac{1}{\alpha_1^c}+\frac{(1+y)^c}{(1-y)^c\alpha_2^c}$ and $a=\alpha_1$, so

$$\frac{b^c}{a^c} = \frac{\frac{1}{\frac{1}{\alpha_1^c}+\frac{(1+y)^c}{(1-y)^c\alpha_2^c}}}{\alpha_1^c} = \frac{1}{1+\frac{(1+y)^c\alpha_1^c}{(1-y)^c\alpha_2^c}} = \frac{(1-y)^c\alpha_2^c}{(1+y)^c\alpha_1^c+(1-y)^c\alpha_2^c}.$$

Therefore,

$$\begin{aligned}
\mathbb{P}(E) &= \int_0^\infty \tfrac{c}{\alpha_1}(\tfrac{x_1}{\alpha_1})^{c-1}e^{-x_1^c(\frac{1}{\alpha_1^c}+\frac{1}{\alpha_2^c})}\mathrm{d}x_1 - \int_0^\infty \tfrac{c}{\alpha_1}(\tfrac{x_1}{\alpha_1})^{c-1}e^{-x_1^c(\frac{1}{\alpha_1^c}+\frac{(1+y)^c}{(1-y)^c\alpha_2^c})}\mathrm{d}x_1 \\
&= \tfrac{\alpha_2^c}{\alpha_1^c+\alpha_2^c} - \tfrac{(1-y)^c\alpha_2^c}{(1+y)^c\alpha_1^c+(1-y)^c\alpha_2^c} \\
&= \alpha_2^c(\tfrac{1}{\alpha_1^c+\alpha_2^c} - \tfrac{(1-y)^c}{(1+y)^c\alpha_1^c+(1-y)^c\alpha_2^c}) \\
&= \alpha_2^c\tfrac{(1+y)^c\alpha_1^c+(1-y)^c\alpha_2^c-(1-y)^c\alpha_1^c-(1-y)^c\alpha_2^c}{((1+y)^c\alpha_1^c+(1-y)^c\alpha_2^c)(\alpha_1^c+\alpha_2^c)} \\
&= \alpha_1^c\alpha_2^c\tfrac{(1+y)^c-(1-y)^c}{((1+y)^c\alpha_1^c+(1-y)^c\alpha_2^c)(\alpha_1^c+\alpha_2^c)}.
\end{aligned}$$

Then

$$\begin{aligned}
F_Y(y) &= \mathbb{P}(\Delta)+\mathbb{P}(E)\\
&= \alpha_1^c\alpha_2^c\frac{(1+y)^c-(1-y)^c}{((1-y)^c\alpha_1^c+(1+y)^c\alpha_2^c)(\alpha_1^c+\alpha_2^c)}+\alpha_1^c\alpha_2^c\frac{(1+y)^c-(1-y)^c}{((1+y)^c\alpha_1^c+(1-y)^c\alpha_2^c)(\alpha_1^c+\alpha_2^c)}\\
&= \frac{\alpha_1^c\alpha_2^c((1+y)^c-(1-y)^c)}{\alpha_1^c+\alpha_2^c}\left(\frac{1}{(1-y)^c\alpha_1^c+(1+y)^c\alpha_2^c}+\frac{1}{(1+y)^c\alpha_1^c+(1-y)^c\alpha_2^c}\right)\\
&= \frac{\alpha_1^c\alpha_2^c((1+y)^c-(1-y)^c)}{\alpha_1^c+\alpha_2^c}\left(\frac{(1+y)^c\alpha_1^c+(1-y)^c\alpha_2^c+(1-y)^c\alpha_1^c+(1+y)^c\alpha_2^c}{((1-y)^c\alpha_1^c+(1+y)^c\alpha_2^c)((1+y)^c\alpha_1^c+(1-y)^c\alpha_2^c)}\right)\\
&= \frac{\alpha_1^c\alpha_2^c((1+y)^c-(1-y)^c)}{\alpha_1^c+\alpha_2^c}\left(\frac{((1+y)^c+(1-y)^c)\alpha_1^c+((1+y)^c+(1-y)^c)\alpha_2^c}{((1-y)^c\alpha_1^c+(1+y)^c\alpha_2^c)((1+y)^c\alpha_1^c+(1-y)^c\alpha_2^c)}\right)\\
&= \frac{\alpha_1^c\alpha_2^c((1+y)^c-(1-y)^c)}{\alpha_1^c+\alpha_2^c}\left(\frac{((1+y)^c+(1-y)^c)(\alpha_1^c+\alpha_2^c)}{((1-y)^c\alpha_1^c+(1+y)^c\alpha_2^c)((1+y)^c\alpha_1^c+(1-y)^c\alpha_2^c)}\right)\\
&= \frac{\alpha_1^c\alpha_2^c((1+y)^c-(1-y)^c)((1+y)^c+(1-y)^c)}{((1-y)^c\alpha_1^c+(1+y)^c\alpha_2^c)((1+y)^c\alpha_1^c+(1-y)^c\alpha_2^c)}\\
&= \frac{\alpha_1^c\alpha_2^c((1+y)^{2c}-(1-y)^{2c})}{((1-y)^c\alpha_1^c+(1+y)^c\alpha_2^c)((1+y)^c\alpha_1^c+(1-y)^c\alpha_2^c)}.
\end{aligned}$$

Let $z=\frac{1-y}{1+y}$. Then

$$\begin{aligned}
F_Y(y) &= \frac{\alpha_1^c\alpha_2^c((1+y)^{2c}-(1-y)^{2c})}{((1-y)^c\alpha_1^c+(1+y)^c\alpha_2^c)((1+y)^c\alpha_1^c+(1-y)^c\alpha_2^c)}*\frac{\frac{1}{(1+y)^{2c}}}{\frac{1}{(1+y)^{2c}}}\\
&= \frac{\alpha_1^c\alpha_2^c(1-(\frac{1-y}{1+y})^{2c})}{((\frac{1-y}{1+y})^c\alpha_1^c+\alpha_2^c)(\alpha_1^c+(\frac{1-y}{1+y})^c\alpha_2^c)}\\
&= \frac{\alpha_1^c\alpha_2^c(1-z^{2c})}{(z^c\alpha_1^c+\alpha_2^c)(\alpha_1^c+z^c\alpha_2^c)}.
\end{aligned}$$

Therefore,

$$F_Y(y)=\begin{cases} 0 & \text{if } y\le 0\\ \frac{\alpha_1^c\alpha_2^c(1-z^{2c})}{(\alpha_1^c+\alpha_2^c z^c)(\alpha_1^c z^c+\alpha_2^c)} & \text{if } 0<y<1\\ 1 & \text{if } y\ge 1\end{cases}.$$

We can see that this CDF is continuous at $y=0$ $(z=1)$ and $y=1$ $(z=0)$.

Recall that an exponentially distributed random variable with parameter $\lambda$ is a special case of a Weibull-distributed random variable, having parameters $c=1$ and $\alpha=\frac{1}{\lambda}$. So if we plug in $c=1$, $\alpha_1=\frac{1}{\lambda_1}$, and $\alpha_2=\frac{1}{\lambda_2}$ into the above expression, for $0<y<1$ we get

$$\begin{aligned}
F_Y(y) &= \frac{\alpha_1\alpha_2(1-z^2)}{(\alpha_1+\alpha_2 z)(\alpha_1 z+\alpha_2)}\\
&= \frac{1-z^2}{\lambda_1\lambda_2(\frac{1}{\lambda_1}+\frac{z}{\lambda_2})(\frac{z}{\lambda_1}+\frac{1}{\lambda_2})}\\
&= \frac{1-z^2}{(\lambda_2+\lambda_1 z)(\frac{z}{\lambda_1}+\frac{1}{\lambda_2})}*\frac{\lambda_1\lambda_2}{\lambda_1\lambda_2}\\
&= \frac{\lambda_1\lambda_2(1-z^2)}{(\lambda_2+\lambda_1 z)(\lambda_2 z+\lambda_1)},
\end{aligned}$$

which is exactly what we calculated for the CDF of Y in the exponential case, along with $F_Y(y)=0$ for $y\le 0$ and $F_Y(y)=1$ for $y\ge 1$.

Using $F_Z(\zeta) = F_Y(\frac{\zeta}{\sqrt{2}})$ and $\gamma = \frac{1-\frac{\zeta}{\sqrt{2}}}{1+\frac{\zeta}{\sqrt{2}}} = z(\frac{\zeta}{\sqrt{2}})$, we finally get

$$F_Z(\zeta) = \begin{cases} 0 & \text{if } \zeta \le 0 \\ \frac{\alpha_1^c \alpha_2^c (1-\gamma^{2c})}{(\alpha_1^c + \alpha_2^c \gamma^c)(\alpha_1^c \gamma^c + \alpha_2^c)} & \text{if } 0 < \zeta < \sqrt{2} \\ 1 & \text{if } \zeta \ge \sqrt{2} \end{cases} .$$

##### 2.2.2 Marginal PDF for $Z$

We finally find the marginal PDF $f_Z(\zeta)$ for $Z$, starting by finding the marginal PDF $f_Y(y)$ of $Y$. We know that $f_Y(y) = \frac{\mathrm{d}F_Y(y)}{\mathrm{d}y}$, but the differentiation will be easier if we first simplify $F_Y(y)$:

$$\begin{aligned} F_Y(y) &= \frac{\alpha_1^c \alpha_2^c (1-z^{2c})}{(\alpha_1^c + \alpha_2^c z^c)(\alpha_1^c z^c + \alpha_2^c)} \\ &= \frac{\alpha_1^c \alpha_2^c (1-z^{2c})}{\alpha_1^{2c} z^c + \alpha_1^c \alpha_2^c + \alpha_1^c \alpha_2^c z^{2c} + \alpha_2^{2c} z^c} \\ &= \frac{\alpha_1^c \alpha_2^c (1-z^{2c})}{(\alpha_1^{2c} + \alpha_2^{2c}) z^c + (1+z^{2c}) \alpha_1^c \alpha_2^c} \\ &= \frac{\alpha_1^c \alpha_2^c (1-z^{2c})}{(\frac{\alpha_1^c}{\alpha_2^c} + \frac{\alpha_2^c}{\alpha_1^c}) \alpha_1^c \alpha_2^c z^c + (1+z^{2c}) \alpha_1^c \alpha_2^c} \\ &= \frac{1-z^{2c}}{z^{2c} + \alpha_0 z^c + 1} \end{aligned}$$

using $\alpha_0 = \frac{\alpha_1^c}{\alpha_2^c} + \frac{\alpha_2^c}{\alpha_1^c}$. Notice also that $\alpha_1^c \alpha_2^c (z^{2c} + \alpha_0 z^c + 1) = (\alpha_1^c + \alpha_2^c z^c)(\alpha_1^c z^c + \alpha_2^c)$. These calculations should look similar to ones we did for the exponential case. Using the chain rule, we have

$$
\begin{aligned}
f_Y(y) &= \frac{\mathrm{d}F_Y(y)}{\mathrm{d}y} \\
&= \frac{\mathrm{d}F_Y(z)}{\mathrm{d}z}\frac{\mathrm{d}z}{\mathrm{d}y} \\
&= \frac{\mathrm{d}}{\mathrm{d}z}\left(\frac{1-z^{2c}}{z^{2c}+\alpha_0 z^c+1}\right)\frac{\mathrm{d}}{\mathrm{d}y}\left(\frac{1-y}{1+y}\right) \\
&= \frac{(z^{2c}+\alpha_0 z^c+1)(-2cz^{2c-1})-(1-z^{2c})(2cz^{2c-1}+\alpha_0 cz^{c-1})}{(z^{2c}+\alpha_0 z^c+1)^2}\frac{(1+y)(-1)-(1-y)(1)}{(1+y)^2} \\
&= \frac{-2cz^{4c-1}-2\alpha_0 cz^{3c-1}-2cz^{2c-1}-2cz^{2c-1}-\alpha_0 cz^{c-1}+2cz^{4c-1}+\alpha_0 cz^{3c-1}}{\left(\frac{(\alpha_1^c+\alpha_2^c z^c)(\alpha_1^c z^c+\alpha_2^c)}{\alpha_1^c\alpha_2^c}\right)^2}\frac{-1-y-1+y}{(1+y)^2} \\
&= \frac{(-\alpha_0 cz^{3c-1}-4cz^{2c-1}-\alpha_0 cz^{c-1})\alpha_1^{2c}\alpha_2^{2c}}{(\alpha_1^c+\alpha_2^c z^c)^2(\alpha_1^c z^c+\alpha_2^c)^2}\frac{-2}{(1+y)^2} \\
&= \frac{\alpha_0\alpha_1^c\alpha_2^c cz^{3c-1}+4\alpha_1^c\alpha_2^c cz^{2c-1}+\alpha_0\alpha_1^c\alpha_2^c cz^{c-1}}{(\alpha_1^c+\alpha_2^c z^c)^2(\alpha_1^c z^c+\alpha_2^c)^2}\frac{2\alpha_1^c\alpha_2^c}{(1+y)^2} \\
&= \frac{(\frac{\alpha_1^c}{\alpha_2^c}+\frac{\alpha_2^c}{\alpha_1^c})\alpha_1^c\alpha_2^c cz^{3c-1}+4\alpha_1^c\alpha_2^c cz^{2c-1}+(\frac{\alpha_1^c}{\alpha_2^c}+\frac{\alpha_2^c}{\alpha_1^c})\alpha_1^c\alpha_2^c cz^{c-1}}{(\alpha_1^c+\alpha_2^c z^c)^2(\alpha_1^c z^c+\alpha_2^c)^2}\frac{2\alpha_1^c\alpha_2^c}{(1+y)^2} \\
&= \frac{(\alpha_1^{2c}+\alpha_2^{2c})cz^{3c-1}+4\alpha_1^c\alpha_2^c cz^{2c-1}+(\alpha_1^{2c}+\alpha_2^{2c})cz^{c-1}}{(\alpha_1^c+\alpha_2^c z^c)^2(\alpha_1^c z^c+\alpha_2^c)^2}\frac{2\alpha_1^c\alpha_2^c}{(1+y)^2} \\
&= \frac{(\alpha_1^{2c}+\alpha_2^{2c})z^{2c}+4\alpha_1^c\alpha_2^c z^c+\alpha_1^{2c}+\alpha_2^{2c}}{(\alpha_1^c+\alpha_2^c z^c)^2(\alpha_1^c z^c+\alpha_2^c)^2}\frac{2\alpha_1^c\alpha_2^c cz^{c-1}}{(1+y)^2} \\
&= \frac{\alpha_1^{2c}z^{2c}+2\alpha_1^c\alpha_2^c z^c+\alpha_2^{2c}+\alpha_2^{2c}z^{2c}+2\alpha_1^c\alpha_2^c z^c+\alpha_1^{2c}}{(\alpha_1^c+\alpha_2^c z^c)^2(\alpha_1^c z^c+\alpha_2^c)^2}\frac{2\alpha_1^c\alpha_2^c cz^{c-1}}{(1+y)^2} \\
&= \frac{(\alpha_1^c z^c+\alpha_2^c)^2+(\alpha_1^c+\alpha_2^c z^c)^2}{(\alpha_1^c+\alpha_2^c z^c)^2(\alpha_1^c z^c+\alpha_2^c)^2}\frac{2\alpha_1^c\alpha_2^c cz^{c-1}}{(1+y)^2} \\
&= \left(\frac{1}{(\alpha_1^c+\alpha_2^c z^c)^2}+\frac{1}{(\alpha_1^c z^c+\alpha_2^c)^2}\right)\frac{2\alpha_1^c\alpha_2^c cz^{c-1}}{(1+y)^2}.
\end{aligned}
$$

We cannot simplify this further, plugging in $z=\frac{1-y}{1+y}$, until we know the power $c$. Notice that this agrees with one of our simplified expressions for $f_Y(y)$ in the exponential case, if we let $c=1$, $\alpha_1=\frac{1}{\lambda_1}$, and $\alpha_2=\frac{1}{\lambda_2}$:

$$f_Y(y)=\left(\frac{1}{(\lambda_1+\lambda_2 z)^2}+\frac{1}{(\lambda_1 z+\lambda_2)^2}\right)\frac{2\lambda_1\lambda_2}{(1+y)^2}.$$

Back in the Weibull case, $f_Y(y)=\frac{\mathrm{d}F_Y(y)}{\mathrm{d}y}=0$ for $y<0$ and $y>1$, since $F_Y(y)$ is constant in these regions. We can define $f_Y(y)=0$ for $y=0$ $(z=1)$ and $y=1$ $(z=0)$ to make the PDF be defined everywhere. So, we have

$$f_Y(y)=\begin{cases}\left(\frac{1}{(\alpha_1^c+\alpha_2^c z^c)^2}+\frac{1}{(\alpha_1^c z^c+\alpha_2^c)^2}\right)\frac{2\alpha_1^c\alpha_2^c cz^{c-1}}{(1+y)^2} & \text{if } 0\le y\le 1 \\ 0 & \text{otherwise}\end{cases}.$$

Using $f_Z(\zeta)=\frac{1}{\sqrt{2}}f_Y(\frac{\zeta}{\sqrt{2}})$ and $\gamma=\frac{1-\frac{\zeta}{\sqrt{2}}}{1+\frac{\zeta}{\sqrt{2}}}=z(\frac{\zeta}{\sqrt{2}})$, we finally have the PDF for $Z$:

$$f_Z(\zeta)=\begin{cases}\left(\frac{1}{(\alpha_1^c+\alpha_2^c\gamma^c)^2}+\frac{1}{(\alpha_1^c\gamma^c+\alpha_2^c)^2}\right)\frac{\sqrt{2}\alpha_1^c\alpha_2^c c\gamma^{c-1}}{(1+\frac{\zeta}{\sqrt{2}})^2} & \text{if } 0\le y\le 1 \\ 0 & \text{otherwise}\end{cases}.$$

#### 2.3 Generalized Gamma Family of Models

The generalized gamma model includes many familiar models with support on $\mathbb{R}_+ \cup \{0\}$ as special cases. These include the gamma, exponential, chi-square, Weibull, half-normal, chi, Rayleigh and Maxwell-Boltzmann, thus making it an extremely flexible family that allows modeling of a variety of shapes. Using the parametrization introduced in §4 of the main paper, we have

$$c = 1 \implies X \sim Gamma(\alpha, \tfrac{1}{\beta})$$

$$\begin{aligned} \alpha &= \tfrac{n}{2}, \\ \beta &= 2, \\ c &= 1, \\ n &\in \mathbb{N} \implies X \sim \chi^2_n \end{aligned}$$

$$\begin{aligned} \alpha &= 1, \\ c &= 1 \implies X \sim Exp(\tfrac{1}{\beta}) \end{aligned}$$

$$\alpha = 1 \implies X \sim Weibull(c, \beta)$$

$$\begin{aligned} \alpha &= \tfrac{1}{2}, \\ \beta &= \sqrt{2}\sigma, \\ c &= 2, \\ \sigma &> 0 \implies X \sim Half\text{-}Normal(\sigma) \end{aligned}$$

$$\begin{aligned} \alpha &= \tfrac{n}{2}, \\ \beta &= \sqrt{2}, \\ c &= 2, \\ n &\in \mathbb{N} \implies X \sim \chi_n \end{aligned}$$

$$\begin{aligned} \alpha &= 1, \\ \beta &= \sqrt{2}\sigma, \\ c &= 2, \\ \sigma &> 0 \implies X \sim Rayleigh(\sigma) \end{aligned}$$

$$\begin{aligned} \alpha &= \tfrac{3}{2}, \\ \beta &= \sqrt{2}\sigma, \\ c &= 2, \\ \sigma &> 0 \implies X \sim Maxwell\text{-}Boltzmann(\sigma). \end{aligned}$$

##### 2.3.1 Joint density of $(\Delta, Z)$

We will use a variable transform from the joint PDF of $(X_1, X_2)$, assuming $X_1 \sim GG(\alpha_1, \beta_1, c_1)$ and $X_2 \sim GG(\alpha_2, \beta_2, c_2)$ are independent. The joint PDF for $(X_1, X_2)$ is

$$\begin{aligned} f_{X_1X_2}(x_1, x_2|\alpha_1, \beta_1, c_1, \alpha_2, \beta_2, c_2) &= f_{X_1}(x_1|\alpha_1, \beta_1, c_1) f_{X_2}(x_2|\alpha_2, \beta_2, c_2) \\ &= \begin{cases} \frac{c_1 x_1^{c_1\alpha_1 - 1}}{\beta_1^{c_1\alpha_1}\Gamma(\alpha_1)} e^{-(\frac{x_1}{\beta_1})^{c_1}} \times \\ \frac{c_2 x_2^{c_2\alpha_2 - 1}}{\beta_2^{c_2\alpha_2}\Gamma(\alpha_2)} e^{-(\frac{x_2}{\beta_2})^{c_2}} & \text{if } x_1 \geq 0, x_2 \geq 0 \\ 0 & \text{otherwise.} \end{cases} \end{aligned}$$

The variable transformation we will use will not be going from $\mathbb{R}^2$ to $\mathbb{R}^2$, since $(\Delta, Z)$ only exists in $\{(\delta, \zeta) \in \mathbb{R}^2 | 0 \leq \zeta \leq \sqrt{2}\}$. If we have the change of coordinates $\Phi : \{(\delta, \zeta) \in \mathbb{R}^2 | 0 \leq \zeta \leq \sqrt{2}\} \to \{(x_1, x_2) \in \mathbb{R}^2 | x_1 \geq 0, x_2 \geq 0\}$, then

$$\begin{pmatrix} \Delta \\ Z \end{pmatrix} = \Phi^{-1} \begin{pmatrix} X_1 \\ X_2 \end{pmatrix} = \begin{pmatrix} X_1 - X_2 \\ \frac{\sqrt{2}|X_1 - X_2|}{X_1 + X_2} \end{pmatrix}.$$

In order to find the joint PDF (and then later the joint CDF) for $\Delta$ and $Z$, we need both $\Phi$ and its derivative $[\mathbf{D}\Phi\binom{\delta}{\zeta}]$. These will necessarily be piecewise-defined functions, since $\delta > 0$ means $x_1 > x_2$ and $\delta < 0$ means $x_1 < x_2$. We do not need to worry about $\delta = 0$ and $x_1 = x_2$, since these correspond to lines in $\mathbb{R}^2$, which have measure zero and do not contribute to any probability mass. When we invert $\Phi^{-1}$, we find

$$\begin{pmatrix} x_1 \\ x_2 \end{pmatrix} = \Phi \begin{pmatrix} \delta \\ \zeta \end{pmatrix} = \begin{cases} \frac{1}{2} \begin{pmatrix} (\frac{\sqrt{2}}{\zeta} + 1)\delta \\ (\frac{\sqrt{2}}{\zeta} - 1)\delta \end{pmatrix} & \text{if } \delta > 0 \\ \\ -\frac{1}{2} \begin{pmatrix} (\frac{\sqrt{2}}{\zeta} - 1)\delta \\ (\frac{\sqrt{2}}{\zeta} + 1)\delta \end{pmatrix} & \text{if } \delta \leq 0 \end{cases}.$$

Notice that when $\delta > 0$ and $0 < \zeta < \sqrt{2}$, we get $\Phi(\delta, \zeta) \in \{(x_1, x_2) \in \mathbb{R}^2 | x_1 > x_2,\ x_1 > 0,\ x_2 > 0\}$, and when $\delta < 0$ and $0 < \zeta < \sqrt{2}$, we get $\Phi(\delta, \zeta) \in \{(x_1, x_2) \in \mathbb{R}^2 | x_1 < x_2,\ x_1 > 0,\ x_2 > 0\}$.

For the change of variables formula, we will need $|\det([\mathbf{D}\Phi\binom{\delta}{\zeta}])|$ (Rice, 2007; Hubbard and Hubbard, 2009). If we have $\Phi\binom{\delta}{\zeta} = \binom{\Phi_1(\delta,\zeta)}{\Phi_2(\delta,\zeta)}$, then we can compute $[\mathbf{D}\Phi\binom{\delta}{\zeta}]$ as the Jacobian matrix

$$[\mathbf{D}\Phi \begin{pmatrix} \delta \\ \zeta \end{pmatrix}] = \begin{pmatrix} \frac{\partial \Phi_1(\delta,\zeta)}{\partial \delta} & \frac{\partial \Phi_1(\delta,\zeta)}{\partial \zeta} \\ \frac{\partial \Phi_2(\delta,\zeta)}{\partial \delta} & \frac{\partial \Phi_2(\delta,\zeta)}{\partial \zeta} \end{pmatrix}.$$

We find

$$|\det([\mathbf{D}\Phi\begin{pmatrix}\delta\\ \zeta\end{pmatrix}])| = \begin{cases} |-\frac{\sqrt{2}\delta}{2cv^2}| & = & \frac{\sqrt{2}\delta}{2cv^2} & \text{if } \delta > 0 \\ |\frac{\sqrt{2}\delta}{2cv^2}| & = & -\frac{\sqrt{2}\delta}{2cv^2} & \text{if } \delta \le 0 \end{cases}.$$

Then the joint PDF for $\Delta$ and $Z$, using this $2 \times 2$ transform and the change of variables formula (presented in Hubbard and Hubbard (2009) as Theorem 4.10.12 and Proposition A in Section 3.6.2 of Rice (2009)), is

$$\begin{aligned} f_{\Delta,Z}(\delta,\zeta|\alpha_1,\beta_1,c_1,\alpha_2,\beta_2,c_2) &= f_{X_1X_2}(\Phi\tbinom{\delta}{\zeta}|\alpha_1,\beta_1,c_1,\alpha_2,\beta_2,c_2)|\det([\mathbf{D}\Phi\tbinom{\delta}{\zeta}])| \\ &= \begin{cases} \frac{c_1}{\beta_1^{c_1\alpha_1}\Gamma(\alpha_1)}((\frac{\sqrt{2}}{\zeta}+1)\frac{\delta}{2})^{c_1\alpha_1-1}\exp(-((\frac{\sqrt{2}}{\zeta}+1)\frac{\delta}{2\beta_1})^{c_1})\times \\ \frac{c_2}{\beta_2^{c_2\alpha_2}\Gamma(\alpha_2)}((\frac{\sqrt{2}}{\zeta}-1)\frac{\delta}{2})^{c_2\alpha_2-1}\exp(-((\frac{\sqrt{2}}{\zeta}-1)\frac{\delta}{2\beta_2})^{c_2})\times \\ \frac{\sqrt{2}\delta}{2cv^2} \\ \text{if } \delta \ge 0, 0 \le \zeta \le \sqrt{2} \\ \\ \frac{c_1}{\beta_1^{c_1\alpha_1}\Gamma(\alpha_1)}((-\frac{\sqrt{2}}{\zeta}+1)\frac{\delta}{2})^{c_1\alpha_1-1}\exp(-((-\frac{\sqrt{2}}{\zeta}+1)\frac{\delta}{2\beta_1})^{c_1})\times \\ \frac{c_2}{\beta_2^{c_2\alpha_2}\Gamma(\alpha_2)}((-\frac{\sqrt{2}}{\zeta}-1)\frac{\delta}{2})^{c_2\alpha_2-1}\exp(-((-\frac{\sqrt{2}}{\zeta}-1)\frac{\delta}{2\beta_2})^{c_2})\times \\ (-\frac{\sqrt{2}\delta}{2cv^2}) \\ \text{if } \delta < 0, 0 \le \zeta \le \sqrt{2} \\ \\ 0 \\ \text{otherwise.} \end{cases} \end{aligned}$$

### Supplementary Figures

Supplementary figures referenced in the main paper are included below.

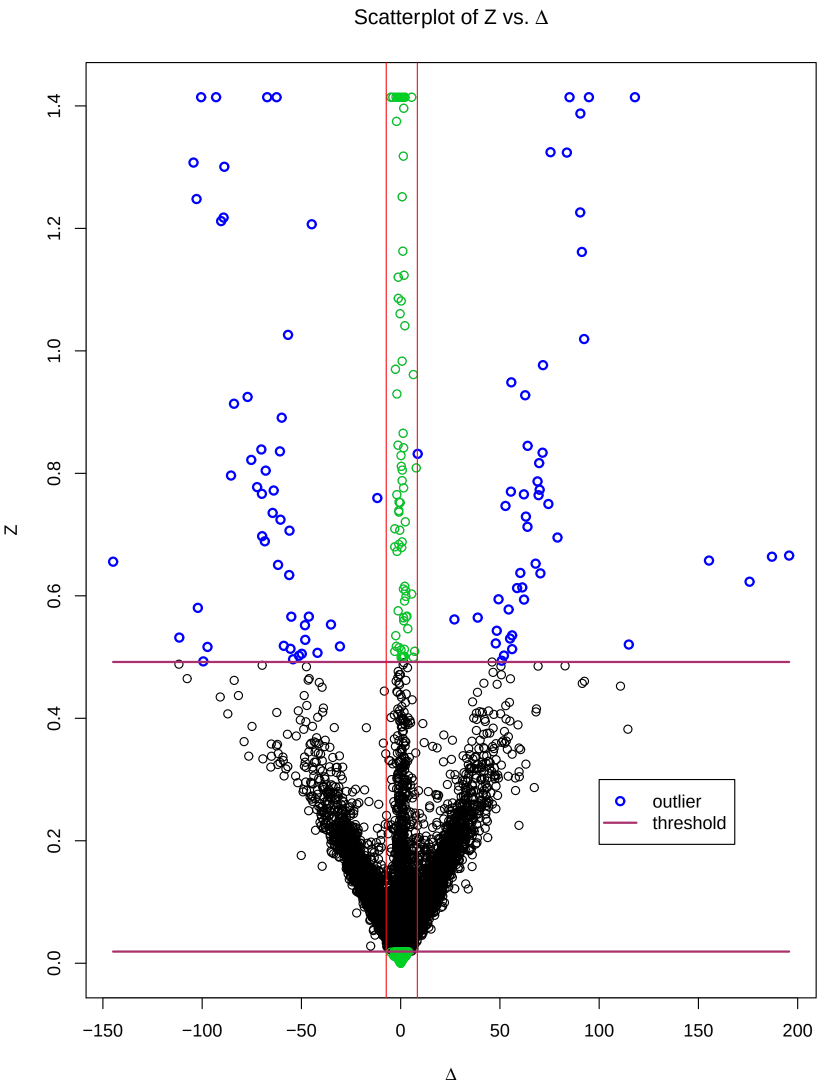


**Supplementary Figure 1:**

Relative activity of 181 commercially available kinase inhibitors against a panel of 292 recombinant protein kinases where the activity for each kinase-inhibitor pair measured in duplicate (Anastassiadis et al., 2011; 2013). A scatterplot of the coefficient of variation $Z$ versus the difference $\Delta$ for all pairs of replicates, resembling a "volcano plot", can be used to assess reproducibility of the data. The red vertical lines represent $k = 1$ standard deviation (of $\Delta$) away from $\hat{\mu}+\hat{\theta}$ and green circles within this band represent replicates exhibiting large $Z$ but with small $\Delta$. The maroon horizontal lines represent the outlier determination thresholds for $Z$ and replicates whose $Z$ is above the level indicated by top horizontal line or below the level indicted by the bottom horizontal line *and* $\Delta$ is outside this band of red vertical lines are considered to be outliers (shown as blue circles). Replicates shown as green and black circles are not considered to be outliers.

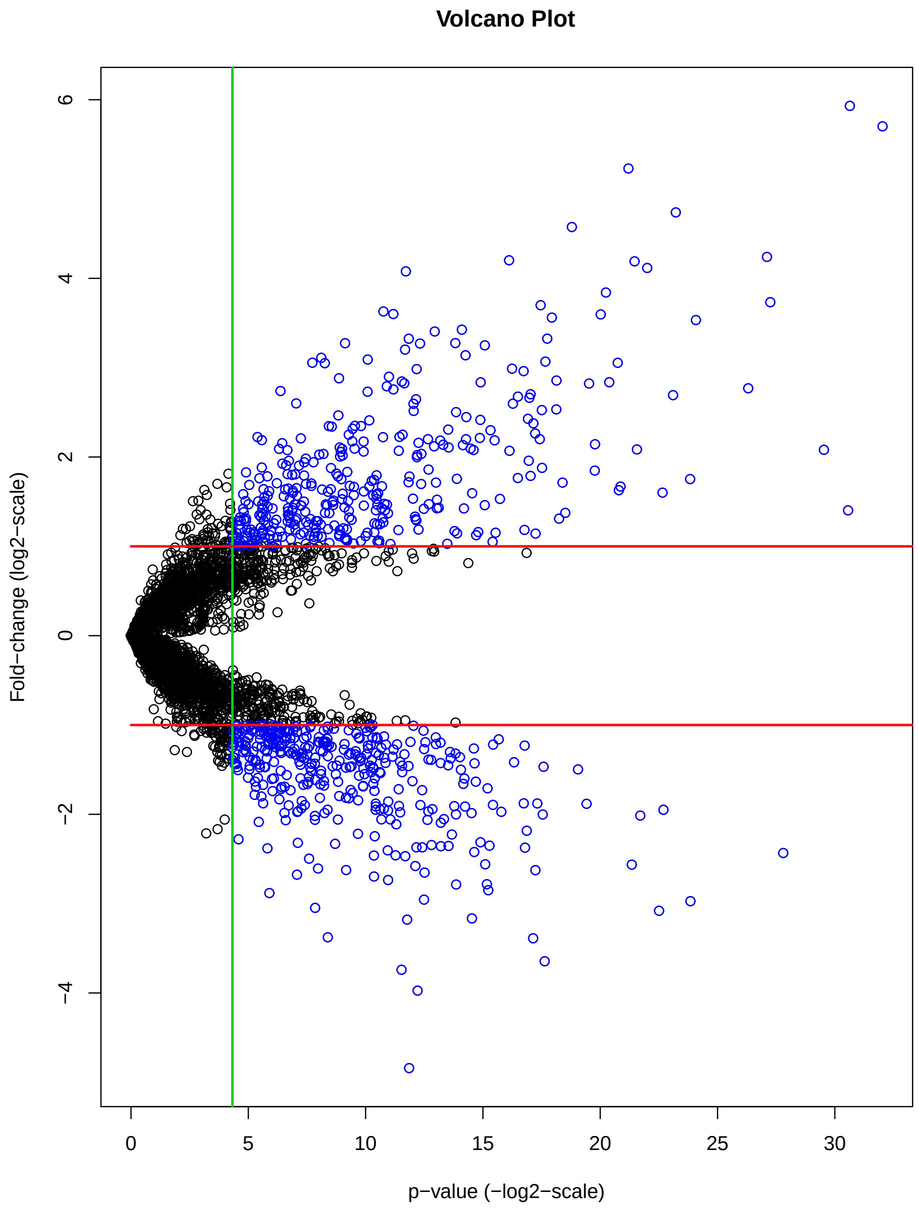


**Supplementary Figure 2:**

Example of a traditional "volcano plot" from gene expression data. The green vertical line represents the threshold for determining statistical significance ($p \leq 0.05$, shown on the negative $\log_2$-scale) and the red horizontal lines represent the threshold for determining biological significance (two-fold change in expression, shown on the $\log_2$-scale). Each point represents a genomic feature of interest in the study. Features that appear to the right of the green vertical line *and* above the top horizontal line or below the bottom horizontal line (shown as blue circles) are typically considered to be both statistically and biologically significant and deemed worthy of further investigation.

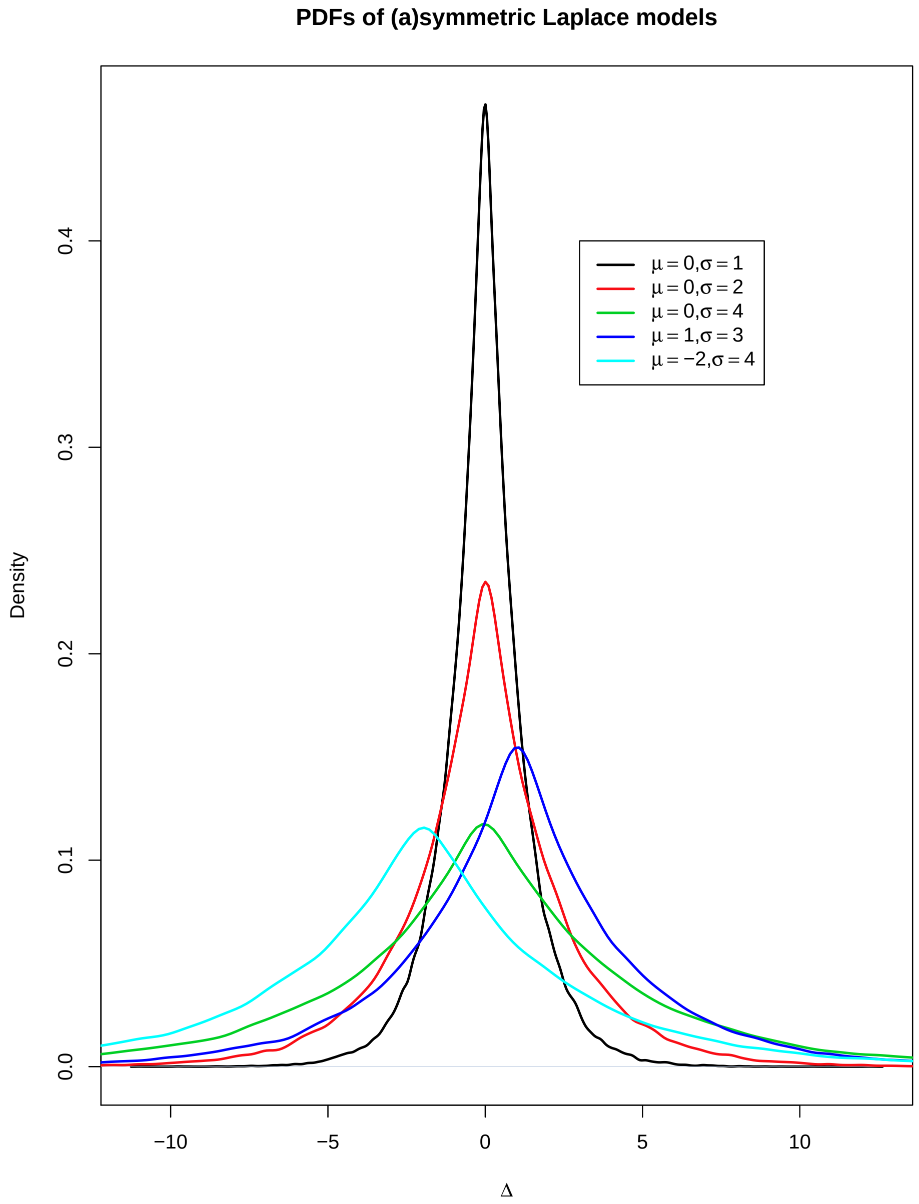


**Supplementary Figure 3:**

PDFs of symmetric and asymmetric Laplace models for different values of the parameters $\mu$ and $\sigma$ in equation (2.1).

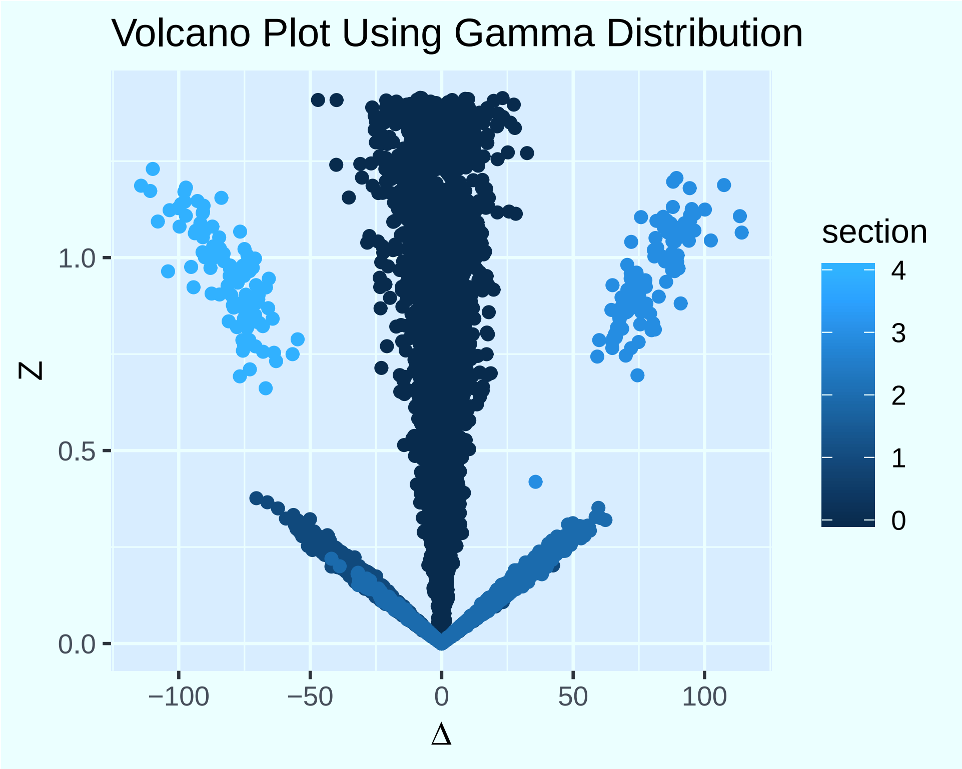


**Supplementary Figure 4:**

$(\Delta, Z)$-plots corresponding to data generated from the gamma model. We colored different sections to emphasize the similarities to volcano plots presented in Supplementary Figures 1 & 2. We created all figures using the `R` package `ggplot2` (Wickham, 2016).

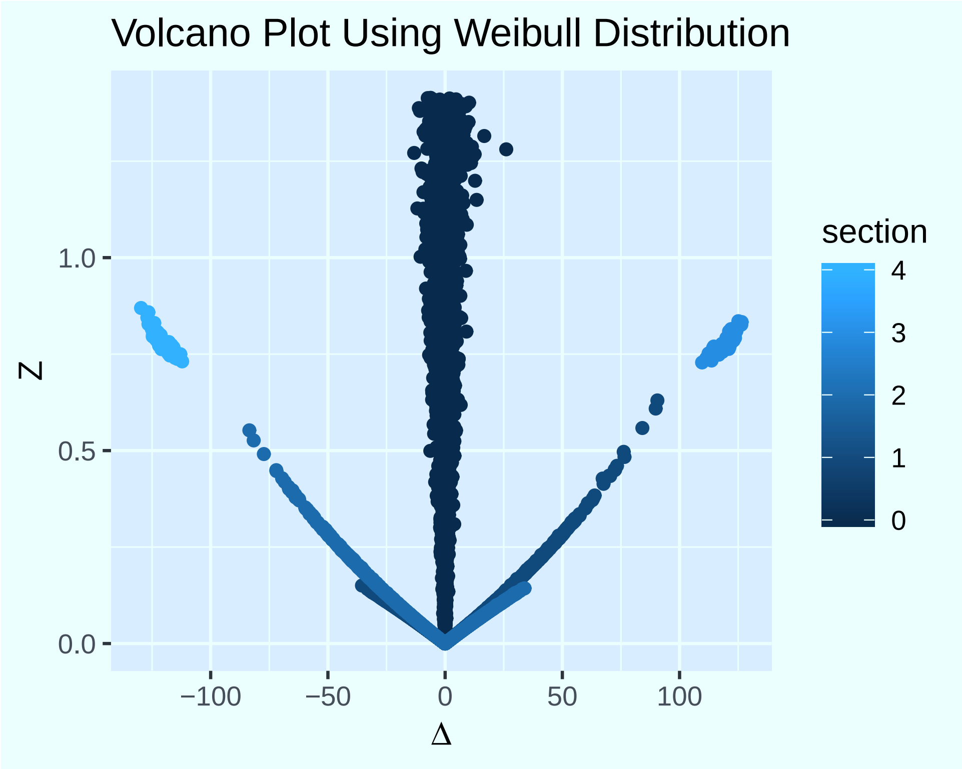


**Supplementary Figure 5:**

$(\Delta, Z)$-plots corresponding to data generated from the Weibull model using the `R` package `ggplot2` (Wickham, 2016).

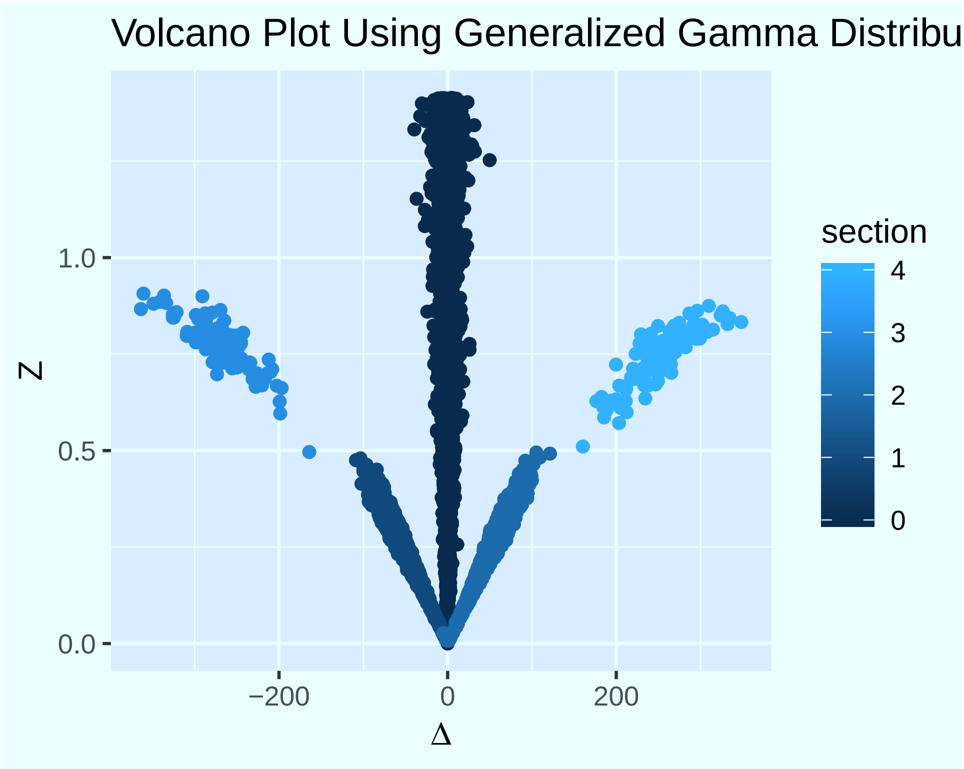


**Supplementary Figure 6:**

$(\Delta, Z)$-plots corresponding to data generated from the generalized gamma (GG) model. We colored different sections to emphasize the similarities to volcano plots presented in Supplementary Figures 1 & 2. We created all figures using the `R` package `ggplot2` (Wickham, 2016).

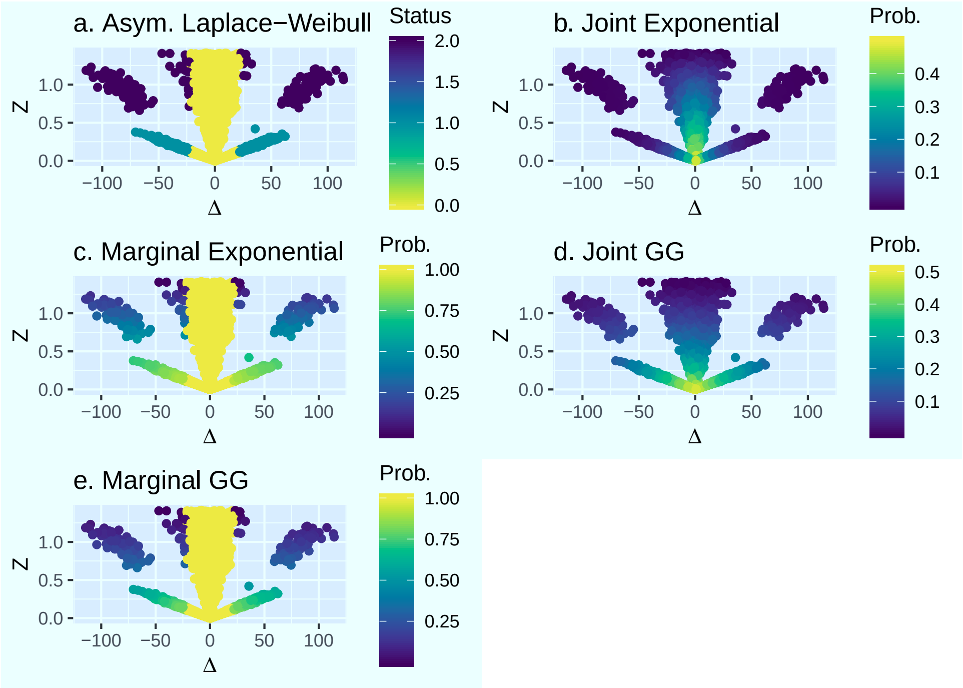


**Supplementary Figure 7:**

$(\Delta, Z)$-plots of data generated from the gamma model, labeled by the method used to identify outliers. We colored the asymmetric Laplace-Weibull plot by outlier status and other plots by the calculated probability $q(\Delta, Z)$. All plots were made using the `R` package `ggplot2` and arranged using the package `gridExtra` (Wickham, 2016; Auguie, 2017). The color scheme is based on the `R` package `viridis` (Garnier, 2018).

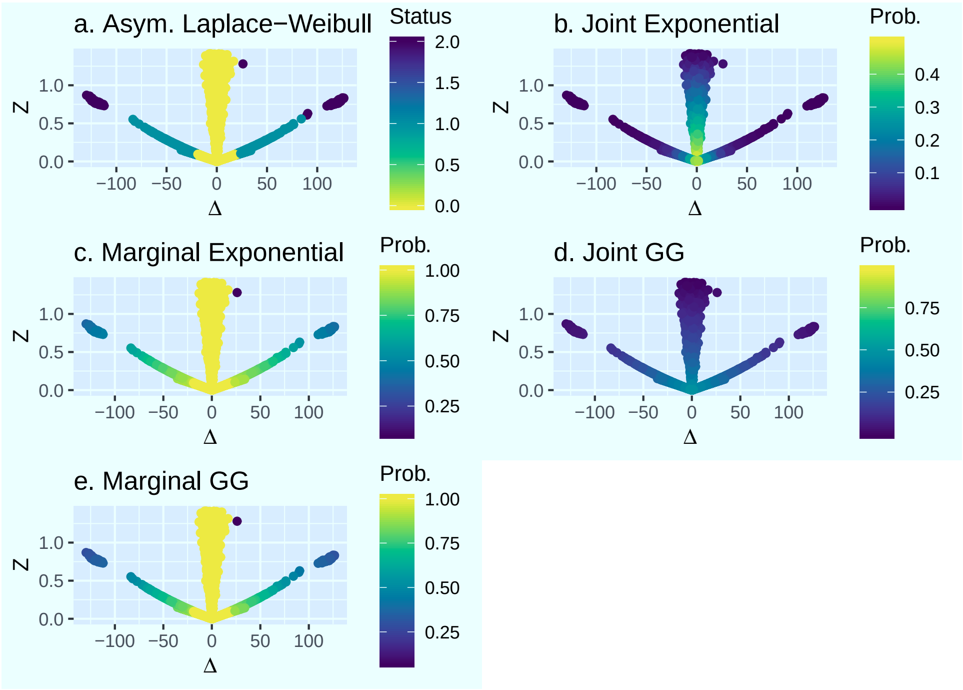


**Supplementary Figure 8:**

$(\Delta, Z)$-plots of data generated from the Weibull model, labeled by the method used to identify outliers. We colored the plots using the same approach and `R` packages as in Supplementary Figure 7.

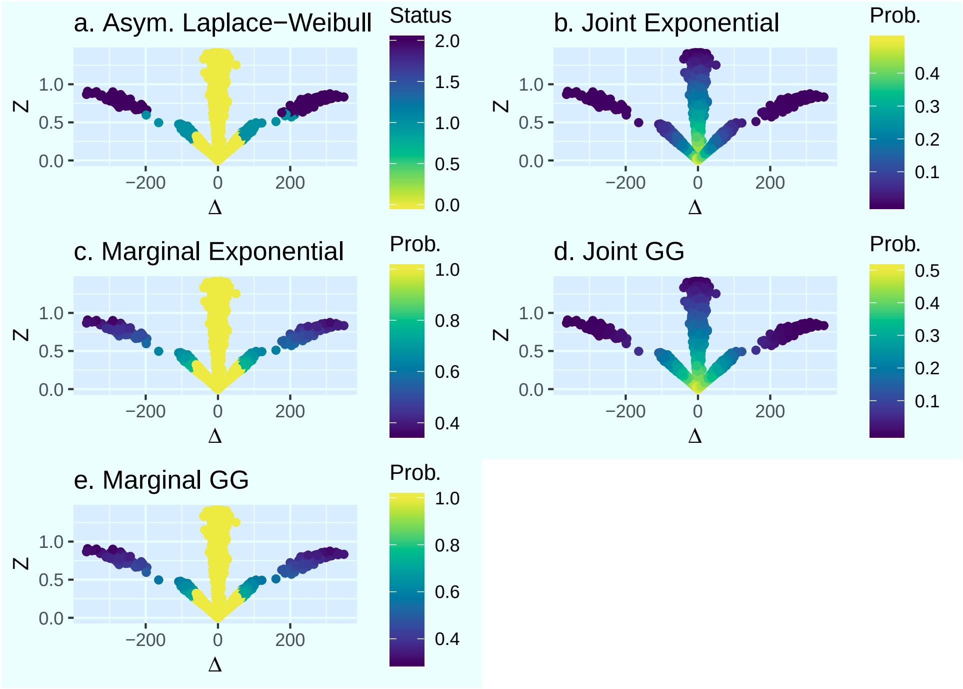


**Supplementary Figure 9:**

$(\Delta, Z)$-plots of data generated from the GG model, labeled by the method used to identify outliers. We colored the plots using the same approach and `R` packages as in Supplementary Figure 7.

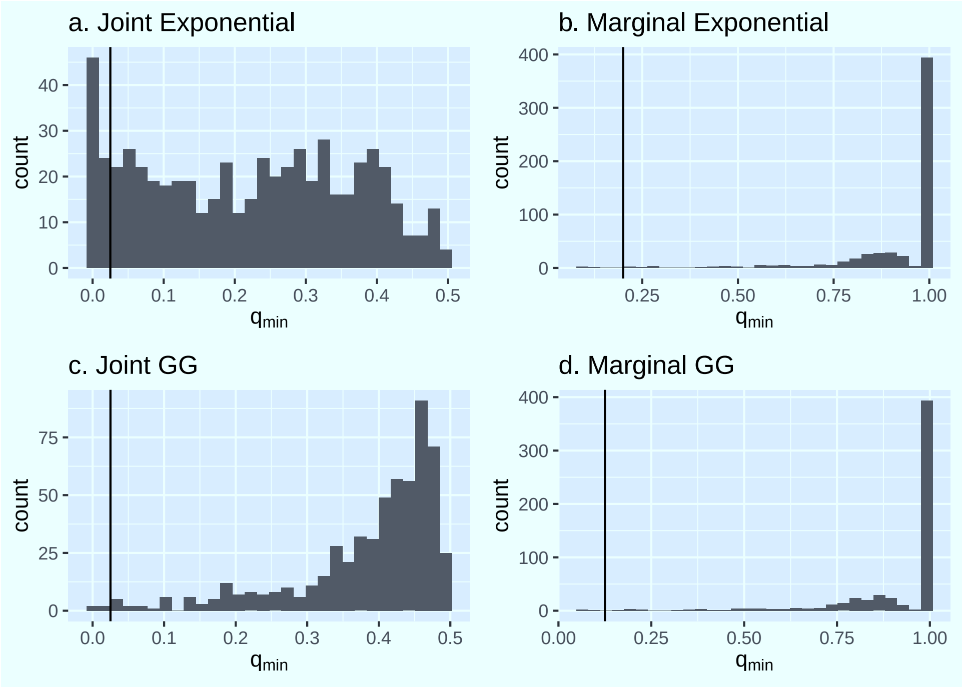


**Supplementary Figure 10:**

For the proteomics data, we calculated outlier probabilities for each of the three pairs from triplicate measurements, using the joint exponential, marginal exponential, joint GG, and marginal GG methods. We then plotted the empirical distributions of the minimum $q$-value for each measurement trio. We plotted lines at $q_{\text{min}} = 0.025$ on the joint exponential plot, $q_{\text{min}} = 0.2$ on the marginal exponential plot, $q_{\text{min}} = 0.025$ on the joint GG plot, and $q_{min} = 0.125$ on the marginal GG plot. We created these plots using the `R` package `ggplot2` and arranged them using `gridExtra` (Wickham, 2016; Auguie, 2017).
